# Exploring unique sequence repeat patterns and secondary structures in rDNA internal transcribed spacers ITS1 and ITS2 for characterization of catfish species

**DOI:** 10.1101/2024.01.10.575031

**Authors:** Mohd Imran, Sana Nafees

## Abstract

The genetic variabilities in internal transcribed spacers ITS1 and ITS2 of rDNA are explored to a limited extent in vertebrates including fish species. This study explores unique molecular signature in these spacers from multiple fish species from the order siluriformes. Supporting the concerted evolution, both ITS1 and ITS2 were found prevalent with microsatellite and random repeats, with diverse distribution in each species in term of nucleotide composition, position and length, giving a species-specific impression in repeats. Their secondary structures too have shown distinct structural homology with variations in helix length, loops, and non-canonical bps, where ITS2 4-helix carries additional nine conserved motifs while that of ITS1 was found highly branched with multiple sub-branching unique to each species and a long tail helix common in all. This combination of secondary structures and repeats have formed unique molecular signatures distinguishing each species. Since the evolutionary and maturation biology of rDNA is similar in vertebrates, these molecular signatures in internal spacers could effectively contribute in the molecular characterization of other species too.

## INTRODUCTION

The primary structure of DNA, i.e. linear nucleotide sequence often reveals some striking features when investigated at their secondary structure level. Not only the coding segment but the non-coding DNA especially rDNA also keeps a highly complex structural composition when it comes to the primary and secondary structures of its internal spacers-Internal Transcribed Spacer region 1 (ITS1) and region 2 (ITS2). These ITS1 and ITS2 are located in between 17-18s rDNA and 5.8s rDNA, 5.8s rDNA and 25-28s rDNA, respectively; and hence share all its general features viz. multi-chromosomal origin, tandemly repeated presence (Reed et al. 2000) and concerted evolution (Hillis and Dixon 1991). The peculiar property associated with both the internal spacers is their variable size, which varies not just in between different species but also within the same species (Harris et al. 2000; Kuriiwa et al. 2007). They show this dynamicity in their size due to the prevalence of multiple microsatellites in their sequences. Besides, the multi-chromosomal location of NORs (Nucleolus Organizer Regions) is also suggested a probable factor behind (Ritland et al. 1993). These microsatellites are tandemly repeated sequences of short nucleotide sequence/motif ranges from 1 to 6 bp (Kelkar et al. 2010). Depending on the composition, they could be mononucleotide repeats, di-, tri-, tetra-, penta-, and hexanucleotide repeats. They are basically known for their unique feature, i.e. the variation in the number of tandem repeats, their number may vary between different species or even among individual of the same species. Because of all these associated variables, the ITS regions are found with wide differences in length among different organisms. For example, in loach, the ITS1 was reported just 343 bp while it was reported 1095 bp long in humans (Gonzalez et al. 1990). Similarly, such a wide difference is also observed in ITS2 which is reportedly just 363-369 bp long in *Anopheles nuneztovari* (fritz et al. 1994), 384-490 bp in catfishes (Jansen et al. 2006) and reaches to a length of 1300 bp in Crayfish (Harris et al. 2000).

Although both the ITS1 and ITS2 are non-coding markers, they still play a crucial role in the maturation of the precursor rRNA molecule, where they provide spatial secondary structure geometry crucial for the binding of snoRNA and RAC protein complex necessary for proper ribosomal maturation. These internal spacers carry margin to bear changes in their nucleotide composition and sequence length to the point of non-disruption of their secondary structure which appears as an evolutionary adaptation to conserve their functional aspect (Pleyte et al. 1992; Cote and Peculis 2001; Abeyrathne et al. 2002). The ITS2 also provides better information for taxonomic inferences when used along with its secondary structure (Coleman 2003). The same approach isused in this study for both the internal spacers ITS1 and ITS2.

The presence of extremely conserved rDNA segments at their 5’ and 3’ end of ITS1 and ITS2 makes them an easy candidate for PCR based study. They have been studied extensively for barcoding fungal species (Schoch et al. 2012; Kapoor et al. 2018), resolving biogeography (16) and species complex in plants (Ritland et al. 1993; Vander et al. 1998). Even within the animal kingdom they are explored more in the invertebrate, viz. phylogenetic relationship in Ciliates (Chen et al. 2021), corals (Aguilar and Reimer 2010), resolving cryptic speciation in nematodes (Pinacho-Pinacho et al. 2018), phylogeography of Mosquito (Fritz et al. 1994), species identification of Molluscs (Hurtado et al. 2011). Several studies have also been conducted in vertebrates for phylogenetic relationship and species identification (Voronov et al. 2006; Coleman 2013) including several reports from fish species as well (Booton et al. 1997; Domanica et al. 1997; Sajdak and Phillips 1997; Presa et al. 2002; Montoya-Burgos 2003; Vinas and Tudela 2009; Vera et al. 2011; Myslowski et al. 2013; Puncher et al. 2015) but still it is not widely explored in any vertebrate group even in pisces.

Since the structural organization of rDNA is conserved across the eukaryotes and the complex process of ribosomal maturations too, is similar (Greelings et al. 2000; Abeyrathne et al. 2002), five fish species from the order silurifomes, class pisces of the vertebrate, were included in the study to trace the distribution of repeated sequences in the internal transcribed spacer regions ITS1 & ITS2 of rDNA, to predict their secondary structures, and to look for their combined potential in species diagnosis. The catfish species included were *Clarias batrachus* and *C. gariepinus* (family: Clariidae); *Heteropneustes fossilis* (family: Heteropneustidae); *Sperata seenghala* (family: Bagridae), and *Wallago attu* (family: Siluridae). Additionally, no prior study has explored the rDNA spacer structures in vertebrate species in this perspective.

## MATERIALS AND METHODS

### Sample and DNA Isolation

Multiple specimens of the fish species were collected from three distant locations on the northern riverine system of Ganges i.e., Garhwal (30°24’5.35“N, 78°45’46.11“E), Aligarh (27°58’41.57“N, 78° 8’17.96“E) and Varanasi (25°19′08″N 83°00′46″E). Specimens were identified using standard identification features (Talwar and Jhingran 1992). Fishes were anesthetized prior to blood extraction with MS222 anesthetic. Approximately 0.5 ml of the blood extracted from the heart and caudal vein in EDTA coated vial. Extraction of total genomic content, including both nuclear and mitochondrial genome, was performed according to the high salt method (Montgomery and Sise 1990).

### Primer Designing

To deal with the questions under study, ITS1 and ITS2 were targeted separately. Since, no such primers were available that could specifically target these spacer regions in the aforesaid species, new sets of primers were designed from complete rDNA cistron of *Clarias gariepinus* (Genbank accession no. AJ876383.1) using NCBI primer designing tool (https://www.ncbi.nlm.nih.gov/tools/primer-blast/) (Ye et al. 2012) which include ITS1 forward (ITS.1.F1-TCGATCAAACTTGACTATCTAGAGG; ITS.1.F2-CGATCAAACTTGACTATCTAGAGG A); ITS1 reverse (ITS.1.R1-CAGCTGGCTGCGTTCTTCAT); ITS2 forward (ITS2.F-CAGGACACATTGATCATCGAC AC); ITS2 reverse (ITS2.R-AATTCAGCGGGTCGTCTC). The primers were found compatible with species from 159 genera, where 37 genera were found with compatibility in multiple sister species (Supplementary data S1).

### PCR standardization, Gel electrophoresis and Sequencing

The PCR conditions were standardized for all five species using MgCl_2_ range of 1-3mM, annealing temperature (Ta) range of 46-56°C for 45-60 seconds depending upon the ITS marker. The amplification was done using peqSTAR 2X thermocycler (PEQLAB Biotechnologie, Germany) in 15µl PCR reaction mixture containing 50-100ng of DNA template, 1.5µl of 10X PCR reaction buffer, 1.5 to 3.5mM MgCl_2_, 0.3µl of 10mM dNTPs mix (fermentas), 0.25µl each of 10pm forward and reverse primers, 0.25 unit of TaqDNA polymerase (fermentas). The PCR cycle involved an initial denaturation at 95°C for 5 min, followed by 35 cycles as (i) template denaturation at 95°C for 1min, (ii) primer annealing varied with species within a range of 55-62°C, and (iii) extension at 72°C for 1 min. A final extension was also performed at 72°C for 5 minutes at the end of 35 cycles.

The PCR products were subjected to electrophoresis on 1.5% agarose gel. Both the set of primers provided a very specific noise free amplification of ITS1 and ITS2 in all five species, with a clear indication of reproducible results as shown by the identical band positions among the multiple individuals of each species. The difference in the band positions in different species for both the spacers (ITS1 and ITS2) unquestionably depict that neither of them possesses size uniformity and thus have shown considerable size range (Fig. 1). Sanger sequencing was performed using ABI 3730XL DNA Analyser with BigDyeTM Terminator v3.1 Cycle Sequencing Kit (Thermo Fisher Scientific), and the sequences were submitted to NCBI GenBank.

**Figure 1.**
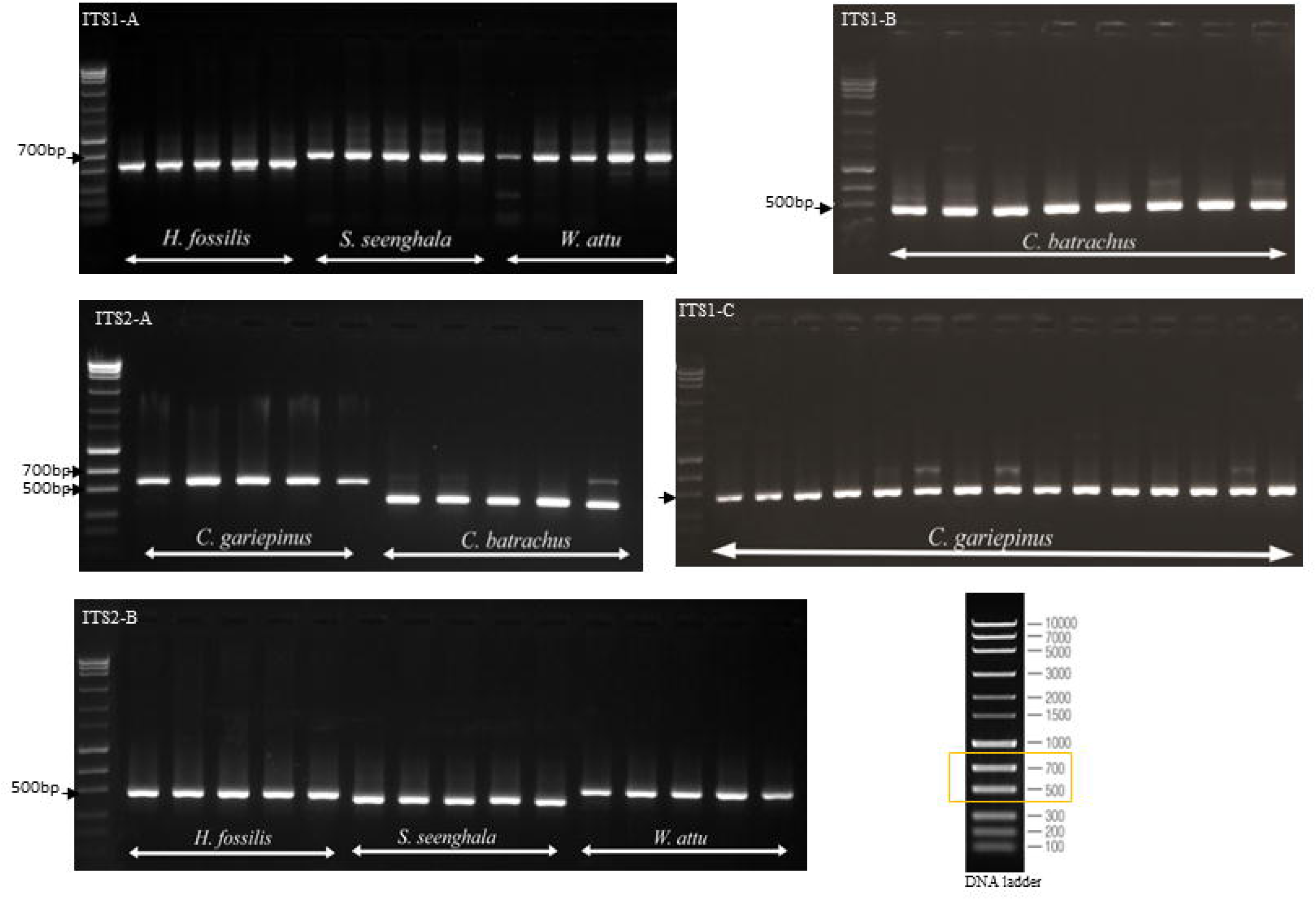
Agarosegel images of ITSI andITS2 amplicons. The → indicate the amplicon size of ITSI and ITS2 against the standardDNA ladder in the range of 500-700bp.

### Sequence Analysis

The ITS1 and ITS2 sequences from multiple replicates of all five species were aligned with Clustal W, and required editing viz. labelling was done in ‘Mega7: Text File Editor and Format Converter’ (Kumar et al. 2016). The repeated sequence tracing was done with MISA-a web based repeat finding tool (https://webblast.ipk-gatersleben.de/misa/) (Beier et al. 2017) and with manual tracing of sequences. The aligned sequences were then compared and analyzed within and between species for their relative differences, to find the overall pattern of repeats and unique microsatellites if any from the point of view of species identity.

### Secondary Structure Analysis

For ITS2, the secondary structure analysis was done at ITS2 database. This includes annotation of ITS2 sequences, secondary structure prediction using homology modelling, multiple alignment of secondary structures, conserved motif of secondary structures (Koetschan et al. 2010). The ITS1 secondary structure prediction was done using RNAfold web server with minimum free energy method (http://rna.tbi.univie.ac.at/cgi-bin/RNAWebSuite/RNAfold.cgi) (Gruber et al. 2008). The LocARNA web server (https://rna.informatik.uni-freiburg.de) (Will et al. 2007, 2012; Raden et al. 2018) was used for the multiple sequence alignment of ITS1 sequences on the basis of their secondary structure. The annotation of both ITS1 and ITS2 secondary structures was done in a similar manner using software-4SALE (Seibel et al. 2006, 2008) and VARNA (Darty et al. 2009) (Supplementary data S2).

## RESULTS

### Description of ITS Repeats

The ITS1 and ITS2 sequences of each catfish species were found with the presence of multiple repeats. Some repeats qualify the standard definition of microsatellites (Oliveira et al. 2006) while many don’t fit into that but effectively describe the repetitive nature of both ITS1 and ITS2 and placed in the category of the random repeats. The highly variable distribution of distinct microsatellites and random repeats are found affecting the lengths of both ITS1 and ITS2 in a considerable manner.

### Microsatellite and Random repeats

The repeat distributions and patterns are found highly diverse in five species ranging from short to long; simple to complex. Repeats of every kind from mononucleotide to tetranucleotide; perfect, imperfect, interrupted and composite (Table 1 & 2, Supplementary data S3). The ITS1 and ITS2 both have shown microsatellite and random repeats except the *H. fossilis* ITS2 which consists of only random repeats. Not all five species were found prevalent with every type of repeat. A considerable degree of intraspecific similarity was observed in repeats, where each species was found containing its own set of repeats differing from that of other species in terms of nucleotide composition, position (locus) and length. From the secondary structural point of view, this distinct composition and distribution of repeats causes wide differences between species in terms of their location on the different helices of secondary structure, and thus providing a species specific architecture to secondary structures.

**Table 1.**
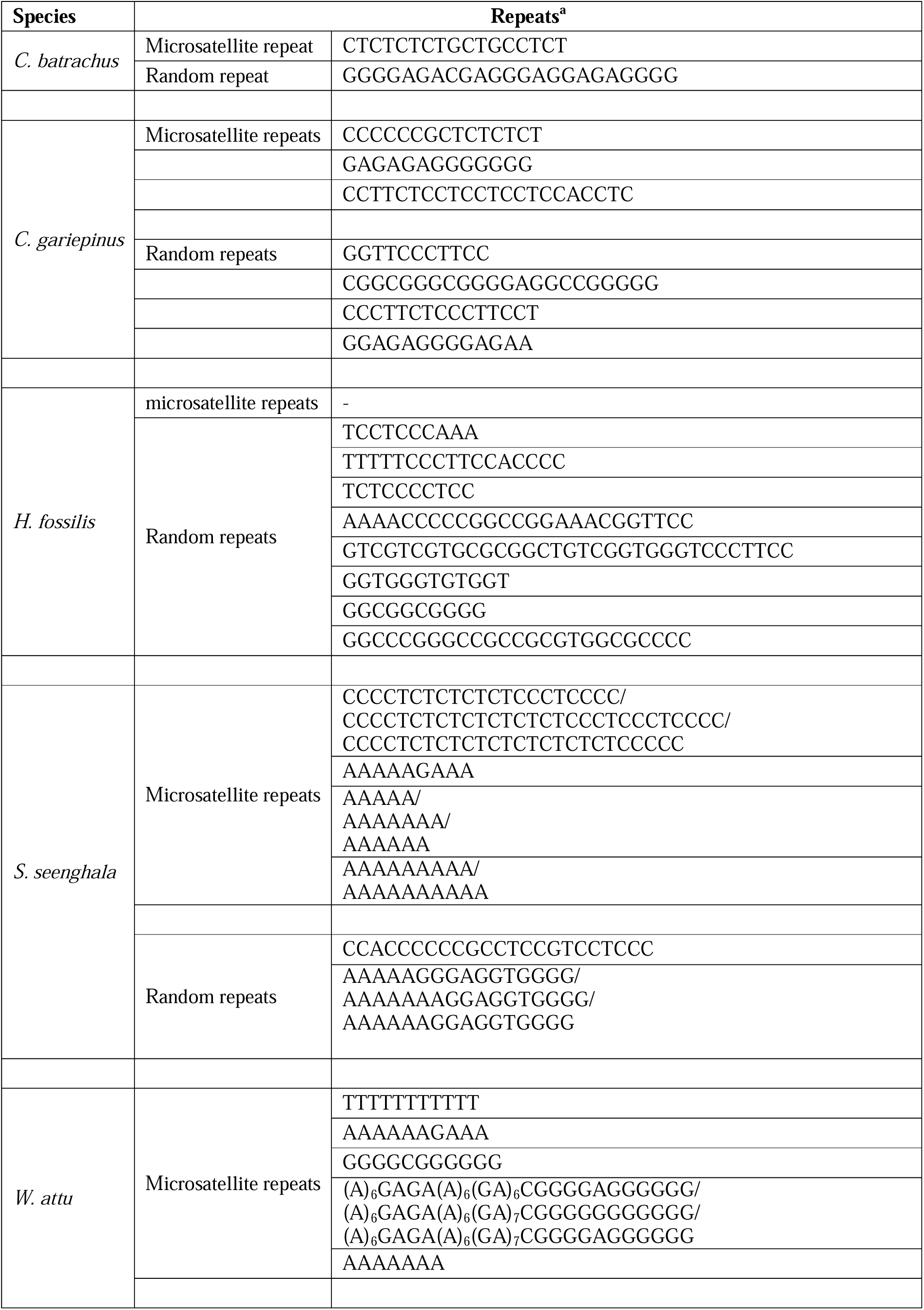
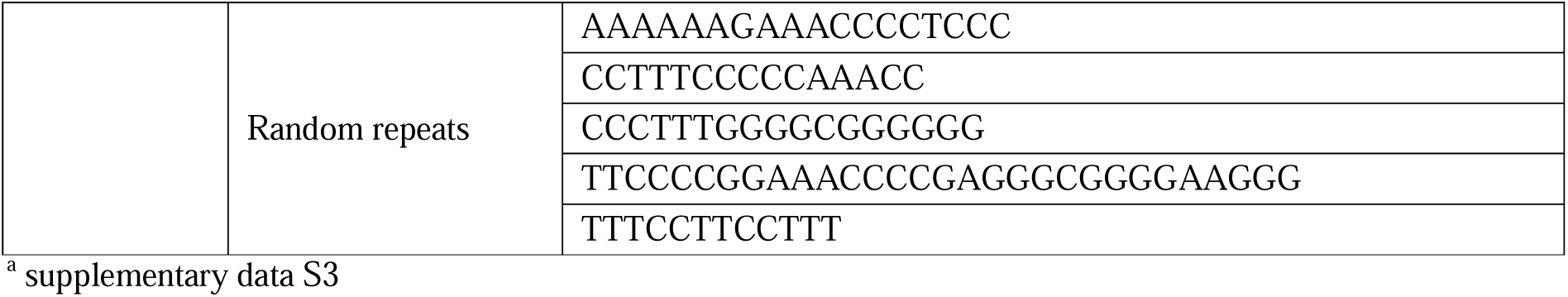
ITS2 nucleotide repeats. Table showing the microsatellite and random repeats in the ITS2 of catfish *C. batrachus, C. gareipinus, H. fossilis, Sperata seenghala* and *W. attu*. The variants of the same microsatellite between different individuals of the same species are separated by (/).

**Table 2.**
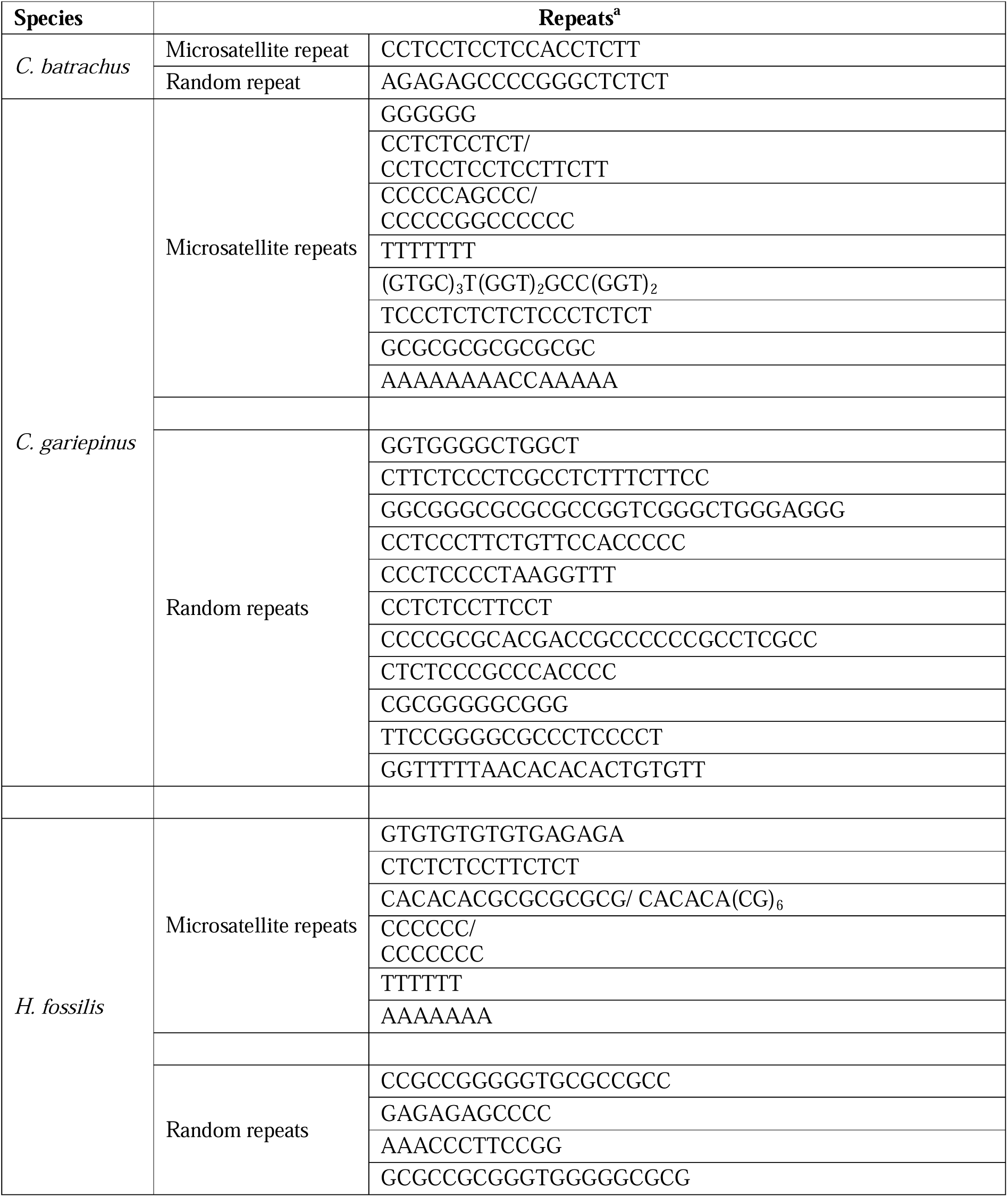
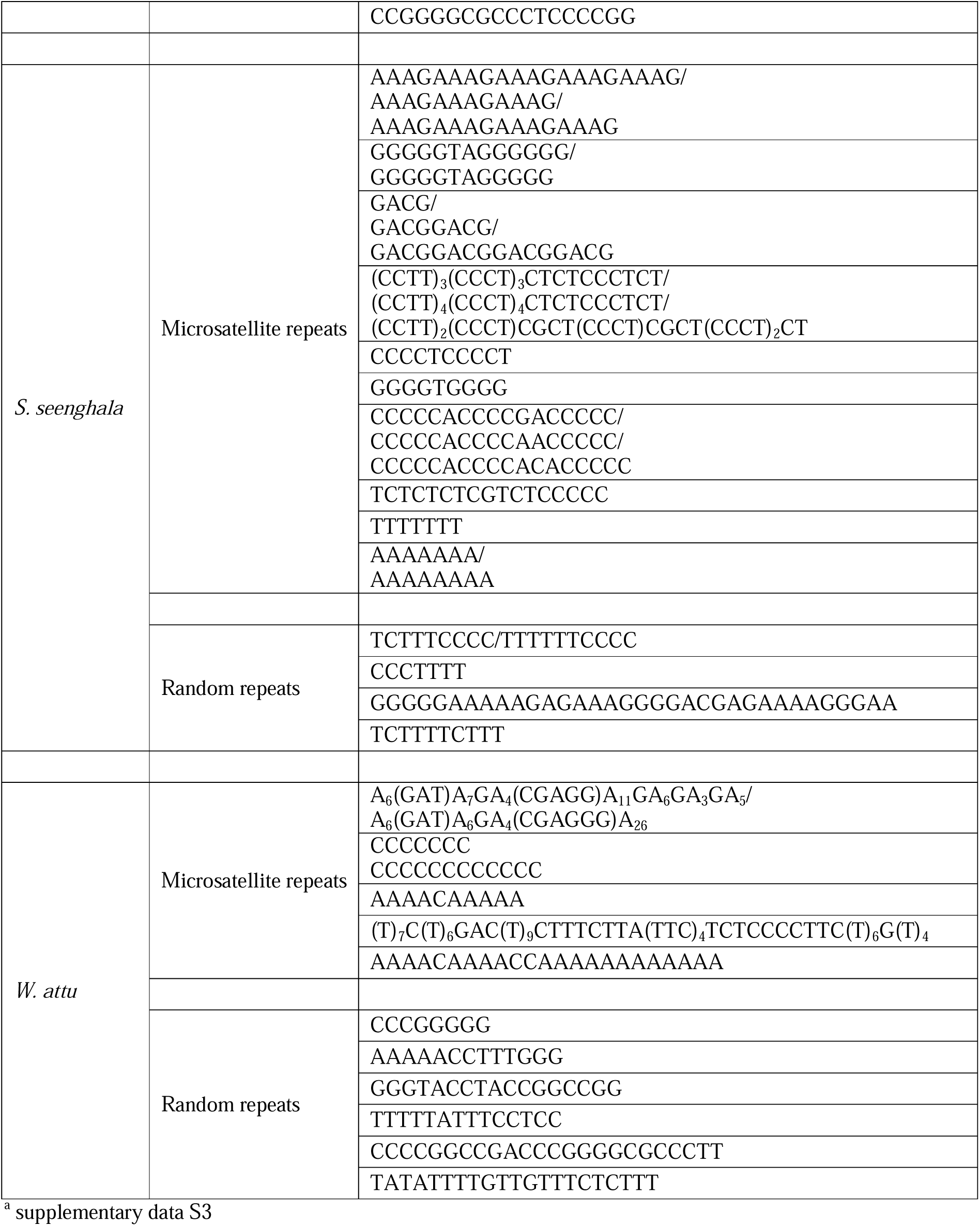
ITS1 nucleotide repeats. Table showing the microsatellite and random repeats in the ITS1 of catfish *C. batrachus, C. gareipinus, H. fossilis, Sperata seenghala* and *W. attu*. The variants of the same microsatellite between different individuals of the same species are separated by (/).

### Secondary Structure of ITS2

#### Structure prediction

All the species were found possessing similar architecture of secondary structure with all consist of 4 helices branching off from the central ring which is one of the hallmarks of eukaryotic ITS2 structure with helix III is the longest among the four. The central ring is not usually a perfect circular ring rather quite variable in shape which continues into a stalk through which the ITS2 structure remains attached to the parent rRNA gene transcript. Helix I, II, IV and ring stalk are closely situated at one side of the central ring while the helix III is oriented in a direction opposite to the rest (Fig. 2 & 3, Supplementary data S4).

**Figure 2.**
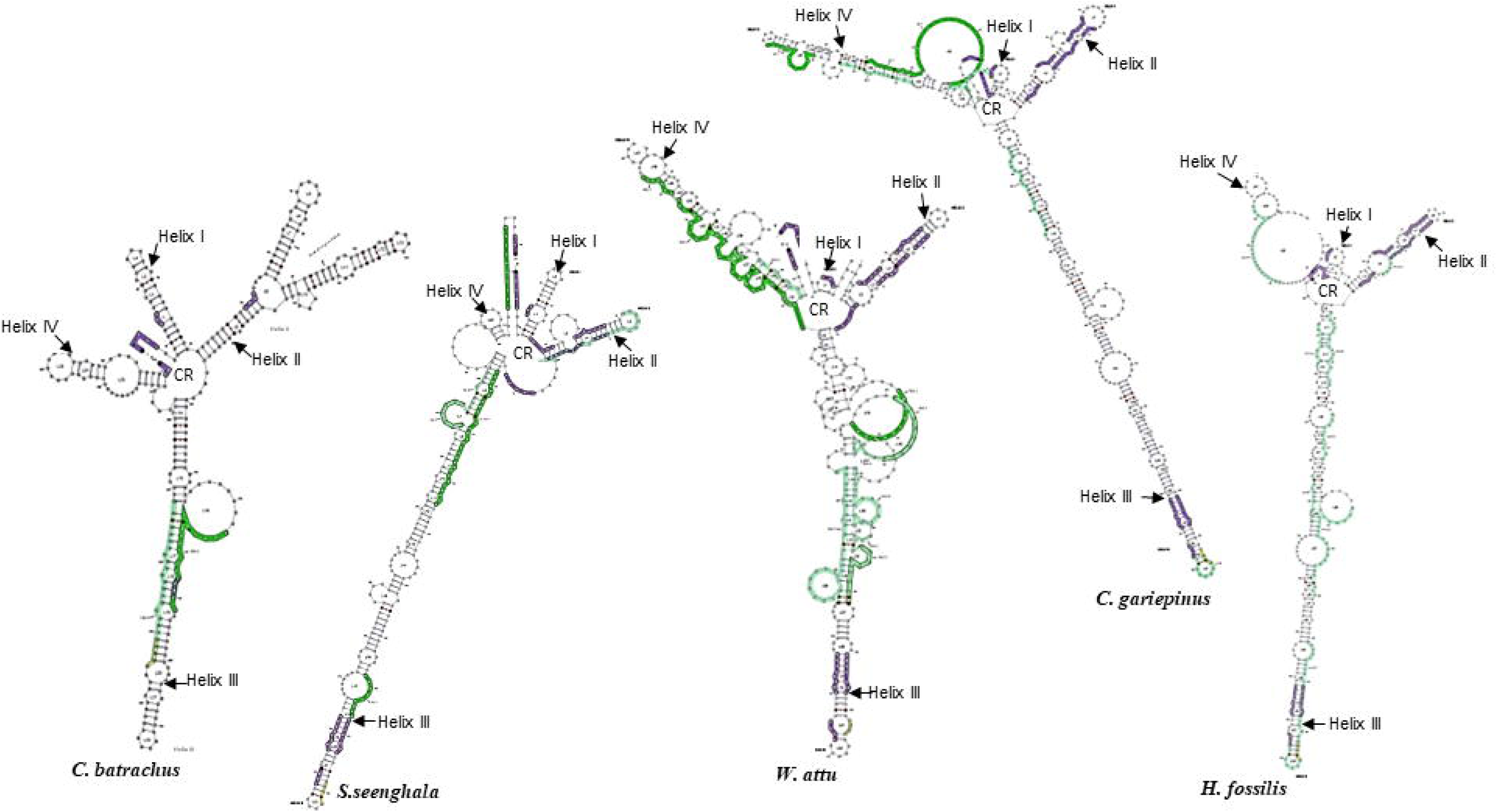
Secondary structureof ITS2. Thefigure showing the 4-helix secondary structure of ITS2 in *C. batrachus, S. seenghala, W. attu, C. gariepinus and H. fossilis*., vith a clockwise arrangemet of helix I to IV rowida central ring (CR).

#### Motif search

The eukaryotic ITS2 possess some specific nucleotide features known as hallmark of the eukaryotic ITS2 sequences and its secondary structure as they are observed ubiquitously in different eukaryotic species. The motif search using ITS2 database server has found these hallmark features in the ITS2 sequences of all five catfish species, too, which include – i) A conserved UG motif in different variants of UGGU/UGGGU/UGGGG string located in the helix III of the secondary structure near its terminal loop (Fig. 4). ii) Pyrimidine-Pyrimidine mismatch in the helix II of the ITS2 secondary structure which consists of placement of a pyrimidine base against a pyrimidine base instead of a purine base. It is found to be cytosine-uracil in *C. batrachus, C. gareipinus, H. fossilis, W. attu*, and cytosine-cytosine in *S. seenghala*. It leads to a slight bulge in helix II of the secondary structure because of the absence of any bonding between the bases of the same group (Fig. 3). The exact position of these motifs varies from species to species depending upon the length of the ITS2 which ranges over a considerable size as previously described. The deviations from these hallmarks of secondary structure are also reported (48).

**Figure 3:**
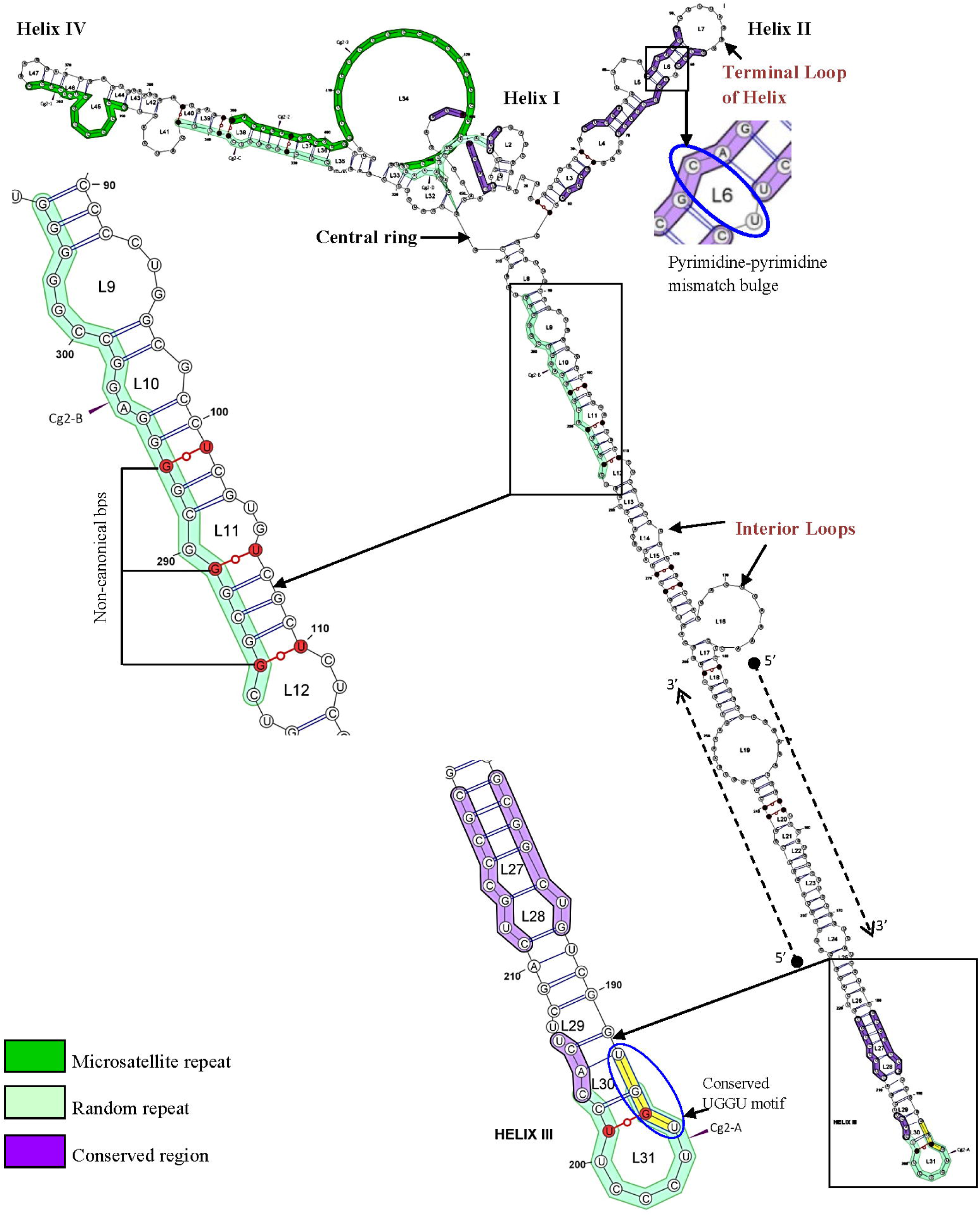
Structural organisation of ITS2 secondary structure *(Clarias gariepinus).* The 4-helix secondary structure showing the central ring with its four helices from Helix I to Helix IV projecting outward in clockwise direction. The helices are interrupted with internal loops of variable size with each helix ends into a terminal loop. The annotation was done with java-based software VARNA (Visualization Applet for RNA). Micro satellite repeats, Random repeats, eukaryotic conserved motifs, catfish conserved motifs, non-canonical bps and pyrimidine-pyrimidine mismatch are shown in magnified view round the secondary structure.

**Figure 4.**
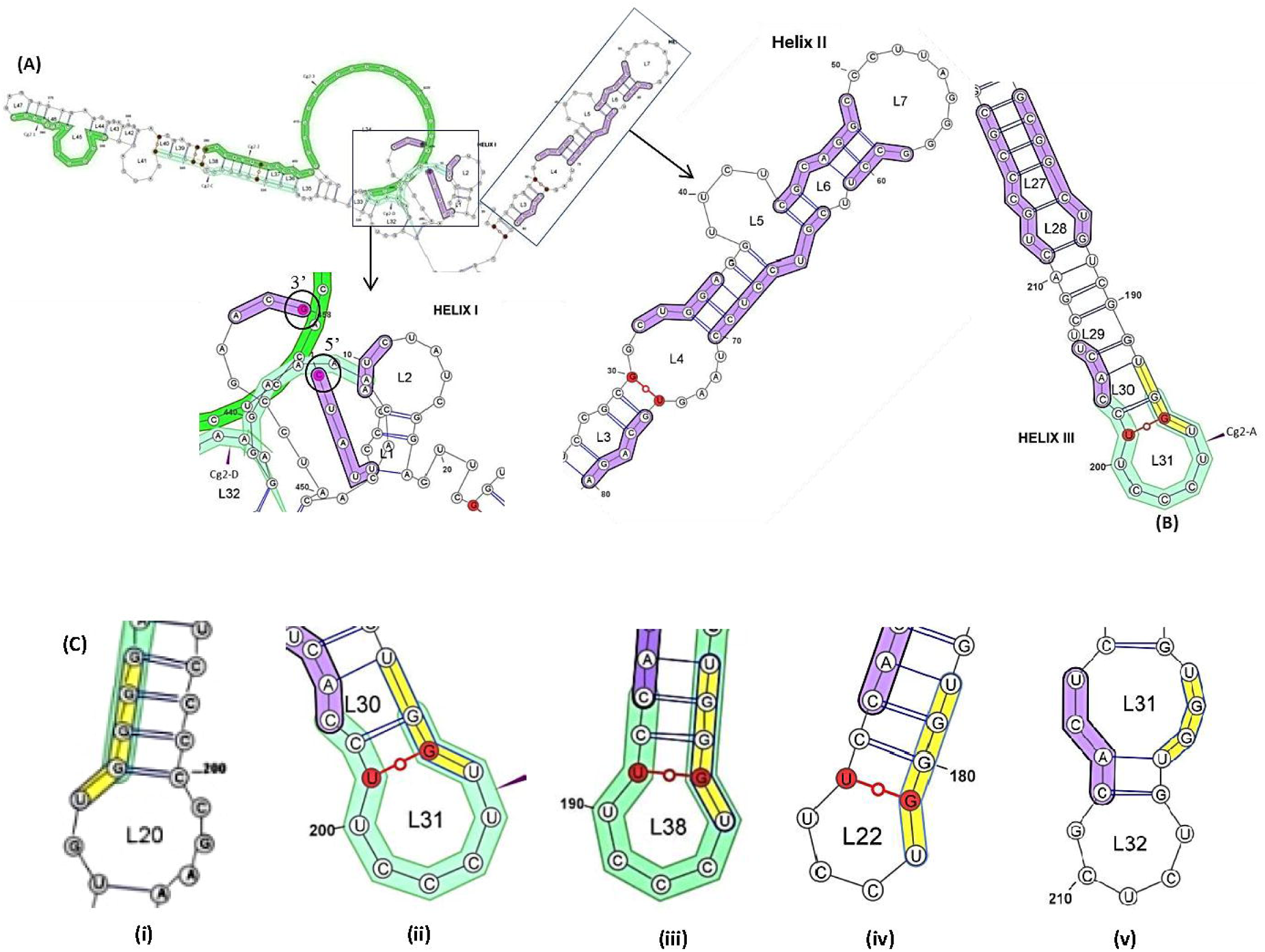
Conserved nucleotide sequences on ITS2 secondary structure. Catfish conserved regions are highlighted in blue where fig. (A) showing the conserved sequences on 5’ (CUAUU) and 3’ end (ACG), helix I (AAUC) and II (CUGGA, CGCAGGC, GCCU, CGUCCUCC, GCAGA); fig. (B) conserved sequences on the helix III (GCGGCUG, CACU, CUGCCCGC); and fig. (C) showing the eukaryotic conserved UG motif highlighted in yellow: (i) UGGGG in *C. batrachus,* (ii) UGGU in *C. gareipinus,* (iii) UGGGU in *H.ſossilis,* (iv) UGGGU in 5. *seenghala,* (v) UGGU in *W. attu.*

In addition to the eukaryotic conserved motifs, the secondary structure based multiple sequence alignment of the ITS2 have shown the presence of nine motifs which are conserved between these catfish species (Fig. 4).

#### Loops

The helices are interrupted with loops of different sizes at multiple positions which are called as interior loops. Along with intermittent loops all four helices ending into terminal loops as well (Fig. 3). The helix III possesses the maximum number of loops while the helix I possess the lowest number of loops; the position, number and the pattern of the loops in helices varied with species and together showing a species specific pattern. The four helices have shown a considerably wider range of loops between different species (Supplementary Table S1).

#### Non-canonical base-pairs

Non-canonical base pairing goes against the standard Watson & Crick base pairing pattern of DNA and RNA, where Adenine(A) could bind with Cytosine(C), and Guanine(G) could bind with Uracil(U) (Fig. 3). All the species have a shown a distinct pattern of non-canonical base pairing where only one type of base pairing i.e. G-U is found in all the secondary structure with only a single adenine with cytosine (A-C) in the helix IV of all the individuals of *Wallago attu*. All four helices in all the species are found prevalent with only G-U that appeared as a very common non-canonical base pair in ITS2 secondary structure. It ranged from a lowest of 05 to a highest of 15 G-U bp in *H. fossilis* and *C. gariepinus*, respectively (Supplementary Table S2).

#### Length of helices

The ITS2 secondary structure possessed considerable differences among its four helices which was clearly evident in all the species (Supplementary Table S3, Fig. 3). The helix III was found the longest helix in all species while the helix I was the shortest one with the exception of *S. seenghala* where helix IV was the shortest. Besides this huge difference between different helices within the same species, each helix was found with notable length range between different species. The helix I varied over a range of 06 nucleotides in *W. attu* to 33 nucleotides in *C. batrachus*, Helix II ranged from 44 nucleotides in *S. seenghala* to 105 nucleotides again in *C. batrachus*, the helix III was found varied over a comparatively higher range from 114 to 288 nucleotides in *C. batrachus* and *C. gariepinus*, respectively, and the helix IV also varied over a wider range with merely 13 nucleotides in *S. seenghala* to a count of 137 nucleotides in *C. gariepinus*. These wider length differences are because ITS2 was found to vary over a wider nucleotide range as previously discussed. Despite these major differences, no species has shown any deviation from the general attribute of ITS2 secondary structure, thus emphasizing the general structural conservation as reported in other eukaryotes (Schultz et al. 2005). Helix branching is also reported in several species in eukaryotes, and here among catfishes, single branching was found in helix II of *C. batrachus*.

#### Secondary Structure of ITS1

The secondary structure of ITS1 in five catfish species has shown a structural analogy with that of ITS2 with a resemblance to the overall contour but highly distinct structural attributes. All the secondary structures, in general, possesses a long helix similar in appearance to helix III of ITS2 and rest all helices are gathered very near to each other on the opposite end of the structure in a more or less similar manner to the secondary structure of ITS2 (Fig. 5), although the origin of ITS1 structure from the parent rRNA gene transcript was appeared different from that of ITS2 where the 5’ and 3’ points of attachment were located on one side of the structure unlike the presence of central ring and stalk in ITS2 (Fig. 6). Additionally, unlike ITS2, no eukaryotic structural hallmarks have been reported in ITS1. The helix branching is very extensive in ITS1, and the number of helices were also found to be quite variable between the five species, unlike the fixed number of helices in ITS2. Keeping in view the high degree of variability in ITS1 secondary structures, and no previous related literature on catfish species, the structure has been divided into two parts for ease of understanding, i.e. upper branched region which is attached directly to the parent rRNA gene transcript and the lower tailed region which consist of long outward helix. The pattern of helix branching having major differences between species has given a unique secondary structure to the ITS1 of each catfish species. Additionally, unlike ITS2, no conserved sequence motif was found in multiple sequence alignment of ITS1 secondary structures. (supplementary data S4).

**Figure 5.**
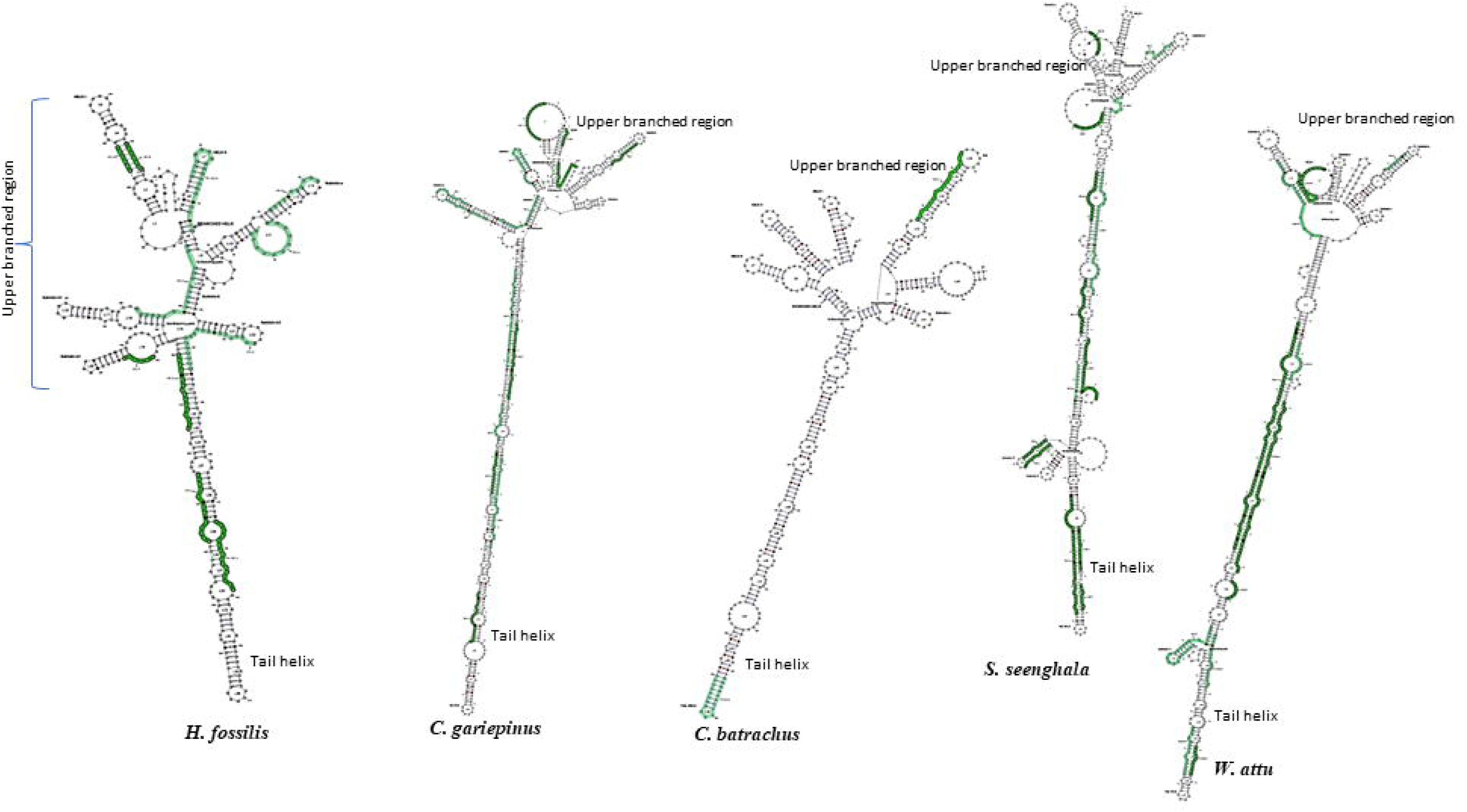
Secondary structure of ITS1. The ITS1 of *H fossilis, C. gariepinus, C. batrchus, S seenghala, and W attu showing* a similar secondary structwe without any fix number of helices unlike ITS2, with a upper branched region and a long tail helix.

**Figure 6.**
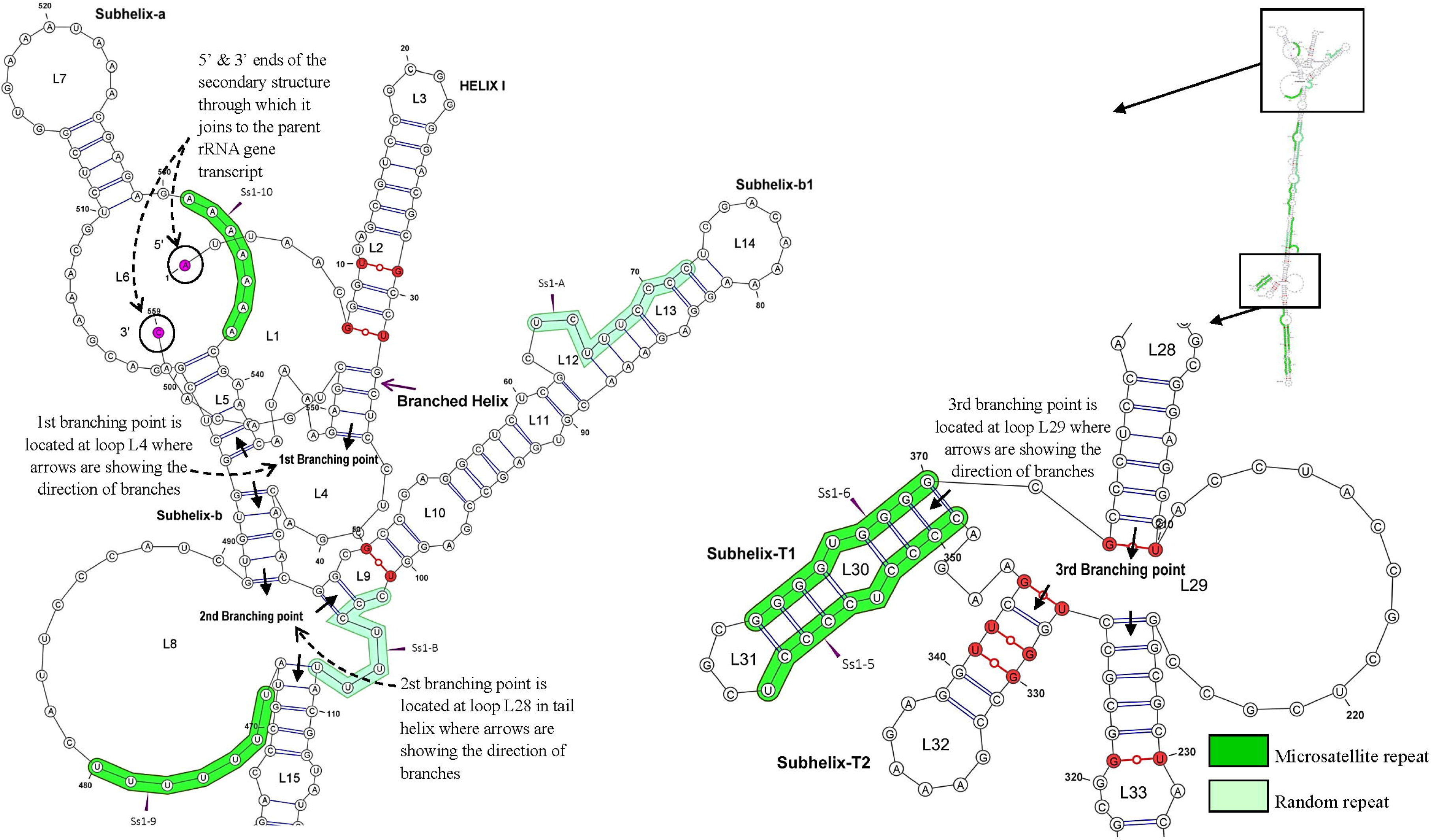
Structural organisation ofITSI secondary structure *(Sperata seenghala).* The ITS1 secondary structure is composed of multiple helices with one large branched helix. The helices are interrupted with loops (L) of variable size. The arrow → indicates the direction of the branching. The annotation was done with java-based software VARNA. Microsatellite repeat, Random repeat, branching points, non-canonical bp are shown in magnified view round the secondary structure.

#### Loops and Non-canonical basepairs

The loops and non-canonical bps in ITS1 have shown a trend similar to that of ITS2 with a species-specific pattern. The internal and terminal loops dispersed with variable positions, numbers and patterns in different species. The non-canonical G-U basepair in the only form abundant in ITS1 with different distribution pattern in each species. The minimum 12 G-U bp and maximum 21 G-U bp were present in *W. attu* and *C. gariepinus*, respectively. (supplementary Table S4).

## DISCUSSION

The structural attributes associated with the rDNA internal spacers found exclusive to the point that they provided visible distinction between them in different species in terms of their repeat distribution and secondary structures which together formed a unique rDNA signature for each species. At the primary level, their nucleotide sequences found abundant with the one highly variable attribute i.e. repeated sequences. The repeats are also well reported from diverse taxa including corals (Grajales et al. 2007), nematodes (Conole et al. 2001), beetles (Gomez-Zurita et al. 2000), and fishes (Sajdak and Phillips 1997; Chow et al. 2006). But there was no previous report of repeat distribution as observed in these catfish species, even the type and distribution of nucleotide repeats was similar between multiple individuals of each species in terms of no. of repeats and position of repeats, and forming a pattern similar in trend but unique in each species in both the internal spacers. The presence of diverse combination of microsatellite and random repeats in spacer sequences have shown their diversity of sequence motifs within the same nucleotide string which possess the loci of different sequence combination that could be as perfect as the mononucleotide repeat of thymine (T_12_) to the highly complex microsatellite repeat as A_6_(GA)_2_A_6_(GA)_7_CG_11_ in *W. attu* along with strings of complex random repeats such as T_2_C_4_G_2_A_3_C_4_GAG_3_CG_4_A_2_G_3_ which is without any basic repeating unit like microsatellite repeats. Although exception was found in *H. fossilis* ITS2 which was abundant with only randomly repeated sequences but that too was conserved within species.

The possible factor behind this uniformity in variability in the multi-chromosomal rRNA gene is the pattern of its evolution i.e. the concerted evolution, which involves homogenization of ribosomal gene from different chromosomes (Elder and Turner 1995), preventing intragenomic ITS variant due to multichromosomal origin, allowing only the accumulation of interspecific difference (Hillis and Dixon 1991). Although the concerted evolution is reported as the general trend in eukaryotes emphasizing on the complete homogenization process, it is not universal, several cases of incomplete and non-concerted evolution are also reported reflecting towards the incomplete homogenization of ribosomal gene (Eickbush and Eickbush 2007). In evolutionary perspective, the rDNA shows a range of non-concerted, incomplete concerted to concerted pattern of evolution depending upon the execution of the process of homogenization which is influenced by the genetic and epigenetic factors (Wang et al. 2023). Since we got only single band in multiple individuals of each species, it shows the absence of intragenomic length variation which in turn refers to the complete homogenization necessary for concerted evolution, thus emphasizing on the presence of concerted mode of evolution of rDNA in catfish species. But as already said that homogenization process is influenced by several factor, so at this point the wide compatibility of the primers designed in this study with 159 genera of fish species further adds to the prospect of exploring these dynamic rDNA internal spacer for the better evolutionary understanding of non-coding rDNA not just in fish species but for the vertebrates in general.

Although both ITS1 and ITS2 spacers showed a similar trend in their intra– and interspecific variabilities, both the spacers found highly diverse to each other. This diversity of ITS concealed in the nucleotide sequences was impossible to visualize without secondary structures which defined their unique sequences composition hidden in the plain sight among the sequence repeats. The spatial geometry of secondary structures and its functional relevance associated with evolutionary conserved structural attributes (Pleyte et al. 1992; Cote and Peculis 2001; Abeyrathne et al. 2002) appears as the factor responsible for the degree of uniformity observed in both the internal spacers between different catfish species. But within the limits of uniformity both ITS1 and ITS2 possess considerable differences between all five species in terms of different secondary structure attributes viz. helix length, no. of loops, non-canonical bp, distribution of repeats and conserved motif on helices which together contributed distinguished contour to each species, thus provided a relatively good amount of distinguishing information for each species both at their primary and secondary structure levels.

Out of the two spacers, ITS2 has been well reported for interspecies differentiation in eukaryotes and recommended for better taxonomic inferences when used along with its 4-helix secondary structure which is evolutionary conserved across the eukaryotes (Mai and Coleman 1997; Joseph et al. 1999; Chen et al. 2004; Young and Coleman 2004; Schultz et al. 2005; Schultz and Wolf 2009), hence found conserved between these catfish species as well. It allows the proper multiple alignment between species even when the sequences are highly diverged and could not be aligned at primary structural level (13), which is clearly observed in this study where the multi-species alignment of ITS2 secondary structures showed the presence of nine more conserved sequence motifs between five catfish species in addition to the standard eukaryotic conserved motifs at the same time when their primary structure failed to align between species. Moreover, within this conserved structural contour each species carries its unique attributes viz. helix I was longest in *C. batrachus* with branched helix II, *C. gariepinus* was found consisting of helix IV with consecutive looping, *H. fossilis* with a pattern in helix I and IV similar to that of *C. gariepinus* but unlike other species it has a unique pattern of three consecutive loop with just four bonded pairs. Unique to *S. seenghala,* the central ring which was not a perfect ring structure even in other catfish species was also present as bi-lobed structure with a comparatively long stalk, and also unlike other species helix IV was shortest in *S. seenghala*. Likewise, the unique structural characteristic in *W. attu* was the excessive looping specially in the helix III, near to the central ring. Hence this diversity in uniformity is the key to successful utilization of ITS2 for the differentiation of catfish species.

The structural uniformity was also traced in the secondary structure of ITS1, although contrary to ITS2, absence of any standard eukaryotic ITS1 secondary structure makes it more diverse which was quite apparently having no interspecific conserved region between five catfish species. But at the same time structural conservation was well obvious in ITS1 as well with upper branched region and a long tail helix, though the branching is diverged so much between species that it formed a unique pattern of helices and sub-helices in each species, thus providing species diversity among the structural uniformity in ITS1 on a pattern similar to ITS2.

Conclusively, the diverse structures of both the internal spacers have differentiated all five catfish species and also hold the potential to contribute in the molecular genetics and systematics of not only catfish species but other fish species in general, and could also contribute for the other vertebrate species as they share similar process of ribosomal maturation where these ITS regions play a very crucial role by offering their secondary structures as potential binding sites for several maturation factors such as snoRNA and RAC protein. The functional dynamics of diverse repeat patterns in internal spacers in the conserved maturation process is a thing to be explored further.

Although there are several reports of ITS from fish species, but overall, there are limited studies in vertebrates from pisces to mammals, and this is the first time such a detailed description has been given in any vertebrate species about its composition of short sequence repeats. Such a dynamic sequence composition in internal spacers reflect towards their possible sequence diversity in the vertebrate groups. Since the pisces are the earliest in vertebrate evolution followed by amphibia, reptilia, aves and mammalia, the comparative repeats distributions among the higher vertebrates could give new insights into the evolution of rDNA as well as non-coding DNA in general which is known by the abundance of repeated sequences. Although the methodology is quite very extensive, the picture which come out in terms of repeat distribution and secondary structure makes ITS a very promising DNA segment which hold the key to species delimitation and discovery in vertebrates.

### Accession numbers

“ITS1 GenBank accession nos. MT105349, MT105350, MT105351, MT105352, MT105353, MT105354, MT105355, MT105356, MT105357, MT105358, MT105359, MT105360, MT105361, MT105362, MT105363, MT105364, MT105365.

ITS2 GenBank accession nos. MT234369, MT234372, MT234373, MT234374, MT234376, MT234377, MT234378, MT234380, MT234381, MT234382, MT234383, MT234384, MT234385, MT234386, MT234387.

### Authors Contribution

**Mohd Imran:** Conceptualization; Methodology; Formal analysis; Funding acquisition; Writing –original draft; review & editing. **Sana Nafees:** Conceptualization; Methodology, Formal analysis; review & editing.

## Acknowledgments

The Authors are especially grateful to Prof. Mohammad Afzal for his constructive insights into this work. The authors also acknowledge the Department of Zoology, Aligarh Muslim University, for providing necessary laboratory facilities. The Council of Scientific and Industrial Research (CSIR), New Delhi, India is also gratefully acknowledged for providing funding support (09/112(0603)/2018-EMR-I).

