## Supplementary material for "Exploring unique sequence repeat patterns and secondary structures in rDNA internal transcribed spacers ITS1 and ITS2 for characterization of catfish species": Supplementary Data S1.docx

**Supplemental Data S1**

**Primer Designing**

To deal with the questions under study, we had to target the ITS1 and ITS2 specifically. No such primers were available that could specifically target only the spacer regions of the species investigated. So, to accomplish this task, we designed our own primers and also checked their compatibility with numerous other species available in the NCBI database for their broader application.

The almost complete rDNA cistron of *Clarias gariepinus* was taken from NCBI (accession no. AJ876383.1), and the locations of ITS1 and ITS2 were traced. The primers were designed using NCBI primer designing tool (https://www.ncbi.nlm.nih.gov/tools/primer-blast/) keeping the default parameters (Ye et al. 2012). The DNA sequencing aspect was also kept in mind during the process. The appropriate primer regions were located keeping in mind that the 30-40 nucleotides segment at the 5’ end and 15-20 nucleotides at the 3’end of a sequence always receives low resolution in DNA sequencing process and hence could be trimmed without any loss of information. So, accordingly, both forward and reverse primer ranges at the respective 5’ and 3’ end positioned at a distance of 40-50 nucleotides from the target locus. The nucleotide range for primers was targeted with an emphasis on other important parameters like (i) length of primers kept approx. 18-24 nucleotides, (ii) GC content kept within 40-60%, and (iii) melting temperature (Tm) in the range of 55-65°C with Tm of both the primer kept within 5°C difference. So, out of the various possible set of the primers generated by the primer tool, the primers were finalized accordingly. After designing, the primers were sent for synthesis to the commercially available facility at Integrated DNA Technologies, Inc. USA.

**Table A:**	Newly designed primers for ITS1 & ITS2 using NCBI primer designing tool.

| **Primer** | **Sequence (5’-3’)** | **Tm (°C)** | **GC(%)** |
| --- | --- | --- | --- |
| ITS.1.F1 (ITS1 forward 1) | TCG ATC AAA CTT GAC TAT CTA GAG G | 53.6 | 40.0 |
| ITS.1.F2 (ITS1 forward 2) | CGA TCA AAC TTG ACT ATC TAG AGG A | 53.6 | 40.0 |
| ITS.1.R1 (ITS1 reverse) | CAG CTG GCT GCG TTC TTC AT | 58.3 | 55.0 |
| ITS2.F (ITS2 forward) | CAG GAC ACA TTG ATC ATC GAC AC | 55.8 | 47.8 |
| ITS2.R (ITS2 reverse) | AAT TCA GCG GGT CGT CTC | 54.9 | 55.6 |

**Table B:**	Standardized PCR conditions for ITS1& ITS2.

| **Primer** | **Species** | **MgCl2 (mM)** | **Ta (°C)/Sec.** | **PCR Product^¶^ (bp)** |
| --- | --- | --- | --- | --- |
|  | *C. batrachus* | 3.5 | 53°C /60sec | ~500 |
| ITS.1.F1/F2 | *C. gariepinus* | 1.5 | 53°C /60sec | ~500 |
| ITS.1.R1 | *H. fossilis* | 2.5 | 49°C /60sec | 500-600 |
|  | *S. seenghala* | 2.5 | 49°C /60sec | ~700 |
|  | *W. attu* | 2.5 | 49°C /60sec | ~700 |
|  | *C. batrachus* | 1.5 | 50°C/45sec | 400-500 |
| ITS2.F | *C. gariepinus* | 1.5 | 50°C/45sec | ~600 |
| ITS2.R | *H. fossilis* | 1.5 | 50°C/45sec | ~500 |
|  | *S. seenghala* | 1.5 | 50°C/45sec | ~500 |
|  | *W. attu* | 1.5 | 50°C/45sec | ~500 |

^¶^ include small rDNA segment at 5’and 3’ ends

**Primer compatibility**

Since the primers had given good and specific amplifications, we decided to check their compatibility with other species to figure out their broad application. The primers complementary sequences were traced in numerous species in the NCBI database with a maximum target of 1000 subjects. All three rRNA genes (18S, 5.8S & 28S) data were checked individually- sequence by sequence, species by species, to find those species having the corresponding forward and reverse primer complementary sequences for both ITS1 and ITS2.

**Table C:**	Compatibility of newly designed ITS1 and ITS2 primers with sequences of fish species available on NCBI database. Species from 159 genera are found with compatible primer regions and 37 genera are found with compatibility in multiple sister species. [─] indicate data not found at NCBI database.

| ***S.No.*** | ***Species***  **(Family)** | **18S rRNA gene**  **(ITS1F)** | **5.8S rRNA gene**  **(ITS1R and ITS2F)** | **28S**  **rRNA gene**  **(ITS2R)** | **Genbank Accession no.**  **(18s; 5.8s; 28s rRNA gene)** |
| --- | --- | --- | --- | --- | --- |
| 1. | *Dinotopterus cunningtoni*  (Clariidae) | ✓ | ✓ | ✓ | AJ876385.1 |
| 2. | *Tanganikallabes spp.*  (Clariidae) | ✓ | ✓ | ✓ | JF297969.1; AJ876380.1 |
| 3. | *Channallabes spp.*  (Clariidae) | ✓ | ✓ | ✓ | AJ876391.1; AJ876393.1; AJ876401.1 |
| 4. | *Clariallabes longicauda*  (Clariidae) | ✓ | ✓ | ✓ | AJ876390.1; AJ876389.1; AJ876390.1 |
| 5. | *Heterobranchus isopterus*  (Clariidae) | ✓ | ✓ | ✓ | AJ876382.1 |
| 6. | *Pangasianodon hypophthalmus*  (Pangasiidae) | ✓ | ✓ | ✓ | AJ876376.1; XR_004577714.1 |
| 7. | *Kryptopterus bicirrhis*  (Siluridae) | ✓ | ✓ | ✓ | AJ876375.1 |
| 8. | *Tachysurus fulvidraco*  (bagridae) | ✓ | ✓ | ✓ | XR_003440535.1; XR_003440548.1; XR_003440545.1 |
| 9. | *Hemibagrus spp.*  (Bagridae) | ✓ | ✓ | ✓ | GQ465251.1; GQ465844.1; GQ465251.1, KC009848.1 |
| 10. | *Clarias spp.*  (Clariidae) | ✓ | ✓ | ✓ | AJ876400.1, AJ876386.1, AJ876378.1; AJ876379.1, AJ876398.1,  AJ876397.1, AJ876381.1, AJ876402.1; AJ876395.1 |
| 11. | *Pelteobagrus spp.*  (Bagridae) | ✓ | ✓ | ✓ | HM104672.1, GQ465236.1, HM104669.1; GQ465839.1 |
| 12. | *Leiocassis longirostris*  (Bagridae) | ✓ | ✓ | ✓ | GQ465843.1; GQ465246.1 |
| 13. | *Silurus asotus*  (Siluridae) | ✓ | ✓ | ✓ | GQ465841.1; GQ465243.1 |
| 14. | *Ompok spp.*  (Siluridae) | ✓ | ✓ | ─ | GU721094.1, GU385711.1, GU385714.1; GU385709.1 |
| 15. | *Ictalurus spp.*  (Ictaluridae) | ✓ | ✓ | ✓ | AF021880.1; GQ465242.1; Z18678.1 |
| 16. | *Isorineloricaria spinosissima*  (Loricariidae) | ─ | ✓ | ✓ | AJ412874.1 |
| 17. | *Hypostomus spp.*  (Loricariidae) | ─ | ✓ | ✓ | AJ412873.1, AJ412862.1, AJ412852.1, AJ412851.1, AJ412865.1,  AJ412864.1, AJ412861.1, AJ412854.1, AJ412853.1, AJ412855.1; AJ412857.1 |
| 18. | *Cochliodon spp.*  (Loricariidae) | ─ | ✓ | ✓ | AJ412866.1, AJ412867.1 |
| 19. | *Pseudopleuronectes spp.*  (Pleuronectidae) | ─ | ✓ | ✓ | MH204395.1, MH204343.1 |
| 20. | *Pterygoplichthys zuliaensis*  (Loricariidae) | ─ | ✓ | ✓ | AJ412871.1 |
| 21. | *Hemiancistrus chlorostictus*  (Loricariidae) | ─ | ✓ | ✓ | AJ412869.1 |
| 22. | *Pseudochaenichthys georgianus*  (Channichthyidae) | ✓ | ✓ | ✓ | XR_004551328.1; XR_004552560.1; XR_004551325.1 |
| 23. | *Plectropomus leopardus*  (Serranidae) | ✓ | ✓ | ✓ | AP022715.1 |
| 24. | *Epinephelus spp.*  (Serranidae) | ✓ | ✓ | ✓ | AP022698.1, JN603833.1; EF472470.1, EF472469.1, EF472468.1,  XR_004502793.1 |
| 25. | *Bothus myriaster*  (Bothidae) | ─ | ✓ | ✓ | MH509956.1 |
| 26. | *Scomber spp.*  (Scombridae) | ✓ | ✓ | ✓ | KF986714.1; LC465235.1, LC464972.1, LC464971.1, LC464970.1 |
| 27. | *Denticeps clupeoides*  (Denticipitidae) | ─ | ✓ | ✓ | XR_003742790.1; XR_003749006.1 |
| 28. | *Gouania willdenowi*  (Gobiesocidae) | ✓ | ✓ | ✓ | XR_003675558.1; XR_003675553.1; XR_003672927.1 |
| 29. | *Clidoderma asperrimum*  (Pleuronectidae) | ─ | ✓ | ✓ | MH204262.1 |
| 30. | *Cleisthenes pinetorum*  (Pleuronectidae) | ✓ | ✓ | ✓ | EF126039.1; MH204267.1 |
| 31. | *Hippoglossoides elassodon*  (Pleuronectidae) | ─ | ✓ | ✓ | MH204481.1 |
| 32. | *Eopsetta grigorjewi*  (Pleuronectidae) | ─ | ✓ | ✓ | MH204524.1 |
| 33. | *Verasper variegates*  (Pleuronectidae) | ─ | ✓ | ✓ | MH204446.1 |
| 34. | *Platichthys stellatus*  (Pleuronectidae) | ✓ | ✓ | ✓ | MG904953.1; MH204386.1 |
| 35. | *Lepidopsetta polyxystra*  (Pleuronectidae) | ─ | ✓ | ✓ | MH204293.1; MH204295.1 |
| 36. | *Limanda aspera*  (Pleuronectidae) | ─ | ✓ | ✓ | MH204280.1 |
| 37. | *Sphaeramia orbicularis*  (Apogonidae) | ✓ | ✓ | ✓ | XR_003934901.1; LR597473.1; XR_003935038.1 |
| 38. | *Crossorhombus kobensis*  (Bothidae) | ✓ | ✓ | ✓ | MH036908.1; MH036924.1 |
| 39. | *Neostethus bicornis*  (Phallostethidae) | ✓ | ✓ | ✓ | LR738553.1 |
| 40. | *Engyprosopon grandisquama*  (Bothidae) | ✓ | ✓ | ✓ | MH179080.1; MH179076.1 |
| 41. | *Arnoglossus macrolophus*  ((Bothidae) | ✓ | ✓ | ✓ | MH179078.1; MH179074.1 |
| 42. | *Dicentrarchus labrax*  (Moronidae) | ✓ | ✓ | ─ | HM014387.1; LC387525.1 |
| 43. | *Maccullochella macquariensis*  (Percichthyidae) | ✓ | ✓ | ─ | KX342081.1 |
| 44. | *Channa spp.*  (Channidae) | ─ | ✓ | ─ | KJ451611.1, KJ451605.1, KJ451601.1 |
| 45. | *Nothobranchius furzeri*  (Nothobranchiidae) | ✓ | ✓ | ✓ | EU780557.1 |
| 46. | *Serranus subligarius*  (Serranidae) | ─ | ✓ | ─ | EF472500.1, EF472464.1 |
| 47. | *Centropristis spp.*  (Serranidae) | ─ | ✓ | ─ | EF472499.1, EF472490.1, EF472480.1 |
| 48. | *Diplodus holbrooki*  (Sparidae) | ─ | ✓ | ─ | EF472471.1 |
| 49. | *Mycteroperca microlepis*  (Serranidae) | ─ | ✓ | ─ | EF472466.1 |
| 50. | *Serraniculus pumilio*  (Serranidae) | ─ | ✓ | ─ | EF472465.1 |
| 51. | *Kareius bicoloratus*  (Pleuronectidae) | ✓ | ✓ | ─ | EU647401.1 |
| 52. | *Anoplopoma fimbria*  (Anoplopomatidae) | ─ | ✓ | ─ | AB244631.1 |
| 53. | *Auxis rochei*  (Scombridae) | ✓ | ✓ | ✓ | AB193742.1 |
| 54. | *Aphanotorulus ammophilus*  (Loricariidae) | ─ | ✓ | ✓ | AJ412872.1 |
| 55. | *Macquaria australasica*  (Percichthyidae) | ✓ | ✓ | ─ | KX342084.1 |
| 56. | *Symphurus plaguisa*  (Cynoglossidae) | ─ | ✓ | ─ | MG026869.1 |
| 57. | *Tribolodon spp.*  (Cyprinidae) | ─ | ✓ | ─ | KY250509.1, KY250512.1, KY250494.1 |
| 58. | *Myoxocephalus spp.*  (Cottidae) | ✓ | ✓ | ✓ | MT497872.1, MT497870.1, MT497867.1, MT497865.1, MT497869.1 |
| 59. | *Microcottus sellaris*  (Cottidae) | ✓ | ✓ | ✓ | MT497863.1 |
| 60. | *Porocottus spp.*  (Cottidae) | ✓ | ✓ | ✓ | MT497862.1, MT497857.1, MT497859.1 |
| 61. | *Megalocottus platycephalus*  (Cottidae) | ✓ | ✓ | ✓ | MT484272.1 |
| 62. | *Hippoglossus spp.*  (Pleuronectidae) | ✓ | ✓ | ✓ | XR_004613416.1; XR_004613415.1, XR_004697965.1; XR_004613417.1,  XR_004697937.1 |
| 63. | *Cyclopterus lumpus*  (Cyclopteridae) | ─ | ✓ | ─ | XR_004608934.1 |
| 64. | *Gymnodraco acuticeps*  (Bathydraconidae) | ✓ | ✓ | ─ | XR_004567119.1; XR_004567118.1 |
| 65. | *Periophthalmus magnuspinnatus*  (Gobiidae) | ✓ | ✓ | ─ | XR_004542209.1; XR_004542202.1 |
| 66. | *Paraplagusia japonica*  (Cynoglossidae) | ─ | ✓ | ✓ | MK679720.1 |
| 67. | *Salarias fasciatus*  (Blenniidae) | ✓ | ✓ | ✓ | XR_003930914.1; XR_003930922.1; XR_003930553.1 |
| 68. | *Myripristis murdjan*  (Holocentridae) | ✓ | ✓ | ✓ | XR_003928004.1; XR_003928002.1; XR_003928005.1 |
| 69. | *Cottoperca gobio*  (Bovichtidae) | ✓ | ✓ | ─ | XR_003832029.1; XR_003832028.1 |
| 70. | *Labrus bergylta*  (Labridae) | ✓ | ✓ | ✓ | XR_003807100.1; XR_003807867.1; XR_003808014.1 |
| 71. | *Parambassis ranga*  (Ambassidae) | ✓ | ✓ | ─ | XR_003670172.1; XR_003670170.1 |
| 72. | *Larimichthys crocea*  (Sciaenidae) | ─ | ✓ | ─ | XR_003463126.1 |
| 73. | *Apolemichthys spp.*  (Pomacanthidae) | ✓ | ✓ | ✓ | KU363798.1, KU244245.1, KU244239.1, KU240018.1 |
| 74. | *Centropyge spp.*  (Pomacanthidae) | ✓ | ✓ | ✓ | KU363797.1, KU356792.1, KU356790.1, KU356788.1, KU356787.1,  KU356786.1, KU356785.1, KU356784.1, KU356783.1, KU356781.1,  KU356780.1, KU244251.1, KU244249.1 |
| 75. | *Holacanthus spp.*  (Pomacanthidae) | ✓ | ✓ | ✓ | KU356793.1, KU244250.1, KU244247.1, KU244246.1,  KU244238.1 |
| 76. | *Gemicanthus spp.*  (Pomacanthidae) | ✓ | ✓ | ✓ | KU356791.1, KU356789.1, KU244253.1, KU244252.1 |
| 77. | *Paracentropyge multifasciata*  (Pomacanthidae) | ✓ | ✓ | ✓ | KU356779.1 |
| 78. | *Pygoplites diacanthus*  (Pomacanthidae) | ✓ | ✓ | ✓ | KU244248.1; KU244248.1; |
| 79. | *Chaetodontoplus spp.*  (Pomacanthidae) | ✓ | ✓ | ✓ | KU244244.1, KU244241.1 |
| 80. | *Pomacanthus spp.*  (Pomacanthidae) | ✓ | ✓ | ✓ | KU244243.1, KU244242.1 |
| 81. | *Synechogobius ommaturus*  (Gobiidae) | ✓ | ✓ | ✓ | KP893562.1 |
| 82. | *Pleurogrammus spp.*  (Hexagrammidae) | ─ | ✓ | ─ | AB244630.1, AB244629.1 |
| 83. | *Hexagrammos spp.*  (Hexagrammidae) | ─ | ✓ | ─ | AB244628.1, AB244627.1, AB244626.1, AB244625.1, AB244624.1,  AB244623.1 |
| 84. | *Takifugu rubripes*  (Tetraodontidae) | ─ | ✓ | ✓ | XR_003887403.1, XR_003887306.1 |
| 85. | *Gymnocypris przewalskii*  (Cyprinidae) | ✓ | ─ | ✓ | MN909518.1 |
| 86. | *Chanos chanos*  (Chanidae) | ✓ | ─ | ✓ | XR_004025993.1; XR_004025972.1 |
| 87. | *Schizothorax prenanti*  (Cyprinidae) | ✓ | ─ | ─ | MK439424.1 |
| 88. | *Neolissochilus hexagonolepis*  (Cyprinidae) | ✓ | ─ | ─ | GU568380.1 |
| 89. | *Tor mosal* *mahanadicus*  (Cyprinidae) | ✓ | ─ | ─ | GU568374.1, GU568359.1, GU568354.1, GU568370.2,  GU568371.1, GU568364.1 |
| 90. | *Engraulis encrasicolus*  (Engraulidae) | ✓ | ─ | ─ | MF796393.1 |
| 91. | *Onychostoma spp.*  (Cyprinidae) | ✓ | ─ | ─ | MF598162.1, MF598161.1 |
| 92. | *Puntius chelynoides*  (Cyprinidae) | ✓ | ─ | ─ | GU568345.1 |
| 93. | *Oreochromis niloticus*  (Cichlidae) | ✓ | ─ | ─ | MF460347.1, GU289229.1 |
| 94. | *Sprattus sprattus*  (Clupeidae) | ✓ | ─ | ─ | MF796435.1 |
| 95. | *Hemibarbus labeo*  (Cyprinidae) | ✓ | ─ | ─ | MH843153.1 |
| 96. | *Tinca tinca*  (Cyprinidae) | ✓ | ─ | ─ | MH362745.1 |
| 97. | *Acipenser ruthenus*  (Acipenseridae) | ✓ | ─ | ─ | XR_004544382.2 |
| 98. | *Archocentrus centrarchus*  (Cichlidae) | ✓ | ─ | ─ | XR_004019531.1 |
| 99. | *Arnoglossus macrolophus*  (Bothidae) | ✓ | ─ | ─ | MH179078.1 |
| 100. | *Maylandia zebra*  (Cichlidae) | ✓ | ─ | ─ | XR_003024144.1 |
| 101. | *Amphiprion ocellaris*  (Pomacentridae) | ✓ | ─ | ✓ | XR_002748043.1, XR_002748044.1 |
| 102. | *Cnidoglanis macrocephalus*  (Plotosidae) | ✓ | ─ | ─ | KM588845.1 |
| 103. | *Plectorhinchus spp.*  (Haemulidae) | ✓ | ─ | ✓ | JN850649.1, JQ766314.1 |
| 104. | *Pagrus spp.*  (Sparidae) | ✓ | ─ | ─ | JN903346.1, JN903328.1 |
| 105. | *Argyrosomus regius*  (Sciaenidae) | ✓ | ─ | ─ | JF799401.1 |
| 106. | *Siganus spp.*  (Siganidae) | ✓ | ─ | ─ | AB277081.1, AB277065.1, AB277053.1, AB277035.1, AB277029.1,  AB277022.1, AB277020.1.... total 19 spp. |
| 107. | *Prionurus scalprum*  (Acanthuridae) | ✓ | ─ | ─ | AB277067.1 |
| 108. | *Xiphophorus hellerii*  (Poeciliidae) | ✓ | ─ | ✓ | XR_004336940.1, XR_003594260.1, XR_002752243.1;  XR_004336943.1, XR_003598564.1, XR_002752701.1 |
| 109. | *Erpetoichthys calabaricus*  (Polypteridae) | ✓ | ─ | ✓ | XR_003714693.1; XR_003714991.1 |
| 110. | *Aesopia cornuta*  (Soleidae) | ✓ | ─ | ─ | MG895442.1 |
| 111. | *Cynoglossus spp.*  (Cynoglossidae) | ✓ | ─ | ✓ | MG895427.1; KU177475.1 |
| 112. | *Pleuronichthys cornutus*  (Pleuronectidae) | ✓ | ─ |  | MG890467.1 |
| 113. | *Perca flavescens*  (Percidae) | ✓ | ─ | ✓ | MT458484.1, KX342086.1; Z18686.1 |
| 114. | *Coregonus spp.*  (Salmonidae) | ✓ | ─ | ✓ | MT458451.1, MT458407.1, MT458287.1; LR778282.1 |
| 115. | *Stenodus leucichthys*  (Salmonidae) | ✓ | ─ |  | MT458404.1 |
| 116. | *Amblyraja radiate*  (Rajidae) | ✓ | ─ | ✓ | XR_004411840.1; XR_004411188.1 |
| 117. | *Etheostoma spectabile*  (Percidae) | ✓ | ─ | ✓ | XR_004329252.1; XR_004329248.1 |
| 118. | *Sander lucioperca*  (Percidae) | ✓ | ─ | ✓ | XR_004104006.1; XR_004104007.1 |
| 119. | *Plagiopsetta glossa*  (Samaridae) | ✓ | ─ | ─ | MG895432.1 |
| 120. | *Samariscus latus*  (Samaridae) | ✓ | ─ | ─ | MG895431.1 |
| 121. | *Oncorhynchus mykiss*  (Salmonidae) | ✓ | ─ | ─ | KJ851798.1 |
| 122. | *Sebastes spp.* (29 spp.) (Sebastidae) | ✓ | ─ | ─ | DQ678714.1, DQ678713.1, DQ678712.1, DQ678710.1,  DQ678709.1, DQ678708.1.... * |
| 123. | *Helicolenus avius*  (Sebastidae) | ✓ | ─ | ─ | DQ678711.1 |
| 124. | *Hozukius emblemarius*  (Sebastidae) | ✓ | ─ | ─ | DQ678705.1 |
| 125. | *Tandanus bostocki* (Plotosidae) | ✓ | ─ | ─ | KJ774764.1 |
| 126. | *Porochilus spp.*  (Plotosidae) | ✓ | ─ | ─ | HQ615581.1, KJ774735.1 |
| 127. | *Herklotsichthys castelnaui*  (Clupeidae) | ✓ | ─ | ─ | KJ774685.1 |
| 128. | *Potamalosa richmondia*  (Clupeidae) | ✓ | ─ | ─ | KJ774739.1 |
| 129. | *Misgurnus anguillicaudatus*  (Cobitidae) | ✓ | ─ | ─ | KJ774694.1 |
| 130. | *Clupea harengus*  (Clupeidae) | ✓ | ─ | ✓ | X98845.1; Z18764.1 |
| 131. | *Culter alburnus*  (Cyprinidae) | ✓ | ─ | ─ | AB860216.1 |
| 132. | *Megalobrama amblycephala*  (Cyprinidae) | ✓ | ─ | ✓ | AB860215.1; MG830472.1 |
| 133. | *Anguilla spp.*  (Anguillidae) | ✓ | ─ | ✓ | FM946070.1, FM946071.1; XR_004766723.1, U34342.1 |
| 134. | *Salmo trutta*  (Salmonidae) | ✓ | ─ | ✓ | XR_003880455.1; XR_003868235.1 |
| 135. | *Anarrhichthys ocellatus*  (Anarhichadidae) | ✓ | ─ | ✓ | XR_004213541.1; XR_004212307.1 |
| 136. | *Cyprinodon variegatus*  (Cyprinodontidae) | ✓ | ─ | ─ | EF431912.1 |
| 137. | *Pseudorhombus arsius*  (Paralichthyidae) | ✓ | ─ | ─ | MG895443.1 |
| 138. | *Colistium guntheri*  (Pleuronectidae) | ✓ | ─ | ─ | MG895430.1 |
| 139. | *Solea senegalensis*  (Soleidae) | ✓ | ─ | ─ | EF126042.1 |
| 140. | *Cirrhinus mrigala*  (Cyprinidae) | ✓ | ─ | ─ | GU967675.1 |
| 141. | *Hyporhamphus australis*  (Hemiramphidae) | ✓ | ─ | ─ | KJ774686.1 |
| 142. | *Acanthopagrus spp.*  (Sparidae) | ✓ | ─ | ✓ | KJ774634.1; FJ960540.1 |
| 143. | *Gambusia holbrooki*  (Poeciliidae) | ✓ | ─ | ─ | FJ710842.1 |
| 144. | *Phalloceros caudimaculatus*  (Poeciliidae) | ✓ | ─ | ─ | KJ774729.1 |
| 145. | *Geophagus sp.*  (Cichlidae) | ✓ | ─ | ─ | KJ774680.1 |
| 146. | *Rocio octofasciata*  (Cichlidae) | ✓ | ─ | ─ | KJ774653.1 |
| 147. | *Amatitlania nigrofasciata*  (Cichlidae) | ✓ | ─ | ─ | KJ774642.1 |
| 148. | *Polypterus senegalus*  (Polypteridae) | ✓ | ─ | ─ | AF188367.1 |
| 149. | *Liza parsia*  (Mugilidae) | ─ | ─ | ✓ | JQ814469.1 |
| 150. | *Lophius piscatorius*  (Lophiidae) | ─ | ─ | ✓ | Z18765.1 |
| 151. | *Anabas testudineus*  (Anabantidae) | ─ | ─ | ✓ | LR132047.1 |
| 152. | *Trigla lucerna*  (Triglidae) | ─ | ─ | ✓ | Z18768.1 |
| 153. | *Sardina pilchardus*  (Clupeidae) | ─ | ─ | ✓ | Z18767.1 |
| 154. | *Pethia spp.*  (Cyprinidae) | ─ | ─ | ✓ | KJ509176.1, KJ509180.1 |
| 155. | *Pampus argenteus*  (Stromateidae) | ─ | ─ | ✓ | FJ960553.1 |
| 156. | *Mystus spp.*  (Bagridae) | ─ | ─ | ✓ | KC009847.1, KC009843.1, KC009842.1 |
| 157. | *Astatotilapia calliptera* (Cichlidae) | ─ | ─ | ✓ | XR_003270566.1 |
| 158. | *Thalassophryne amazonica*  (Batrachoididae) | ─ | ─ | ✓ | XR_004566219.1 |
| 159. | *Syngnathus acus*  (Syngnathidae) | ─ | ─ | ✓ | LR594598.1 |
