## Supplementary material for "Exploring unique sequence repeat patterns and secondary structures in rDNA internal transcribed spacers ITS1 and ITS2 for characterization of catfish species": Supplementary Data S2.docx

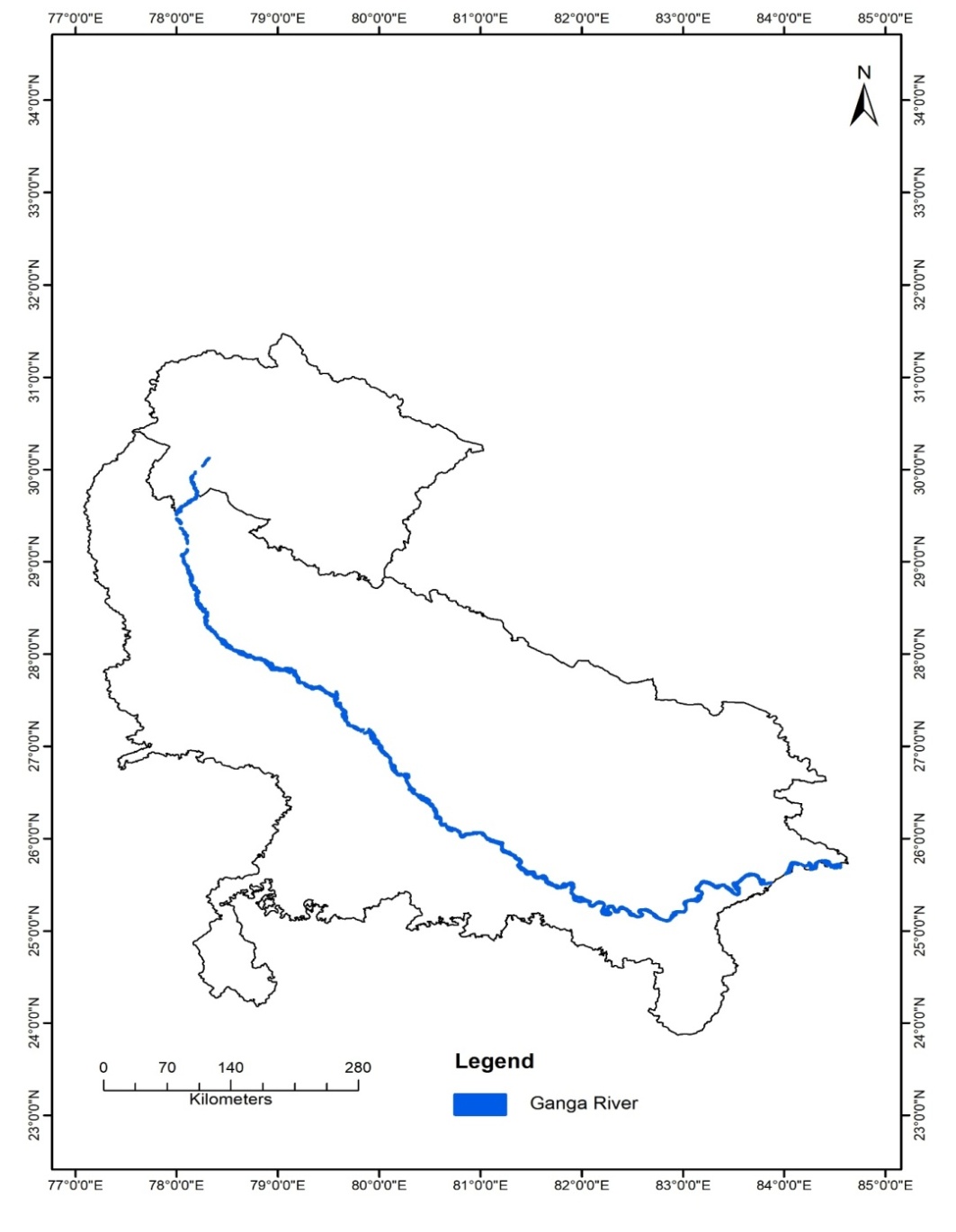

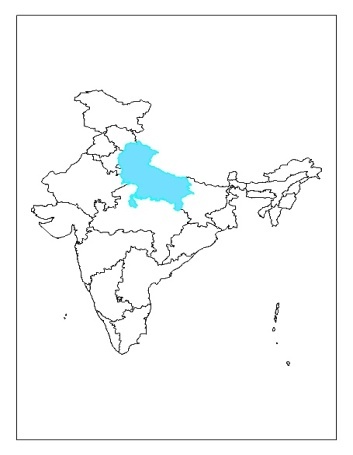

Garhwal

Aligarh

Varanasi

Sampling Site

**Sampling sites:** The three sampling sites- Garhwal, Aligarh and Varanasi on the stretch of river Ganga in the north India (shown in inset).

**Secondary Structure Analysis ITS2**

***ITS2 database***

The ITS2 database holds information about sequences, secondary structures and taxonomic classifications of all the ITS2 sequences in GenBank (Koetschan et al. 2010). It performs the identification and delineation of the ITS2 sequences. The ITS2 database together with its web server works as an efficient tool for the secondary structures analysis. It harbours key tools like Blast, Annotate sequence, Predict sequence, Model sequence, Multiple alignment on sequence & structure and search for conserved Motifs. The multiple sequence alignment with their corresponding secondary structures allows performing secondary structure based analysis between species with an emphasis on secondary structure homology.

- **Annotation of ITS2 sequences**

The process of annotation involves the demarcation of the exact location of the 5’ and 3’ end nucleotide of the ITS2 sequence in the rRNA gene transcript which consist of a continuous string of 18s rDNA-ITS1-5.8s rDNA-ITS2-28s rDNA.

This annotation process of ITS2 utilizes the conserved nature of trailing 5.8s rDNA and 28s rDNA which are highly conserved regions across the vertebrates. The annotation process traces these 5.8s rDNA and 28s rDNA at the 5’ and 3’ ends of the ITS2 sequences, respectively, to locate ITS2 sequences. This annotation portal has precisely annotated the ITS2 encompassing rDNA sequences from the multiple individual all five catfish species. (Table A). The annotation process simultaneously generated the fasta output file for all the annotated ITS2 sequences for further analysis.

**Table (A):** Annotation of the ITS2 sequences. The annotation output is presented in the fasta format. The central ITS2 sequence (in black) in amplified rDNA segment is separated from the terminal 5.8s and 28s rDNA nucleotide motifs (in grey & red, respectively). Multiple individuals are annotated to avoid any discrepancy in tracing the exact location of 5’ and 3’ end of ITS2 sequences in all the species. (a) Annotation of *C. batrachus* ITS2, (b) Annotation of *C. gariepinus* ITS2, (c) Annotation of *H. fossilis* ITS2, (d) Annotation of *S. seenghala* ITS2, and (e) Annotation of *W. attu* ITS2.

(a)	Annotation of *C. batrachus* ITS2. Cb32 and Cb35 representing the individuals of *C. batrachus.*

| **>Cb32** | | | | | |
| --- | --- | --- | --- | --- | --- |
| 1 | 47 | 48 | 371 | 372 | 385 |
| ATTAAATT ... GTCCCTCC  TGGGGCTACGTCTGTCTGACTG | | 5’  CTATTCCCATCGATCAGACTTCGGTCCGTGGCTGTCGGTGGTTCCCTTCCACTTCGACTGGCCGCTTCACACCTCTGCAGGGCCGGTGGTCGCGCGAAAAGCTAGTGGGTGCGGTTTGGCGCGTGTGAACGTGTGGGCGGGGAACAGCTGGGGGTCTGGCGGTGCTCTCTCCGAGACGCTTTTCACTTCTCTCTCTGCTGCCTCTATCTCCCCCGAACTTCTCTGCGGAGAGACGTGTGGGGAGACGAGGGAGGAGAGGGGTCAGGGAGGCTCACCTTTCTCTCGAAAGATGCATCCGATCACAACTTCAGATTAAACGAGACG 3’ | | ACC  CGCTGAATTAA | |
| **5.8S motif** | | **ITS2** | | **28S motif** | |
| **>Cb35** | | | | | |
| 1 | 43 | 44 | 354 | 355 | 370 |
| ATTGNNGG ... GTCCCTCC  TGGGGCTACGTCTGTCTGACTG | | 5’  CTATTCCCATCGATCAGACTTCGGTCCGTGGCTGTCGGTGGTTCCCTTCCACTTCGACTGGCCGCTTCACACCTCTGCAGGGCCGGTGGTCGCGCGAAAAGCTAGTGGGTGCGGTTTGGCGCGTGTGAACCTGTGGGCGGGGAACAGCTGGGGGTCTCCGAGACGCTTTTCACTTCTCTCTCTGCTGCCTCTATCTCCCCCGAACTTCTCTGCGGAGAGACGTGTGGGGAGACGAGGGAGGAGAGGGGTCAGGGAGGCTCACCTTTCTCTCGAAAGATGCATCCGATCACAACTTCAGATTAAACGAGACG 3’ | | ACC  CGCTGAATTAAGT | |
| **5.8S motif** | | **ITS2** | | **28S motif** | |

(b)	Annotation of *C. gariepinus* ITS2. Cg07 and Cg11 are representing the individuals of *C. gariepinus*

| **>Cg07** | | | | | |
| --- | --- | --- | --- | --- | --- |
| 1 | 49 | 50 | 507 | 508 | 544 |
| ATTAAATT ... GGTCCTCC  CGGGGCTACGCCTGTCTGAGGGTCG | | 5’ CTATTCCCATCTATCGGACTTCGGTCCGCGGCTGGAGGTTCTCGCAGGCCCTTAGGGGCCTTCGTCCTCCTAAGTGCAGACTCGTCTTTCCCCTGGCGCCTCGTGTCGCTCTCCCCTCCGTGTCCACGACTCAAAACCCCTCCGGCCGGAAACGGTTTCCGTGCCCTGCGGTCGTCGTGCGCGGCTGTCGGTGGTTCCCTTCCACTTCGACTGCCCGCGTCACGCCTCCGCAGGCCGGGGCCGCGCGAAAAGCCGGTGGGGCAGTGGGCGCAGTGCAGGCGGTCGGCGGGCGGGGAGGCCGGGGGTCGGACGGCGCTCTCCGAGACGCCCTTCTCCCTTCCTCATCACTCCCCCCGCTCTCTCCGAGGAGAGAGTGGAAGGGAGCGAGGGAGAGAGGGGGGGCTCACCTTCTCCTCCTCCTCCACCTCCGTGGAGAGGGGAGAACACACATCCGAACG 3’ | | CGACCTCAGATCAGACGAGACGACC  CGACTGAATTAA | |
| **5.8S motif** | | **ITS2** | | **28S motif** | |

| **>Cg11** | | | | | |
| --- | --- | --- | --- | --- | --- |
| 1 | 50 | 51 | 508 | 509 | 546 |
| AATANATT ... GTCCCTCC  CGGGGCTACGCCTGTCTGAGGGTCG | | 5’  CTATTCCCATCTATCGGACTTCGGTCCGCGGCTGGAGGTTCTCGCAGGCCCTTAGGGGCCTTCGTCCTCCTAAGTGCAGACTCGTCTTTCCCCTGGCGCCTCGTGTCGCTCTCCCCTCCGTGTCCACGACTCAAAACCCCTCCGGCCGGAAACGGTTTCCGTGCCCTGCGGTCGTCGTGCGCGGCTGTCGGTGGTTCCCTTCCACTTCGACTGCCCGCGTCACGCCTCCGCAGGCCGGGGCCGCGCGAAAAGCCGGTGGGGCAGTGGGCGCAGTGCAGGCGGTCGGCGGGCGGGGAGGCCGGGGGTCGGACGGCGCTCTCCGAGACGCCCTTCTCCCTTCCTCATCACTCCCCCCGCTCTCTCTGAGGAGAGAGTGGAAGGGAGCGAGGGAGAGAGGGGGGGCTCACCTTCTCCTCCTCCTCCACCTCCGTGGAAAAGGGAAAACACACATCCGAACG 3’ | | CGACCTCAGATCAGAAGAGACGACT  AACTGAATTAATT | |
| **5.8S motif** | | **ITS2** | | **28S motif** | |

(c)	Annotation of *H. fossilis* ITS2. Hf40, Hf41, Hf42 & Hf44 are representing the multiple individuals of *H. fossilis.*

| **>Hf40** | | | | | |
| --- | --- | --- | --- | --- | --- |
| 1 | 51 | 52 | 417 | 418 | 460 |
| ATTANATT ... GTCCCTCC  CGGGGCTACGCCTGTCTGAGGGTCG | | CTATTCTCATCGATCGGACTTCGGTCCGCGACTGGAGATTCGCAGGCACCCGCCTTCGTCCTCCCAAACGCAGACTCGTTTTTCCCTTCCACCCCGTATTGTTCTCCCCTCCGCGTCCACGACTCAAAACCCCCGGCCGGAAACGGTTCCGTGCCCTGCGGTCGTCGTGCGCGGCTGTCGGTGGGTCCCTTCCACTTAGACTGCCCGCGTCACGACTCCGCAGGCCTGGGCCGCGTGAGAAGCCGGTGGGTGTGGTCGCGCGTAGCGGGAGAACGGCGGGTAGGCGGCGGGGAGAGGCGGCGCTTTAGGCCCGGGCCGCCGCGTGGCGCCCCACTGCCGAAGCGTTAAGCTTCGGAAATCCGAACG | | CGACCTCAGATCAGACGAGACGACC  CGCTGAAT ... GGAGAAAG | |
| **5.8S motif** | | **ITS2** | | **28S motif** | |
| **>Hf41** | | | | | |
| 1 | 51 | 52 | 417 | 418 | 455 |
| TTTACATT ... GTCCCTCC  CGGGGCTACGCCTGTCTGAGGGTCG | | CTATTCTCATCGATCGGACTTCGGTCCGCGACTGGAGATTCGCAGGCACCCGCCTTCGTCCTCCCAAACGCAGACTCGTTTTTCCCTTCCACCCCGTATTGGTCTCCCCTCCGCGTCCACGACTCAAAACCCCCGGGCGGAAACGGTTCCGTGGCCTGGGGTCGTCGTGCGCGGCTGGCGGTGGGTCCCTTCCACTTAGACTGGCCGCGTCACGACTCCGCAGGGCTGGGCCGCGTGAGAAGCCGGTGGGTGTGGTCGCGCGTAGCGGGAGAACCGCGGGTAAGCGGCGGGGAAAAGCGGCGCTTTAAGGCCGGGCCGCCGCGTGGCGCCCCACTGGCGAAGCGTTAAGCTTCGGAAATCCGAACG | | CGACCTCCGATCAGACGAGACGACC  CGGCTGAATTAAA | |
| **5.8S motif** | | **ITS2** | | **28S motif** | |
| **>Hf42** | | | | | |
| 1 | 50 | 51 | 417 | 418 | 454 |
| TTNCATTG ... GTCCCTCC  CGGGGCTACGCCTGTCTGAGGGTCG | | CTATTCTCATCGATCGGACTTCGGTCCGCGACTGGAGATTCGCAGGCACCCGCCTTCGTCCTCCCAAACGCAGACTCGTTTTTCCCTTCCACCCCGTATTGTTCTCCCCTCCGCGTCCACGACTCAAAACCCCCCGGCCGGAAACGGTTCCGTGCCCTGCGGTCGTCGTGCGCGGCTGTCGGTGGGTCCCTTCCACTTAGACTGCCCGCGTCACGACTCCGCAGGCCTGGGCCGCGTGAGAAGCCGGTGGGTGTGGTCGCGCGTAGCGGGAGAACGGCGGGTAGGCGGCGGGGAGAGGCGGCGCTTTAGGCCCGGGCCGCCGCGTGGCGCCCCACTGCCGAAGCGTTAAGCTTCGGAAATCCGAACG | | CGACCTCAGATCAGACGAGACGACC  CGGCTGAATTAA | |
| **5.8S motif** | | **ITS2** | | **28S motif** | |

| **>Hf44** | | | | | |
| --- | --- | --- | --- | --- | --- |
| 1 | 50 | 51 | 416 | 417 | 449 |
| TTTANATT ... GGTCCTCC  CGGGGCTACGCCTGTCTGAGGGTCG | | CTATTCTCATCGATCGGACTTCGGTCCGCGACTGGAGATTCGCAGGCACCCGCCTTCGTCCTCCCAAACGCAGACTCGTTTTTCCCTTCCACCCCGTATTGTTCTCCCCTCCGCGTCCACGACTCAAAACCCCCGGCCGGAAACGGTTCCGTGCCCTGCGGTCGTCGTGCGCGGCTGTCGGTGGGTCCCTTCCACTTAGACTGCCCGCGTCACGACTCCGCAGGCCTGGGCCGCGTGAGAAGCCGGTGGGTGTGGTCGCGCGTAGCGGGAGAACGGCGGGTAGGCGGCGGGGAGAGGCGGCGCTTTAGGCCCGGGCCGCCGCGTGGCGCCCCACTGCCGAAGCGTTAAGCTTCGGAAATCCGAACG | | CGACCTCAGATCAGACGAGACGACC  GGGCTGAA | |
| **5.8S motif** | | **ITS2** | | **28S motif** | |

(d)	Annotation of *S. seenghala* ITS2. Ss01, Ss08, Ss09 & Ss11 are representing the multiple individuals of *S. seenghala.*

| **>Ss01** | | | | | |
| --- | --- | --- | --- | --- | --- |
| 1 | 51 | 52 | 379 | 380 | 417 |
| TTTAAATT ... GTCCCTCC  CGGGGCTACGCCTGTCTGAGGGTCG | | CTATTACCGTCGATCGGACGCCCCCGCGTTCCGCGGCTGGAGGTTTTCTTCGCAGGCGGGCCACCCCCCGCCTCCGTCCTCCCAAAAGCAGACGCGCCCCTCTCTCTCTCTCTCTCTCCCCGCGTCGCCGCCTCGCCCAGTTCGGCGCGAGATAAAAAGAAAGCGGCTGCCGGTGGGTCCTTCCACTCTGGCTGCCCGCTTCGAAAAAAATAGCGCGCCGAGCGACGGGGCACCCACGCGCGACGTCGTGGATGGGGGAGTGAAAAAAGGAGGTGGGGCGCACGCACCTCTCTCTCTCCCTCACACGTCTCTAAAAAAAAAAAACGCG | | CGCCCTCACATCACACGAGACCCCC  CGCTATTATATAT | |
| **5.8S motif** | | **ITS2** | | **28S motif** | |
| **>Ss08** | | | | | |
| 1 | 50 | 51 | 371 | 372 | 408 |
| TTAANTTG ... GTCCCTCC  CGGGGCTACGCCTGTCTGAGGGTCG | | CTATTACCGTCGATCGGACGCCCCCGCGTTCCGCGGCTGGAGGTTTTCTTCGCAGGCGGGCCACCCCCCGCCTCCGTCCTCCCAAAAGCAGACGCGCCCCTCTCTCTCTCCCTCCCCGCGTCGCCGCCTCGCCCAGTTCGGCGCGAGATAAAAAGAAAGCGGCTGCCGGTGGGTCCTTCCACTCTGGCTGCCCGCTTCGAAAAATATAGCGCGCCGAGCGACGGGGCACCCACGCGCGACGTCGTGGATGGGGGAGTGAAAAAGGGAGGTGGGGCGCACGCACCTCTCTCGACCTTACACGTCTCTAAAAAAAAACGAACG | | CGACCTCAGATCAGACGAGACGACC  CGCTGAATTAAA | |
| **5.8S motif** | | **ITS2** | | **28S motif** | |
| **>Ss09** | | | | | |
| 1 | 51 | 52 | 382 | 383 | 418 |
| ATTNAATT ... GTCCCTCC  CGGGGCTACGCCTGTCTGAGGGTCG | | CTATTACCGTCGATCGGACGCCCCCGCGTTCCGCGGCTGGAGGTTTTCTTCGCAGGCGGGCCACCCCCCGCCTCCGTCCTCCCAAAAGCAGACGCGCCCCTCTCTCTCTCTCTCCCTCCCTCCCCGCGTCGCCGCCTCGCCCAGTTCGGCGCGAGATAAAAAGAAAGCGGCTGCCGGTGGGTCCTTCCACTCTGGCTGCCCGCTTCGAAAAATATAGCGCGCCGAGCGACGGGGCACCCACGCGCGACGTCGTGGATGGGGGAGTGAAAAAAAGGAGGTGGGGCGCACGCACCTCTCTCGACCTTACACGTCTCTAAAAAAAAAACGAACG | | CGACCTCAGATCAGACGAGACGACC  CGCTGAATTAG | |
| **5.8S motif** | | **ITS2** | | **28S motif** | |
| **>Ss11** | | | | | |
| 1 | 51 | 52 | 372 | 373 | 409 |
| TTTNCATT ... GTCCCTCC  CGGGGCTACGCCTGTCTGAGGGTCG | | CTATTACCGTCGATCGGACGCCCCCGCGTTCCGCGGCTGGAGGTTTTCTTCGCAGGCGGGCCACCCCCCGCCTCCGTCCTCCCAAAAGCAGACGCGCCCCTCTCTCTCTCCCTCCCCGCGTCGCCGCCTCGCCCAGTTCGGCGCGAGATAAAAAGAAAGCGGCTGCCGGTGGGTCCTTCCACTCTGGCTGCCCGCTTCGAAAAATATAGCGCGCCGAGCGACGGGGCACCCACGCGCGACGTCGTGGATGGGGGAGTGAAAAAGGGAGGTGGGGCGCACGCACCTCTCTCGACCTTACACGTCTCTAAAAAAAAACGAACG | | CGACCTCAGATCAGACGAGACGACC  GGCTGAATTAAA | |
| **5.8S motif** | | **ITS2** | | **28S motif** | |

(e)	Annotation of *W. attu* ITS2. Wa42, Wa32 & Wa51 are representing the multiple individuals of *W. attu.*

| **>Wa42** | | | | | |
| --- | --- | --- | --- | --- | --- |
| 1 | 50 | 51 | 463 | 464 | 502 |
| TTAAATTG ... GTCCCTCC  CGGGGCTACGCCTGTCTGAGGGTCG | | CTATTACCATCGATCGTACGAAACGCGCGCGACTGGAAGCTCGCAGGCCGGCTGACGGCCTTCGTCCTCCTAAAAGCAGACGCTCCGTTTCTCCTTTCCATGCCTTTTTTTTTTTCTCCCAAAAAAGAAACCCCTCCCGAACTCGGCGCCTTTCCCCCAAACCATCCCTTTGGGGCGGGGGGACGCGCGAGCGGCTGCCGGTGGTGTCTCGCACTCTGGCTGCCCGCGAGCGCGCGTTCCCCGGAAACCCCGAGGGCGGGGAAGGGACAGAGGAAGGGCGCGCCACGGGCGAAAGAGGGGGGCGGGAAAAAAGAGAAAAAAAGAGAGAGAGAGACGGGGAGGGGGGCTAAGAAAAGAGGGAGAACCACTCCCTCGTTCCGTTGCCCTCTTATCTATTTCCTTCCTTTCGAACG | | CGACCTCAGATCAGACGAGACGACC  CGCTGAATTAAAAA | |
| **5.8S motif** | | **ITS2** | | **28S motif** | |
| **>Wa32** | | | | | |
| 1 | 50 | 51 | 465 | 466 | 502 |
| TTAAATTG ... GTCCCTCC  CGGGGCTACGCCTGTCTGAGGGTCG | | CTATTACCATCGATCGTACGAAACGCGCGCGACTGGAAGCTCGCAGGCCGGCTGACGGCCTTCGTCCTCCTAAAAGCAGACGCTCCGTTTCTCCTTTCCATGCCTTTTTTTTTTTCTCCCAAAAAAGAAACCCCTCCCGAACTCGGCGCCTTTCCCCCAAACCATCCCTTTGGGGCGGGGGGACGCGCGAGCGGCTGCCGGTGGTGTCTCGCACTCTGGCTGCCCGCGAGCGCGCGTTCCCCGGAAACCCCGAGGGCGGGGAAGGGACAGAGGAAGGGCGCGCCACGGGCGAAAGAGGGGGGCGGGAAAAAAGAGAAAAAAAGAGAGAGAGAGAGACGGGGGGGGGGGCTAAAAAAAGAGGGAAACCCACTCCCTCGTTCCGTTGCCCTCTTATCTATTCCCTTCTTTTCAAACG | | CGACCTCAGATCAGAAGAGACCACC  CGCTGAAATAAA | |
| **5.8S motif** | | **ITS2** | | **28S motif** | |
| **>Wa51** | | | | | |
| 1 | 50 | 51 | 465 | 466 | 503 |
| TTAAATTG ... GTCCCTCC  CGGGGCTACGCCTGTCTGAGGGTCG | | CTATTACCATCGATCGTACGAAACGCGCGCGACTGGAAGCTCGCAGGCCGGCTGACGGCCTTCGTCCTCCTAAAAGCAGACGCTCCGTTTCTCCTTTCCATGCCTTTTTTTTTTTCTCCCAAAAAAGAAACCCCTCCCGAACTCGGCGCCTTTCCCCCAAACCATCCCTTTGGGGCGGGGGGACGCGCGAGCGGCTGCCGGTGGGGTCTCGCACTCTGGCTGCCCGCGAGCGCGCGTTCCCCGGAAACCCCGAGGGCGGGGAAGGGACAGAGGAAGGGCGCGCCACGGGCGAAAGAGGGGGGCGGGAAAAAAGAGAAAAAAAGAGAGAGAGAGAGACGGGGAGGGGGGCTAAAAAAAGAGGGAAAACCACCCCCTCGTTCCGGTGCCCTCTTATCTATTTCCTTCCTTTCAAACG | | CCCCCTCAGATCAAAAGAGACCACC  CGCTGAAATAAAA | |
| **5.8S motif** | | **ITS2** | | **28S motif** | |

- **Secondary structure Prediction**

The Predict option of ITS2 database performs the secondary structure prediction analysis. It works on the basis of homology modelling method where the secondary structure of any ITS2 sequence is predicted by the homology modelling with the complete set of sequences and structures of the database serving as template. The annotated ITS2 sequences of all catfish species generated by the annotation tool were used for the structure prediction. The server generates the secondary structure alignments in xfasta file format for further analysis.

- **Multiple Alignment of Secondary Structure**

ITS2 database offers multiple alignment option both for sequences and their secondary structures. Using this tool, the multiple alignment of all the catfish species was done and scanned for the nucleotide sequences which are conserved in these species.

- **Conserved Motif Search**

The Annotated ITS2 sequences were analyzed for the conserved sequence motifs. These motifs are the hallmark of ITS2 secondary structure in eukaryotes (Coleman 2003, Schultz et al. 2005). This sequence based tool locates the position and annotate these eukaryotic conserved sequence motifs in the dataset (Table B).

**Table (B):** Conserved Motif search of the ITS2 sequences. The motif search output is presented in the fasta format. Multiple individuals are used to avoid any discrepancy in tracing the exact location of pyrimidine-pyrimidine mismatch (U-U/U-C) and UGGU/UGGU motif of ITS2 sequences in all the species. Pyrimidine-Pyrimidine mismatch is shown in green, and UGGU motif is shown in red. (a) Conserved Motif of *C. batrachus* ITS2, (b) Conserved Motif of *C. gariepinus* ITS2, (c) Conserved Motif of *H. fossilis* ITS2, (d) Conserved Motif of *S. seenghala* ITS2, and (e) Conserved Motif of *W. attu* ITS2.

(a)	Conserved Motif of *C. batrachus* ITS2. Cb32 and Cb35 representing the individuals of *C. batrachus.*

| **>Cb32** | | | | | | | | | |
| --- | --- | --- | --- | --- | --- | --- | --- | --- | --- |
| 1 | 47 | 48 | 69 | 70 | 219 | 220 | 239 | 240 | 324 |
| CTATTC...  GTCCGTGGCTGTCGGTGGTTCCCT | | TCCACTTCGACTGGCCGCTTCA | | CACCTCTGCAGGGCCGGTGGTCGCGCGAAAAGCTAGTGGGTGCGGTTTGGCGCGTGTGAACCTGTGGGCGGGGAACAGCTGGGGGTCTGGCGGTGCTCTCTCCGAGACGCTTTTCACTTCTCTCTCTGCTGCCTCTATCTCCCCCGAACTTCTCTGCGGAGAG | | ACGTGTGGGGAGACGAGGG | | AGGAGAGGG... GATTAAACGAGACG | |
|  | | **U-U mismatch** | |  | | **UGGU (III)** | |  | |
| **>Cb35** | | | | | | | | | |
| 1 | 47 | 48 | 69 | 70 | 206 | 207 | 225 | 226 | 311 |
| CTATTC...  GTCCGTGGCTGTCGGTGGTTCCCT | | TCCACTTCGACTGGCCGCTTCA | | CACCTCTGCAGGGCCGGTGGTCGCGCGAAAAGCTAGTGGGTGCGGTTTGGCGCGTGTGAACCTGTGGGCGGGGAACAGCTGGGGTCTCCGAGACGCTTTTCACTTCTCTCTCTGCTGCCTCTATCTCCCCCGAACTTCTCTGCGGAGAG | | ACGTGTGGGGAGACGAGGG | | AGGAGAGGG...  TTAAACGAGACG | |
|  | | **U-U mismatch** | |  | | **UGGU (III)** | |  | |

(b)	Conserved Motif of *C. gariepinus* ITS2. Cg07 and Cg11 are representing the individuals of *C. gariepinus.*

| **>Cg07** | | | | | | | | | | |
| --- | --- | --- | --- | --- | --- | --- | --- | --- | --- | --- |
| 1 | 50 | 51 | 77 | 78 | | 182 | 183 | 197 | 198 | 458 |
| CTATTCC...  ACTTCGGTCCGCGGCTGGAGGTTCTCGCAGGCC | | CTTAGGGGCCTTCGTCCTCCTAAGTGC | | AGACTCGTCTTTCCCCTGGCGCCTCGTGTCGCTCTCCCCTCCGTGTCCACGACTCAAAACCCCTCCGGCCGGAAACGGTTTCCGTGCCCTGCGGTCGTCGTGCGC | | | GGCTGTCGGTGGTTC | | CCTTCCAC ... TCCGAACG | |
|  | | **U-U mismatch** | |  | | | **UGGU (III)** | |  | |
| **>Cg11** | | | | | | | | | | |
| 1 | 50 | 51 | 77 | | 78 | 182 | 183 | 197 | 198 | 458 |
| CTATTCC...  ACTTCGGTCCGCGGCTGGAGGTTCTCGCAGGCC | | CTTAGGGGCCTTCGTCCTCCTAAGTGC | | | AGACTCGTCTTTCCCCTGGCGCCTCGTGTCGCTCTCCCCTCCGTGTCCACGACTCAAAACCCCTCCGGCCGGAAACGGTTTCCGTGCCCTGCGGTCGTCGTGCGC | | GGCTGTCGGTGGTTC | | CCTTCCAC ... TCCGAACG | |
|  | | **U-U mismatch** | | |  | | **UGGU (III)** | |  | |

(c)	Conserved Motif of *H. fossilis* ITS2. Hf40, Hf41, Hf42 & Hf44 are representing the multiple individuals of *H. fossilis.*

| **>Hf40** | | | | | | | | | | |
| --- | --- | --- | --- | --- | --- | --- | --- | --- | --- | --- |
| 1 | 40 | 41 | 52 | 53 | | 172 | 173 | 187 | 188 | 366 |
| CTATTCT...  CTGGAGATT | | CGCAGGCACCCG | | CCTTCGTCCTCCCAAACGCAGACTCGTTTTTCCCTTCCACCCCGTATTGTTCTCCCCTCCGCGTCCACGACTCAAAACCCCCGGCCGGAAACGGTTCCGTGCCCTGCGGTCGTCGTGCGC | | | GGCTGTCGGTGGGTC | | CCTTCCAC ... TCCGAACG | |
|  | | **U-C mismatch** | |  | | | **UGGU (III)** | |  | |
| **>Hf41** | | | | | | | | | | |
| 1 | 40 | 41 | 52 | 53 | | 172 | 173 | 187 | 188 | 366 |
| CTATTCT...  CTGGAGATT | | CGCAGGCACCCG | | CCTTCGTCCTCCCAAACGCAGACTCGTTTTTCCCTTCCACCCCGTATTGTTCTCCCCTCCGCGTCCACGACTCAAAACCCCCGGCCGGAAACGGTTCCGTGCCCTGCGGTCGTCGTGCGC | | | GGCTGTCGGTGGGTC | | CCTTCCAC ... TCCGAACG | |
|  | | **U-C mismatch** | |  | | | **UGGU (III)** | |  | |
| **>Hf42** | | | | | | | | | | |
| 1 | 40 | 41 | 52 | | 53 | 172 | 173 | 187 | 188 | 367 |
| CTATTCT...  CTGGAGATT | | CGCAGGCACCCG | | CCTTCGTCCTCCCAAACGCAGACTCGTTTTTCCCTTCCACCCCGTATTGTTCTCCCCTCCGCGTCCACGACTCAAAACCCCCGGCCGGAAACGGTTCCGTGCCCTGCGGTCGTCGTGCGC | | | GGCTGTCGGTGGGTC | | CCTTCCAC ... TCCGAACG | |
|  | | **U-C mismatch** | |  | | | **UGGU (III)** | |  | |

| **>Hf44** | | | | | | | | | |
| --- | --- | --- | --- | --- | --- | --- | --- | --- | --- |
| 1 | 40 | 41 | 52 | 53 | 172 | 173 | 187 | 188 | 366 |
| CTATTCT...  CTGGAGATT | | CGCAGGCACCCG | | CCTTCGTCCTCCCAAACGCAGACTCGTTTTTCCCTTCCACCCCGTATTGTTCTCCCCTCCGCGTCCACGACTCAAAACCCCCGGCCGGAAACGGTTCCGTGCCCTGCGGTCGTCGTGCGC | | GGCTGTCGGTGGGTC | | CCTTCCAC ... TCCGAACG | |
|  | | **U-C mismatch** | |  | | **UGGU (III)** | |  | |

(d)	Conserved Motif of *S. seenghala* ITS2. Ss01, Ss08, Ss09 & Ss11 are representing the multiple individuals of *S. seenghala.*

| **>Ss01** | | | | | | | | | |
| --- | --- | --- | --- | --- | --- | --- | --- | --- | --- |
| 1 | 50 | 51 | 65 | 66 | 168 | 169 | 186 | 187 | 328 |
| CTATTACC ... AGGTTTTCTT | | CGCAGGCGGGCCACC | | CCCCGCCTCCGTCCTCCCAAAAGCAGACGCGCCCCTCTCTCTCTCTCTCTCTCCCCGCGTCGCCGCCTCGCCCAGTTCGGCGCGAGATAAAAAGAAAGCGGCT | | GCCGGTGGGTCCTTCCAC | | CTTCCACT ... AAAACGCG | |
|  | | **U-U mismatch** | |  | | **UGGU (III)** | |  | |
| **>Ss08** | | | | | | | | | |
| 1 | 50 | 51 | 65 | 66 | 164 | 165 | 182 | 173 | 321 |
| CTATTACC ... AGGTTTTCTT | | CGCAGGCGGGCCACC | | CCCCGCCTCCGTCCTCCCAAAAGCAGACGCGCCCCTCTCTCTCTCCCTCCCCGCGTCGCCGCCTCGCCCAGTTCGGCGCGAGATAAAAAGAAAGCGGCT | | GCCGGTGGGTCCTTCCAC | | CTTCCACT ... AACGAACG | |
|  | | **U-U mismatch** | |  | | **UGGU (III)** | |  | |
| **>Ss09** | | | | | | | | | |
| 1 | 50 | 51 | 65 | 66 | 172 | 173 | 190 | 191 | 331 |
| CTATTACC ... AGGTTTTCTT | | CGCAGGCGGGCCACC | | CCCCGCCTCCGTCCTCCCAAAAGCAGACGCGCCCCTCTCTCTCTCTCTCCCTCCCTCCCCGCGTCGCCGCCTCGCCCAGTTCGGCGCGAGATAAAAAGAAAGCGGCT | | GCCGGTGGGTCCTTCCAC | | CTTCCACT ... AACGAACG | |
|  | | **U-U mismatch** | |  | | **UGGU (III)** | |  | |
| **>Ss11** | | | | | | | | | |
| 1 | 50 | 51 | 65 | 66 | 164 | 165 | 182 | 173 | 321 |
| CTATTACC ... AGGTTTTCTT | | CGCAGGCGGGCCACC | | CCCCGCCTCCGTCCTCCCAAAAGCAGACGCGCCCCTCTCTCTCTCCCTCCCCGCGTCGCCGCCTCGCCCAGTTCGGCGCGAGATAAAAAGAAAGCGGCT | | GCCGGTGGGTCCTTCCAC | | CTTCCACT ... AACGAACG | |
|  | | **U-U mismatch** | |  | | **UGGU (III)** | |  | |

(e)	Conserved Motif of *W. attu* ITS2. Wa42, Wa32 & Wa51 are representing the multiple individuals of *W. attu.*

| **>Wa42** | | | | | | | | | | |
| --- | --- | --- | --- | --- | --- | --- | --- | --- | --- | --- |
| 1 | 40 | 41 | 55 | 56 | 192 | | 193 | 207 | 208 | 413 |
| CTATTACC ...  CTGGAAGC | | TCGCAGGCCGGCTGA | | CGGCCTTCGTCCTCCTAAAAGCAGACGCTCCGTTTCTCCTTTCCATGCCTTTTTTTTTTTCTCCCAAAAAAGAAACCCCTCCCGAACTCGGCGCCTTTCCCCCAAACCATCCCTTTGGGGCGGGGGGACGCGCGAGC | | | GGCTGCCGGTGGTGT | | CTCGCACT ... TTCGAACG | |
|  | | **U-U mismatch** | |  | | | **UGGU (III)** | |  | |
| **>Wa32** | | | | | | | | | | |
| 1 | 40 | 41 | 55 | 56 | | 192 | 193 | 207 | 208 | 415 |
| CTATTACC ...  CTGGAAGC | | TCGCAGGCCGGCTGA | | CGGCCTTCGTCCTCCTAAAAGCAGACGCTCCGTTTCTCCTTTCCATGCCTTTTTTTTTTTCTCCCAAAAAAGAAACCCCTCCCGAACTCGGCGCCTTTCCCCCAAACCATCCCTTTGGGGCGGGGGGACGCGCGAGC | | | GGCTGCCGGTGGTGT | | CTCGCACT ... TTCGAACG | |
|  | | **U-U mismatch** | |  | | | **UGGU (III)** | |  | |
| **>Wa51** | | | | | | | | | | |
| 1 | 40 | 41 | 55 | 56 | | 192 | 193 | 207 | 208 | 415 |
| CTATTACC ...  CTGGAAGC | | TCGCAGGCCGGCTGA | | CGGCCTTCGTCCTCCTAAAAGCAGACGCTCCGTTTCTCCTTTCCATGCCTTTTTTTTTTTCTCCCAAAAAAGAAACCCCTCCCGAACTCGGCGCCTTTCCCCCAAACCATCCCTTTGGGGCGGGGGGACGCGCGAGC | | | GGCTGCCGGTGGTGT | | CTCGCACT ... TTCGAACG | |
|  | | **U-U mismatch** | |  | | | **UGGU (III)** | |  | |

***Secondary Structure Annotation ITS2***

- **4Sale based analysis**

The 4Sale software (Seibel et al. 2006; 2008) which is Java based program was used for the analysis of the secondary structures generated by the aforesaid servers. The multiple secondary structure alignment for each catfish species was observed individually. This software includes an interactive secondary structure viewer which provides a self explanatory and interactive secondary structure with complete information on nucleotide positions and structural features.

- **VARNA based analysis**

VARNA is again a Java based Graphic User Interface (GUI) tool (Darty et al. 2009) which was used for the automatic drawing, interactive annotation and visualization of the diverse features in the secondary structures of ITS2 which were traced by the 4Sale software. It provides deeper insight into the secondary structure with numerous customized structure specific annotating options to further boost the structural properties graphically for an enhanced structural understanding. All the eukaryotic conserved hallmarks, catfish conserved sequences, microsatellite repeats, random repeats, non-canonical base-pairs and other basic structural defining feature were annotated on all the ITS2 sequences by using VARNA tool.

**Secondary Structure Analysis ITS1**

The ITS1 secondary structure prediction was also done separately for all the individuals of each catfish species by using RNAfold web server using minimum free energy (MFE) method (<http://rna.tbi.univie.ac.at/cgi-bin/RNAWebSuite/RNAfold.cgi>) (Gruber et al. 2008). The structural alignments were saved as xfasta file format for further analysis. The LocARNA web server (<https://rna.informatik.uni-freiburg.de>) (Will et al. 2007, Will et al. 2012, Raden et al. 2018) was used for the multiple sequence alignment of ITS1 sequences on the basis of their secondary structure. The Annotation of Secondary Structures of ITS1 was done in a similar way to that of ITS2 using softwares 4Sale and VARNA.
