## Supplementary material for "Exploring unique sequence repeat patterns and secondary structures in rDNA internal transcribed spacers ITS1 and ITS2 for characterization of catfish species": Supplementary Data S3.pdf

Cb: *Clarias batrachus*

```
                                                          5.8s rRNA  ITS2
Cb32 ATTAAATTGCGGCCCGAGTCCCTCCTGGGGCTACGTCTGTCTGACTGCTATTCCCATCGATCAGACTTCG 70
Cb35 ----ATTGNNGGCCCGAGTCCCTCCTGGGGCTACGTCTGTCTGACTGCTATTCCCATCGATCAGACTTCG 70
         * *   ************************************************************

Cb32 GTCCGTGGCTGTCGGTGGTTCCCTTCCACTTCGACTGGCCGCTTCACACCTCTGCAGGGCCGGTGGTCGC 140
Cb35 GTCCGTGGCTGTCGGTGGTTCCCTTCCACTTCGACTGGCCGCTTCACACCTCTGCAGGGCCGGTGGTCGC 140
     **********************************************************************

Cb32 GCGAAAAGCTAGTGGGTGCGGTTTGGCGCGTGTGAACGTGTGGGCGGGGAACAGCTGGGGGTCTGGCGGT 210
Cb35 GCGAAAAGCTAGTGGGTGCGGTTTGGCGCGTGTGAACCTGTGGGCGGGGAACAGCTGGGGGTC------- 210
     ************************************* *************************

                                             Cb2-1
Cb32 GCTCTCTCCGAGACGCTTTTCACTTCTCTCTCTGCTGCCTCTATCTCCCCCGAACTTCTCTGCGGAGAGA 280
Cb35 ------TCCGAGACGCTTTTCACTTCTCTCTCTGCTGCCTCTATCTCCCCCGAACTTCTCTGCGGAGAGA 280
           ****************************************************************

                      Cb2-A
Cb32 CGTGTGGGGAGACGAGGGAGGAGAGGGGTCAGGGAGGCTCACCTTTCTCTCGAAAGATGCATCCGATCAC 350
Cb35 CGTGTGGGGAGACGAGGGAGGAGAGGGGTCAGGGAGGCTCACCTTTCTCTCGAAAGATGCATCCGATCAC 350
     **********************************************************************
                             28s rRNA
Cb32 AACTTCAGATTAAACGAGACGACCCGCTGAATT 383
Cb35 AACTTCAGATTAAACGAGACGACCCGCTGAATT 383
     *********************************
```

Microsatellite repeats: Cb2-1

Random repeats: Cb2-A

**Figure S3-(1).** Multiple sequence alignment of *C. batrachus* ITS2 (Cb2). The highlights are showing the position of microsatellite repeat Cb2-1 and random repeat Cb2-A.

Cg: *Clarias gariepinus*

```
                                                5.8s rRNA  ITS2
Cg07 ATTAAATTGCGGCCCGGGTCC-TCCCGGGGCTACGCCTGTCTGAGGGTCGCTATTCCCATCTATCGGACT 70
Cg11 AATANATTGCGGCCCGGGTCCCTCCCGGGGCTACGCCTGTCTGAGGGTCGCTATTCCCATCTATCGGACT 70
     * ** **************** ************************************************

Cg07 TCGGTCCGCGGCTGGAGGTTCTCGCAGGCCCTTAGGGGCCTTCGTCCTCCTAAGTGCAGACTCGTCTTTC 140
Cg11 TCGGTCCGCGGCTGGAGGTTCTCGCAGGCCCTTAGGGGCCTTCGTCCTCCTAAGTGCAGACTCGTCTTTC 140
     **********************************************************************

Cg07 CCCTGGCGCCTCGTGTCGCTCTCCCCTCCGTGTCCACGACTCAAAACCCCTCCGGCCGGAAACGGTTTCC 210
Cg11 CCCTGGCGCCTCGTGTCGCTCTCCCCTCCGTGTCCACGACTCAAAACCCCTCCGGCCGGAAACGGTTTCC 210
     **********************************************************************

                                       Cg2-A
Cg07 GTGCCCTGCGGTCGTCGTGCGCGGCTGTCGGTGGTTCCCTTCCACTTCGACTGCCCGCGTCACGCCTCCG 280
Cg11 GTGCCCTGCGGTCGTCGTGCGCGGCTGTCGGTGGTTCCCTTCCACTTCGACTGCCCGCGTCACGCCTCCG 280
     **********************************************************************

                                                                Cg2-B
Cg07 CAGGCCGGGGCCGCGCGAAAAGCCGGTGGGGCAGTGGGCGCAGTGCAGGCGGTCGGCGGGCGGGGAGGCC 350
Cg11 CAGGCCGGGGCCGCGCGAAAAGCCGGTGGGGCAGTGGGCGCAGTGCAGGCGGTCGGCGGGCGGGGAGGCC 350
     **********************************************************************

                                    Cg2-C                Cg2-1
Cg07 GGGGGTCGGACGGCGCTCTCCGAGACGCCCTTCTCCCTTCCTCATCACTCCCCCCGCTCTCTCCGAGGAG 420
Cg11 GGGGGTCGGACGGCGCTCTCCGAGACGCCCTTCTCCCTTCCTCATCACTCCCCCCGCTCTCTCTGAGGAG 420
     *************************************************************** ******

                            Cg2-2              Cg2-3                  Cg2-D
Cg07 AGAGTGGAAGGGAGCGAGGGAGAGAGGGGGGGCTCACCTTCTCCTCCTCCTCCACCTCCGTGGAGAGGGG 490
Cg11 AGAGTGGAAGGGAGCGAGGGAGAGAGGGGGGGCTCACCTTCTCCTCCTCCTCCACCTCCGTGGAAAAGGG 490
     *************************************************************** * ***

                            28s rRNA
Cg07 AGAACACACATCCGAACGCGACCTCAGATCAGACGAGACG 530
Cg11 AAAACACACATCCGAACGCGACCTCAGATCAGAAGAGACG 530
     * ******************************* ******
```

Microsatellite repeats: Cg2-1, Cg2-2, Cg2-3

Random repeats: Cg2-A, Cg2-B, Cg2-C, Cg2-D

**Figure S3-(2).** Multiple sequence alignment of *C. gariepinus* ITS2 (Cg2) . The highlights are showing the position of microsatellite repeats Cg2-1 to Cg2-3 and random repeats Cg2-A to Cg2-D.

Hf: *Heteropneustes fossilis*

```
                                                   5.8s rRNA  ITS2
Hf40 ATTANATTGCGGCCCGGGGTCCCTCCCGGGGCTACGCCTGTCTGAGGGTCGCTATTCTCATCGATCGGAC 70
Hf44 TTTANATTGCGGCCCGGGGTCC-TCCCGGGGCTACGCCTGTCTGAGGGTCGCTATTCTCATCGATCGGAC 70
Hf42 -TTNCATTGCGGCCCGGGGTCCCTCCCGGGGCTACGCCTGTCTGAGGGTCGCTATTCTCATCGATCGGAC 70
Hf41 TTTACATTGCGGCCCGGGGTCCCTCCCGGGGCTACGCCTGTCTGAGGGTCGCTATTCTCATCGATCGGAC 70
      **  ***************** ***********************************************

                                            Hf2-A                Hf2-B
Hf40 TTCGGTCCGCGACTGGAGATTCGCAGGCACCCGCCTTCGTCCTCCCAAACGCAGACTCGTTTTTCCCTTC 140
Hf44 TTCGGTCCGCGACTGGAGATTCGCAGGCACCCGCCTTCGTCCTCCCAAACGCAGACTCGTTTTTCCCTTC 140
Hf42 TTCGGTCCGCGACTGGAGATTCGCAGGCACCCGCCTTCGTCCTCCCAAACGCAGACTCGTTTTTCCCTTC 140
Hf41 TTCGGTCCGCGACTGGAGATTCGCAGGCACCCGCCTTCGTCCTCCCAAACGCAGACTCGTTTTTCCCTTC 140
     **********************************************************************

                  Hf2-C                       Hf2-D
Hf40 CACCCCGTATTGTTCTCCCCTCCGCGTCCACGACTCAAAACCCCC-GGCCGGAAACGGTTCCGTGCCCTG 210
Hf44 CACCCCGTATTGTTCTCCCCTCCGCGTCCACGACTCAAAACCCCC-GGCCGGAAACGGTTCCGTGCCCTG 210
Hf42 CACCCCGTATTGTTCTCCCCTCCGCGTCCACGACTCAAAACCCCCCGGCCGGAAACGGTTCCGTGCCCTG 210
Hf41 CACCCCGTATTGGTCTCCCCTCCGCGTCCACGACTCAAAACCCCC-GGGCGGAAACGGTTCCGTGGCCTG 210
     ************ ******************************** ** **************** ****

                Hf2-E
Hf40 CGGTCGTCGTGCGCGGCTGTCGGTGGGTCCCTTCCACTTAGACTGCCCGCGTCACGACTCCGCAGGCCTG 280
Hf44 CGGTCGTCGTGCGCGGCTGTCGGTGGGTCCCTTCCACTTAGACTGCCCGCGTCACGACTCCGCAGGCCTG 280
Hf42 CGGTCGTCGTGCGCGGCTGTCGGTGGGTCCCTTCCACTTAGACTGCCCGCGTCACGACTCCGCAGGCCTG 280
Hf41 GGGTCGTCGTGCGCGGCTGGCGGTGGGTCCCTTCCACTTAGACTGGCCGCGTCACGACTCCGCAGGGCTG 280
      ****************** ************************* ******************** ***

                       Hf2-F                                   Hf2-G
Hf40 GGCCGCGTGAGAAGCCGGTGGGTGTGGTCGCGCGTAGCGGGAGAACGGCGGGTAGGCGGCGGGGAGAGGC 350
Hf44 GGCCGCGTGAGAAGCCGGTGGGTGTGGTCGCGCGTAGCGGGAGAACGGCGGGTAGGCGGCGGGGAGAGGC 350
Hf42 GGCCGCGTGAGAAGCCGGTGGGTGTGGTCGCGCGTAGCGGGAGAACGGCGGGTAGGCGGCGGGGAGAGGC 350
Hf41 GGCCGCGTGAGAAGCCGGTGGGTGTGGTCGCGCGTAGCGGGAGAACCGCGGGTAAGCGGCGGGGAAAAGC 350
     ********************************************** ******* ********** * **

                      Hf2-H                                          28s rRNA
Hf40 GGCGCTTTAGGCCCGGGCCGCCGCGTGGCGCCCCACTGCCGAAGCGTTAAGCTTCGGAAATCCGAACGCG 420
Hf44 GGCGCTTTAGGCCCGGGCCGCCGCGTGGCGCCCCACTGCCGAAGCGTTAAGCTTCGGAAATCCGAACGCG 420
Hf42 GGCGCTTTAGGCCCGGGCCGCCGCGTGGCGCCCCACTGCCGAAGCGTTAAGCTTCGGAAATCCGAACGCG 420
Hf41 GGCGCTTTAAGGCCGGGCCGCCGCGTGGCGCCCCACTGGCGAAGCGTTAAGCTTCGGAAATCCGAACGCG 420
     ********* * ************************** *******************************

Hf40 ACCTCAGATCAGACGAGACGACC 443
Hf44 ACCTCAGATCAGACGAGACGACC 443
Hf42 ACCTCAGATCAGACGAGACGACC 443
Hf41 ACCTCCGATCAGACGAGACGACC 443
     ***** *****************
```

Random repeats: Hf2-A, Hf2-B, Hf2-C, Hf2-D, Hf2-E, Hf2-F, Hf2-G, Hf2-H

**Figure S3-(3).** Multiple sequence alignment of *H. fossilis* ITS2 (Hf2). The highlights are showing the position of random repeats Hf2-A to Hf2-H.

Ss: *Sperata seenghala*

```
                                                          5.8s rRNA <-||-> ITS2
Ss08 -TTAANTTGCGGCCCGGGGTCCCTCCCGGGGCTACGCCTGTCTGAGGGTCGCTATTACCGTCGATCGGAC 70
Ss11 TTTNCATTGCGGCCCGGGGTCCCTCCCGGGGCTACGCCTGTCTGAGGGTCGCTATTACCGTCGATCGGAC 70
Ss09 ATTNAATTGCGGCCCGGGGTCCCTCCCGGGGCTACGCCTGTCTGAGGGTCGCTATTACCGTCGATCGGAC 70
Ss01 TTTAAATTGCGGCCCGGGGTCCCTCCCGGGGCTACGCCTGTCTGAGGGTCGCTATTACCGTCGATCGGAC 70
      **   ****************************************************************

                                                                Ss2-A
Ss08 GCCCCCGCGTTCCGCGGCTGGAGGTTTTCTTCGCAGGCGGGCCACCCCCCGCCTCCGTCCTCCCAAAAGC 140
Ss11 GCCCCCGCGTTCCGCGGCTGGAGGTTTTCTTCGCAGGCGGGCCACCCCCCGCCTCCGTCCTCCCAAAAGC 140
Ss09 GCCCCCGCGTTCCGCGGCTGGAGGTTTTCTTCGCAGGCGGGCCACCCCCCGCCTCCGTCCTCCCAAAAGC 140
Ss01 GCCCCCGCGTTCCGCGGCTGGAGGTTTTCTTCGCAGGCGGGCCACCCCCCGCCTCCGTCCTCCCAAAAGC 140
     **********************************************************************

                   Ss2-1
Ss08 AGACGCGCCCCTCTCTCTCTC--------CCTCCCCGCGTCGCCGCCTCGCCCAGTTCGGCGCGAGATAA 210
Ss11 AGACGCGCCCCTCTCTCTCTC--------CCTCCCCGCGTCGCCGCCTCGCCCAGTTCGGCGCGAGATAA 210
Ss09 AGACGCGCCCCTCTCTCTCTCTCTCCCTCCCTCCCCGCGTCGCCGCCTCGCCCAGTTCGGCGCGAGATAA 210
Ss01 AGACGCGCCCCTCTCTCTCTCTCTC----TCTCCCCGCGTCGCCGCCTCGCCCAGTTCGGCGCGAGATAA 210
     *********************         ****************************************

      Ss2-2
Ss08 AAAGAAAGCGGCTGCCGGTGGGTCCTTCCACTCTGGCTGCCCGCTTCGAAAAATATAGCGCGCCGAGCGA 280
Ss11 AAAGAAAGCGGCTGCCGGTGGGTCCTTCCACTCTGGCTGCCCGCTTCGAAAAATATAGCGCGCCGAGCGA 280
Ss09 AAAGAAAGCGGCTGCCGGTGGGTCCTTCCACTCTGGCTGCCCGCTTCGAAAAATATAGCGCGCCGAGCGA 280
Ss01 AAAGAAAGCGGCTGCCGGTGGGTCCTTCCACTCTGGCTGCCCGCTTCGAAAAAAATAGCGCGCCGAGCGA 280
     ***************************************************** ****************

                                              Ss2-3       Ss2-B
Ss08 CGGGGCACCCACGCGCGACGTCGTGGATGGGGGAGTGAAAAAG-GGAGGTGGGGCGCACGCACCTCTCTC 350
Ss11 CGGGGCACCCACGCGCGACGTCGTGGATGGGGGAGTGAAAAAG-GGAGGTGGGGCGCACGCACCTCTCTC 350
Ss09 CGGGGCACCCACGCGCGACGTCGTGGATGGGGGAGTGAAAAAAAGGAGGTGGGGCGCACGCACCTCTCTC 350
Ss01 CGGGGCACCCACGCGCGACGTCGTGGATGGGGGAGTGAAAAAA-GGAGGTGGGGCGCACGCACCTCTCTC 350
     ******************************************  **************************
                                         28s rRNA
                         Ss2-4          |-->
Ss08 GA--CCTTACACGTCTCTAAAAAAAAA-CGAACGCGACCTCAGATCAGACGAGACGACCCGCTGAATT 418
Ss11 GA--CCTTACACGTCTCTAAAAAAAAA-CGAACGCGACCTCAGATCAGACGAGACGACCGGCTGAATT 418
Ss09 GA--CCTTACACGTCTCTAAAAAAAAAACGAACGCGACCTCAGATCAGACGAGACGACCCGCTGAATT 418
Ss01 TCTCCCTCACACGTCTCTAAAAAAAAA                                          380
         *** *******************
```

Microsatellite repeats: Ss2-1, Ss2-2, Ss2-3, Ss2-4

Random repeats: Ss2-A, Ss2-B

**Figure S3-(4).** Multiple sequence alignment of *S. seenghala* ITS2 (Ss2). The highlights are showing the position of microsatellite repeats Ss2-1 to Ss2-4 and random repeats Ss2-A & Ss2-B.

Wa: *Wallago attu*

```
                                                       5.8s rRNA  ITS2
Wa32 TTAAATTGCGGCCCGGGGTCCCTCCCGGGGCTACGCCTGTCTGAGGGTCGCTATTACCATCGATCGTACG 70
Wa51 TTAAATTGCGGCCCGGGGTCCCTCCCGGGGCTACGCCTGTCTGAGGGTCGCTATTACCATCGATCGTACG 70
Wa42 TTAAATTGCGGCCCGGGGTCCCTCCCGGGGCTACGCCTGTCTGAGGGTCGCTATTACCATCGATCGTACG 70
     **********************************************************************

Wa32 AAACGCGCGCGACTGGAAGCTCGCAGGCCGGCTGACGGCCTTCGTCCTCCTAAAAGCAGACGCTCCGTTT 140
Wa51 AAACGCGCGCGACTGGAAGCTCGCAGGCCGGCTGACGGCCTTCGTCCTCCTAAAAGCAGACGCTCCGTTT 140
Wa42 AAACGCGCGCGACTGGAAGCTCGCAGGCCGGCTGACGGCCTTCGTCCTCCTAAAAGCAGACGCTCCGTTT 140
     **********************************************************************

                  Wa2-1              Wa2-2         Wa2-A                 Wa2-B
Wa32 CTCCTTTCCATGCCTTTTTTTTTTTCTCCCAAAAAAGAAACCCCTCCCGAACTCGGCGCCTTTCCCCCAA 210
Wa51 CTCCTTTCCATGCCTTTTTTTTTTTCTCCCAAAAAAGAAACCCCTCCCGAACTCGGCGCCTTTCCCCCAA 210
Wa42 CTCCTTTCCATGCCTTTTTTTTTTTCTCCCAAAAAAGAAACCCCTCCCGAACTCGGCGCCTTTCCCCCAA 210
     **********************************************************************

          Wa2-C      Wa2-3
Wa32 ACCATCCCTTTGGGGCGGGGGGACGCGCGAGCGGCTGCCGGTGGTGTCTCGCACTCTGGCTGCCCGCGAG 280
Wa51 ACCATCCCTTTGGGGCGGGGGGACGCGCGAGCGGCTGCCGGTGGGGTCTCGCACTCTGGCTGCCCGCGAG 280
Wa42 ACCATCCCTTTGGGGCGGGGGGACGCGCGAGCGGCTGCCGGTGGTGTCTCGCACTCTGGCTGCCCGCGAG 280
     ******************************************** *************************

                      Wa2-D
Wa32 CGCGCGTTCCCCGGAAACCCCGAGGGCGGGGAAGGGACAGAGGAAGGGCGCGCCACGGGCGAAAGAGGGG 350
Wa51 CGCGCGTTCCCCGGAAACCCCGAGGGCGGGGAAGGGACAGAGGAAGGGCGCGCCACGGGCGAAAGAGGGG 350
Wa42 CGCGCGTTCCCCGGAAACCCCGAGGGCGGGGAAGGGACAGAGGAAGGGCGCGCCACGGGCGAAAGAGGGG 350
     **********************************************************************

                          Wa2-4                              Wa2-5
Wa32 GGCGGGAAAAAAGAGAAAAAAAAGAGAGAGAGAGAGACGGGGGGGGGGGCTAAAAAAAGAGGGAAACCCAC 420
Wa51 GGCGGGAAAAAAGAGAAAAAAAAGAGAGAGAGAGAGACGGGGAGGGGGGCTAAAAAAAGAGGGAAAACCAC 420
Wa42 GGCGGGAAAAAAGAGAAAAAAAAGAGAGAGAGAGA--CGGGGAGGGGGGCTAAAAAAAGAGGGAGAACCAC 420
     **********************************  ***** **********  ********* * ****
                                                28s rRNA
                                   Wa2-E
Wa32 TCCCTCGTTCCGTTGCCCTCTTATCTATTCCCTTCTTTTCAAACGCGACCTCAGATCAGAAGAGACCACC 490
Wa51 CCCCTCGTTCCGGTGCCCTCTTATCTATTTCCTTCCTTTCAAACGCCCCCTCAGATCAAAAGAGACCACC 490
Wa42 TCCCTCGTTCCGTTGCCCTCTTATCTATTTCCTTCCTTTCGAACGCGACCTCAGATCAGACGAGACGACC 490
      *********** **************** ***** **** *****  ********** * ***** ***

Wa32 CGCTGAAATA 500
Wa51 CGCTGAAATA 500
Wa42 CGCTGAATTA 500
     ******* **
```

Microsatellite repeats: Wa2-1, Wa2-2, Wa2-3, Wa2-4, Wa2-5

Random repeats: Wa2-A, Wa2-B, Wa2-C, Wa2-D, Wa2-E

**Figure S3-(5).** Multiple sequence alignment of *W. attu* ITS2 (Wa2). The highlights are showing the position of microsatellite repeats Wa2-1 to Wa2-5 and random repeats Wa2-A to Wa2-E.

Cb: *Clarias batrachus*

```
ITS1                                    Cb1-1
Cb10 ACGGGTAGCCCCCGCAGCGGGGCGCACCTCCTCCTCCACCTCTTGCCCTTGAGATGGACCGAGGCGGCCT 70
Cb16 ACGGGTAGCCCCCGCAGCGGGGCGCACCTCCTCCTCCACCTCTTGCCCTTGAGATGGACCGAGGCGGCCT 70
Cb25 ACGGGTAGCACCCGCAGCGGGGCGCACCTCCTCCTCCACCTCTTGCCCTTGAGATGGACCGAGGCGGCCT 70
     ********* ************************************************************

Cb10 CCGTGACGCTGATGCCTTCCCGAAACCTTTCCGGTGACCGAGAAACAAACGATAGGGCGAGGGTACCTGC 140
Cb16 CCGTGACGCTGATGCCTTCCCGAAACCTTTCCGGTGACCGAGAAACAAACGATAGGGCGAGGGTACCTGC 140
Cb25 CCGTGACGCTGATGCCTTCCCGAAACCTTTCCGGTGACCGAGAAACAAACGATAGGGCGAGGGTACCTGC 140
     ********************************************************************** 140

Cb10 TGCTCGAGTGCGGCGCAACGGGGTTTGGGCTGGCCGTGGTCGGGCGCGTGCGCCGGTCGGGGAGGACATG 210
Cb16 TGCTCGAGTGCGGCGCAACGGGGTTTGGGCTGGCCGTGGTCGGGCGCGTGCGCCGGTCGGGGAGGACATG 210
Cb25 TGCTCGAGTGCGGCGCAACGGGGTTTGGGCTGGCCGTGGTCGGGCGCGTGCGCCGGTCGGGGAGGACATG 210
     **********************************************************************

                                                        Cb1-A
Cb10 TAGGTGTGGGGGTTGGGCATGTGGGTGAAAAGGTGCTACGCAGAGAGCCCCGGGCTCTCTGTGCGGTCGT 280
Cb16 TAGGTGTGGGGGTTGGGCATGTGGGTGAAAAGGTGCTACGCAGAGAGCCCCGGGCTCTCTGTGCGGTCGT 280
Cb25 TAGGTGTGGGGGTTGGGCATGTGGGTGAAAAGGTGCTACGCAGAGAGCCCCGGGCTCTCTGTGCGGTCGT 280
     **********************************************************************

Cb10 CCCGGGATCCCACTCCCTTCTCCCTCGCCTCTGTCTTCCATGCATGCGCGGGCGCTCCTCCCTTTCCCCC 350
Cb16 CCCGGGATCCCACTCCCTTCTCCCTCGCCTCTGTCTTCCATGCATGCGCGGGCGCTCCTCCCTTTCCCCC 350
Cb25 CCCGGGATCCCACTCCCTTCTCCCTCGCCTCTGTCTTCCATGCATGCGCGGGCGCTCCTCCCTTTCCCCC 350
     **********************************************************************

Cb10 CACGCACCCGCCGGCATGGTTCAATGACCCTTGAACGTTCACGGGGAGAGAGTTCTGGGGCGCCCTCCCC 420
Cb16 CACGCACCCGCCGGCATGGTTCAATGACCCTTGAACGTTCACGGGGAGAGAGTTCTGGGGCGCCCTCCCC 420
Cb25 CACGCACCCGCCGGCATGGTTCAATGACCCTTGAACGTTCACGGGGAGAGAGTTCTGGGGCGCCCTCCCC 420
     **********************************************************************
                                                 5.8s rRNA
Cb10 CGAGGTTTTTCCAACACACTGTTGATCTTTTGTGGACTCTTAGCAGTGGATCACTTGCTCGTGTGTCGAT 490
Cb16 CGAGGTTTTTCCAACACACTGTTGATCTTTTGTGGACTCTTAGCAGTGGATCACTTGCTCGTGTGTCGAT 490
Cb25 CGAGGTTTTTCCAACACACTGTTGATCTTTTGTGGACTCTTAGCAGTGGATCACTTGCTCGTGTGTCGAT 490
     **********************************************************************

Cb10 GAAGAACGC 499
Cb16 GAAGAACGC 499
Cb25 GAAGAACGC 499
     *********
```

Microsatellite repeats: Cb1-1

Random repeats: Cb1-A

**Figure S3-(6).** Multiple sequence alignment of *C. batrachus* ITS1. The highlights showing the position of microsatellite repeat Cb1-1 and random repeat Cb1-A.

Cg: *Clarias gariepinus*

```
                                                  18s rRNA   ITS1
                                                                              Cg1-1
Cg06 AAAANTCGTACAGGTTTTCGTAGATAAACCTGCGGAAGGATCATTAACGGGTAGCGT-----GGGGGGCG 70
Cg07 -----NNGTACAGGTTTTCGTAGATAAACCTGCGGAAGGATCATTAACGGGTAGCGT-----GGGGGGCG 70
Cg11 ---AAANGTACAGGTTTCCGTAGGTGAACCTGCGGAAGGATCATTAACGGGTAGCGCCCCGCGGGGGGCG 70
            ********** ***** * ******************************      ********

                 Cg1-2                                             Cg1-3
Cg06 CGCTGCCCTCTCCTCT-AACCCTCGG----------------------------------CCCCCAGCCC-- 140
Cg07 CGCTGCCTTCTCCTCT-AACCCTCGG----------------------------------CCCCCAGCCC-- 140
Cg11 CGCCTCCTCCTCCTCTTAACCCTCGGGACGGAAACCGAGGCTGGCCCGGGGGCTCACACCCCCGGCCCCC 140
     ***   **  ******* *********                                ***** ****

Cg06 -GCCGAGGCCTTCCCG----CTGACCGGGAAC---GAAAGTTGGGGCGAGGGTANCTGCTGCTCGAGCGT 210
Cg07 -GCCGAGGCCTTCCCG----CTGACCGGGAAC---GAAAGTTGGGGCGAGGGTATGTGCTGCTCGAGCGT 210
Cg11 CGCCGAGGCCTTCCCGAAACCTTTCCGGTGACCGTAAAAGATAGGGCGAGGGTACCTGCTGCTCGTGCGT 210
      ***************    **  ****  **     **** * ***********  ********* ****

Cg06 GGCGCGTTTTTTTGGTGGG---GCTGGC----TCGCTTCTCCC--TCGCCTCTTTCTTCCATGTGCGCAC 280
Cg07 GGCGCGTTTTTTTGGTGGG---GCTGGC----TCGCTTCTCCC--TCGCCTCTTTCTTCCATGTGCGCAC 280
Cg11 GGCGTGGTGTGCGTGCGTGCGTGCTGGTGGTGCCGGTGGTGACGGTAGGTGCTGGCCGTGGCGGGCGCGC 280
     **** * * * Cg1-4 * * *   *****       ** *  *                 Cg1-A

Cg06 GCGCCTC---CCTTCTGTTCCA-------CCCCCATGTTCACTAGGGAGAGTTCCGGNGCGCCCTCCCCT 350
Cg07 GCGCCTC---CCTTCTGTTCCA-------CCCCCATGTTCACTAGGGAGAGTTCCGGGGCGCCCTCCCCT 350
Cg11 GCCGGTCGGGCTGGGAGGGCCGAGCGGAGTCTGGATGGATGGTGCGGAGGGTGAAAAGGCGCTACGCGGA 350
     **   **   *     *  **        *   ***     *   **** **      ****    *

Cg06 AAGG------TTTGTACATACATTGTGTTGCATCTTTGTGGACGATGGCTTTCGAACCTAAAAAAC-AAC 420
Cg07 AAGG------TTTGTACATACATTGTGTTGCATCTTTGTGGACGATGGCTTTCGAACCTAAAAAAC-AAC 420
Cg11 GAAAGTGCTCTCTGTGCGGGCGTCATG--GGATCCCTCTCTCTCCCTCTCTCCGTTCTCACTCGCCTGGC 420
      *        * *** *   * *  **  * ***  * *  Cg1-5  * **  *  *     *   *

Cg06 TGACAACTCTTTGCGGTGGATCACTCAGCTCATGCATCAATGAAGAACGCAGCCAGCTGGGGCGGGCTGT 490
Cg07 TGACAACTCTTTGCGGTGGATCACTCAGCTCATGCATCAATGAAGAACGCAGCCAGCTGAAA-------- 482
Cg11 TGCCCCTTCCGTGCGCGCGCGCGCGCGTGCCTCTCCTTCCTCAGCCACCCATCCCCGCGCACGACCGCCC 490
     ** *   **  **** Cg1-6*  * * *   *Cg1-B* *   * *   ** ** **    *  Cg1-C

Cg06 AGACGGTCGATATCTTTATTGGGCGATGGTAAAATTGAAAC----------------------------- 531
Cg07 ----------------------------------------------------------------------
Cg11 CCCGCCTCGCCGGCGTGGTTCAATGACCCCGAGTTCTCTCCCGCCCACCCCGTTCGCGGGGGCGGGTGAG 560
                                             Cg1-D             Cg1-E
Cg06 ----------------------------------------------------------------------
Cg07 ----------------------------------------------------------------------
Cg11 TAGAGTTCCGGGGCGCCCTCCCCTGAGGTTTTTAACACACACTGTGTTGCATTCTCTGCTGGACGATGGC 630
               Cg1-F                  Cg1-G

Cg06 -------------------------------------
Cg07 -------------------------------------
Cg11 TTACCTAAAAAAAACCAAAAATGACAACTCTTAGCGG 667
           Cg1-7
                          5.8s rRNA
```

Microsatellite repeats: Cg1-1, Cg1-2, Cg1-3, Cg1-4, Cg1-5, Cg1-6, Cg1-7

Random repeats: Cg1-A, Cg1-B, Cg1-C, Cg1-D, Cg1-E, Cg1-F, Cg1-G

**Figure S3-(7).** Multiple sequence alignment of *C. gariepinus* ITS1. The highlights are showing the position of microsatellite repeat Cg1-1 to Cg1-7 and random repeats Cg1-A to Cg1-G.

Hf: *Heteropneustes fossilis*

```
                                                            18s rRNA  ITS1
                                                                                      Hf1-A
Hf40 --AANCGTACAGGTTTCCGTAGGTGAACCTGCGGAAGGATCATTAACGGGTAGCACCCCGCCGGGGGTGC 70
Hf44 AAANTCGTACAGGTTTCCGTAGGTGAACCTGCGGAAGGATCATTAACGGGTAGCACCCCGCCGGGGGTGC 70
Hf42 -AAANCGTACAGGTTTCCGTAGGTGAACCTGCGGAAGGATCATTAACGGGTAGCACCCCGCCGGGGGTGC 70
       *  *****************************************************************

                               Hf1-B                 Hf1-C
Hf40 GCCGCCTCCCATCATTACCCTCGAGAGAGCCCCGTGCCTTCTCGAAACCCTTCCGGTGACCGATGAAACA 140
Hf44 GCCGCCTCCCATCATTACCCTCGAGAGAGCCCCGTGCCTTCTCGAAACCCTTCCGGTGACCGATGAAACA 140
Hf42 GCCGCCTCCCATCATTACCCTCGAGAGAGCCCCGTGCCTTCTCGAAACCCTTCCGGTGACCGATGAAACA 140
     **********************************************************************

                                     Hf1-D
Hf40 ATGGGCGAGGGTACCTGCTGCTCGAGCGCCGCGGGTGGGGGCGCGTGCCGTGCGTAGAGGCCAAGAGTCT 210
Hf44 ATGGGCGAGGGTACCTGCTGCTCGAGCGCCGCGGGTGGGGGCGCGTGCCGTGCGTAGAGGCCAAGAGTCT 210
Hf42 ATGGGCGAGGGTACCTGCTGCTCGAGCGCCGCGGGTGGGGGCGCGTGCCGTGCGTAGAGGCCAAGAGTCT 210
     **********************************************************************

             Hf1-1
Hf40 GGGGGTGTGTGTGTGAGAGAAAGGCGCCACGCGGAAGGCCCAGTGCCTCCGTGCGGGCGGCTGGATCGAC 280
Hf44 GGGGGTGTGTGTGTGAGAGAAAGGCGCCACGCGGAAGGCCCAGTGCCTCCGTGCGGGCGGCTGGATCGAC 280
Hf42 GGG--TGTGTGTGTGAGAGAAAGGCGCCACGCGGAAGGCCCAGTGCCTCCGTGCGGGCGGCTGGATCGAC 280
     ***  *****************************************************************

              Hf1-2                   Hf1-3                   Hf1-4
Hf40 CACTCTCTCCTTCTCTATCGCCCCTCACACACGCGCGCGCGC--CCACTCAACCCCCC-GCGCTGTCGGC 350
Hf44 CACTCTCTCCTTCTCTATCGCCCCTCACACACGCGCGCGCGC--CCACTCAACCCCCC-GCGCTGCCGGC 350
Hf42 CACTCTCTCCTTCTCTATCGCCCCTCACACACGCGCGCGCGCGCCCACTCAACCCCCCCGCGCTGCCGGC 350
     ******************************************  ************** ****** ****

                                                              Hf1-E
Hf40 GTGGTTCAATGACCCCGAGTTCTATCTCCGGTGACGGGGAGAGTTCCGGGGCGCCCTCCCCGGATGGTTT 420
Hf44 GTGGTTCAATGACCCCGAGTTCTATCCCCGGTGACGGGGAGAGTTCCGGGGCGCCCTCCCCGGATGGTTT 420
Hf42 GTGGTTCAATGACCCCGAGTTCTATCCCCGGTGACGGGGAGAGTTCCGGGGCGCCCTCCCCGGATGGTTT 420
     ************************** *******************************************
                                                                  5.8s rRNA
                       Hf1-5                       Hf1-6
Hf40 ATACACAGTCGATATCTTTTTTGGGCGATGGCAAAATTGAAACAAAAAAACGAAACTGACAACTCTTAGC 490
Hf44 ATACACAGTCGATATCTTTTTTGGGCGATGGCAAAATTGAAACAAAAAAACGAAACTGACAACTCTTAGC 490
Hf42 ATACACAGTCGATATCTTTTTTGGGCGATGGCAAAATTGAAACAAAAAAACGAAACTGACAACTCTTAGC 490
     **********************************************************************

Hf40 GGTGGATCACTCGGCTCGTGCGTCGATGAAAAACC--AGCCCAGCTGAGA- 541
Hf44 GGTGGATCACTCGGCTCGTGCGTCGATGAAGAACGCAGGCCCAGCTGAA-- 541
Hf42 GGTGGATCACTCGGCTCGTGCGTCGATGAAGAACGC--AGCCAGCTGAAGC 541
     ****************************** ***      ********
```

Microsatellite repeats: Hf1-1, Hf1-2, Hf1-3, Hf1-4, Hf1-5, Hf1-6

Random repeats: Hf1-A, Hf1-B, Hf1-C, Hf1-D, Hf1-E

**Figure S3-(8).** Multiple sequence alignment of *H. fossilis* ITS1. The highlights are showing the position of microsatellite repeats Hf1-1 to Hf1-6 and random repeats Hf1-A to Hf1-E.

```
Ss: Sperata seenghala
                                                          18s rRNA  ITS1
Ss02 AAANTCGTACAGGTTTCCGTAGGTGAACCTGCGGAAGGATCATTAACGGGTTAGCGTCCGCGGGGACGCG 70
Ss11 AAANTCGTACAGGTTTCCGTAGGTGAACCTGCGGAAGGATCATTAACGGGTTAGCGTCCGCGGGGACGCG 70
Ss09 AAAATCGTACAGGTTTCCGTAGGTGAACCTGCGGAAGGATCATTAACGGGTTAGCGTCCGCGGGGACGCG 70
Ss01 AAAATCGTACAGGTTTCCGTAGGTGAACCTGCGGAAGGATCATTAACGGGGTAGCGTCCGCGGGGACGCG 70
     *** ********************************************** *******************

                                         Ss1-A
Ss02 CCTGCTCCTCGACACACGGCGCCGAGGCTCTCGCTCTTT-CCCCTCGACAAAAGGAGAAACGTGAGCCGA 140
Ss11 CCTGCTCCTCGACACACGGCGCCGAGGCTCTCGCTCTTT-CCCCTCGACAAAAGGAGAAACGTGAGCCGA 140
Ss09 CCTGCTCCTCGACACACGGCGCCGAGGCTCTCGCTCTTT-CCCCTCGACAAAAGGAGAAACGTGAGCCGA 140
Ss01 CCTGCTCCTCGACACACGGCGCCGAGGCTCGCGCTTTTTTCCCCTTGACAAAAGGAGAAACGTGAGCCGA 140
     ****************************** **** *** ***** ***********************

         Ss1-B                         Ss1-C
Ss02 GGTCCCTTTTACGGTATCCCAAGAAGACGGGGGAAAAAGAGAAAGGGGACGAGAAAAGGGAATCGGAAGG 210
Ss11 GGTCCCTTTTACGGTATCCCAAGAAGACGGGGGAAAAAGAGAAAGGGGACGAGAAAAGGGAATCGGAAGG 210
Ss09 GGTCCCTTTTACGGTATCCCAAGAAGACGGGGGAAAAAGAGAAAGGGGACGAGAAAAGGGAATCGGAAGG 210
Ss01 GGTCCCTTTTACGGTATCCCAAGAAGACGGGGGAAAAAGAGAAAGGGGACGAGAAAAGGGAATCGGAAGG 210
     **********************************************************************

              Ss1-1
Ss02 GGGAAAAGAAAGAAAGAAAGAAAGACGGACGGGGCGGAGGGTACCTACCGCTCGCCGGCGCTACGCGGAG 280
Ss11 GGGAAAAGAAAGAAAGAAAGAAAGACGGACGGGGCGGAGGGTACCTACCGCTCGCCGGCGCTACGCGGAG 280
Ss09 GGGAAAAGAAAGAAAGA--------CGGACGGGGCGGAGGGTACCTACCGCTCGCCGGCGCTACGCGGAG 280
Ss01 GGGAAAAGAAAGAAAGAA----AGACGGACGGGGCGGAGGGTACCTACCGCTCGCCGGCGCTACGCGGAG 280
     *****************        *********************************************

               Ss1-2                Ss1-3              Ss1-4
Ss02 GTCCTTTGGGGGTAGGGGGG-TGA------GACAGACGGGGCTCCTCGTCCCTTCCTTCCTTCC----CT 350
Ss11 GTCCTTTGGGGGTAGGGGGG-TGA------GACGGACGGGGCTCCTCGTCCCTTCCTTCCTTCC----CT 350
Ss09 GTCCTTTGGGGGTAGGGGGGGTGA------GACGGACGGGGCTCCTCGTCCCTTCCTTCCTTCCTTCCCT 350
Ss01 GTCCTTTGGGGGTAGGGGGTGAGACGGACGGACGGACGGGGCTCCTCGTCCCTTCCTTCCCTCG----CT 350
     *******************   **       *** ************************** **    **

                                                          Ss1-5       Ss1-6
Ss02 CCCTCCCTCTCTCCCTCTGTGCGGGCGCCTGGGCCGAAAGAGGTTCGAAGACCCCTCCCCTCGCGGGGTG 420
Ss11 CCCTCCCTCTCTCCCTCTGTGCGGGCGCCTGGGCCGAAAGAGGTTCGAAGACCCCTCCCCTCGCGGGGTG 420
Ss09 CCCTCCCTCTCTCCCTCTGTGCGGGCGCCTGGGCCGAAAGAGGTTCGAAGACCCCTCCCCTCGCGGGGTG 420
Ss01 CCCTCGCTCCCTCCCTCTGTGCGGGCGCCTGGGCCGAAAAAGGTTCGAAGACCCCTCCCCTCGCGGGGTG 420
     ***** *** ****************************** *****************************

                     Ss1-D              Ss1-7
Ss02 GGGCGCCCTCCACTCGTTCTTTTCTTTATTTCTCTCCCCCACCCCGACCCCCGGATCCTGGTTCTACCCT 490
Ss11 GGGCGCCCTCCACTCGTTCTTTTCTTTATTTCTCTCCCCCACCCCGACCCCCGGATCCTGGTTCTACCCT 490
Ss09 GGGCGCCCTCCACTCGTTCTTTTCTTTATTTCTCTCCCCCACCCCAACCCCCGGATCCTGGTTCT-CCCT 490
Ss01 GGGCGCCCTCCACTCGGTCTTTTCTTTATTTTTCTCCCCCACCCCACACCCCGGATCCTGGTTCT-CCCT 490
     **************** ************** ************    ***************** ****

          Ss1-8                       Ss1-9
Ss02 TTCTCTCTCGTCTCCCCC-GATCCTCCAAGACCCGTATTTTTTTCATTCCCATCGTGTGGCTCGGACGAA 560
Ss11 TTCTCTCTCGTCTCCCCC-GATCCTCCAAGACCCGTATTTTTTTCATTCCCATCGTGTGGCTCGGACGAA 560
Ss09 TTCTCTCTCGTCTCCCCC-GATCCTCCAAGACCCGTATTTTTTTCATTCCCATCGTGTGGCTCGGACGAA 560
Ss01 TTCTCTCTCGTCTCCCCCCGATCCTCCAAGACCCGTATTCTTT--ATTCCCATCGGGTGGCTCGGACAAA 560
     ****************** ******************** ***   ********** *********** **

                                Ss1-10              5.8s rRNA
Ss02 CCGTCTCGGTGAAATAAACGAGAGAAAAAAA-CGAACCATACATAG------------------------ 630
Ss11 ACGTCTCGGTGAAATAAACGAGAGAAAAAAA-CGAAGCATAAAGAGCTGACAACTCTTAGCGGTGGATCA 630
Ss09 ACGTCTCGGTGAAATAGACGAGAGAAAAAAAACGAAGCATAAAGAGCTGACAACTCTTAGCGGTGGATCA 630
Ss01 ACGTCTCGGTGAAAGAAACGAGAGAAAAAAA-CCAAGCATAAAGAGCTGACAACTCTTAGCGGTGGATCA 630
      ************* * ********** *** * ** **** * **

Ss02 -------------------------------------- 668
Ss11 CTCGGCTCGTGCGTCGATGAAGAACGCAGGCCAGCTGA 668
Ss09 CTCGGCTCGTGCGTCGATGAAGAACGCAGCCCAGCGGA 668
Ss01 CTCGGCTCGTGCGTCGATGAAGAAGCAGCCCCGGCGGA 668
```

Microsatellite repeats: Ss1-1, Ss1-2, Ss1-3, Ss1-4, Ss1-5, Ss1-6, Ss1-7, Ss1-8, Ss1-9, Ss1-10

Random repeats: Ss1-A, Ss1-B, Ss1-C, Ss1-D

**Figure S3-(9).** Multiple sequence alignment of *S. seenghala* ITS1. The highlights are showing the position of microsatellite repeats Ss1-1 to Ss1-10 and random repeats Ss1-A to Ss1-D.

Wa: *Wallago attu*

```
                                                     18s rRNA   ITS1
Wa42 -AAAATCGTACAGGTTTCCGTAGGTGAACCTGCGGAAGGATCATTAACGGGGCAGCCGCCGTCGGGGCGA 70
Wa32 --AAANCGTACAGGTTTCCGTAGGTGAACCTGCGGAAGGATCATTAACGGGGCAGCCGCCGTCGGGGCGA 70
Wa51 --AANTCGTACAGGTTTCCGTAGGTGAACCTGCGGAAGGATCATTAACGGGGCAGCCGCCGTCGGGGCGA 70
Wa71 AAAAANCGTACAGGTTTCCGTAGGTGAACCTGCGGAAGGATCATTAACGGGGCAGCCGCCGTCGGGGCGA 70
        **   *************************************************************

                                                  Wa1-A
Wa42 GCGCCTTCTCACCCTCCTCCTCACGCGACGTGCCGAAGGCCCGGGGCAGTCGCTCCCGGCCCGAGGCGC 140
Wa32 GCGCCTTCTCACCCTCCTCCTCACGCGACGTGCCGAAGGCCCGGGGCAGTCGCTCCCGGCCCGAGGCGC 140
Wa51 GCGCCTTCCCACCCTCCTCCTCACGCGACGTGCCGAAGGCCCGGGGCAGTCGCTCCCGGCCCGAGGCGC 140
Wa71 GCGCCTTCTCACCCTCCTCCTCACGCGACGTGCCGAAGGCCCGGGGCAGTCGCTCCCGGCCCGAGGCGC 140
     ******** ************************************************************

                                                                  Wa1-B
Wa42 ATTGCTTTCGGTCTCCTTCGATCAAAAAACAAAGGGAAGGGGAAAGCAAAGCAAGGTCAAAAACCTTTGG 210
Wa32 ATTGCTTTCGGTCTCCTTCGATCAAAAAACAAAGGGAAGGGGAAAGCAAAGCAAGGTCAAAAACCTTTGG 210
Wa51 ATTGCTTTCGGTCTCCTTCGATCAAAAAACAAAGGGAAGGGGAAAGCAAAGCAAGGTCAAAAACCTTTGG 210
Wa71 ATTGCTTTCGGTCTCCTTCGATCAAAAAACAAAGGGAAGGGGAAAGCAAAGCAAGGTCAAAAACCTTTGG 210
     **********************************************************************

Wa42 GAAAGAGAGAAAAAAG                   Wa1-1                         Wa1-2 226
Wa32 GAAAGAGAGAAAAAAAGATAAAAAAAAGAAAACGAGGAAAAAAAAAAAAGAAAAAAGAAAGAAAAACACCCCC 280
Wa51 GAAAGAGAGAAAAAAAGATAAAAAAA-GAAAACGAGGGAAAAAAAAAAAAAAAAAAANNAAAAAAAAACCCCCC 280
Wa71 GAAAGAGAGAAAAAA                                                        225
     ************************ ********** ********** ******    * *****   *****

                                 Wa1-C                         Wa1-3
Wa42
Wa32 CCTCCTCCGGCAAGGGTACCTACCGGCCGGACAGAACGGGCTCAGGAATGAAAACAAAAAGTGAATCGAT 350
Wa51 CCCCCC
Wa71
     **  **

                Wa1-D                                         Wa1-E
Wa42
Wa32 TCTCTTTTTATTTCCTCCGTCCCACTGTCGGGCCCAGGTTCAATGACCCCGGCCGACCCGGGGCGCCCTT 420
Wa51
Wa71

                                                Wa1-4
Wa42
Wa32 GCGATGGAGCCGTTGTGATATTTCTTTTTTTTCTTTTTTGACTTTTTTTTTTCTTTCTTATTCTTCTTCTTC 490
Wa51
Wa71

                                                                  Wa1-F
Wa42
Wa32 TCTCCCCTTCTTTTTTTGTTTTGCTGCCGGTTTCCCCGCCATTCCCTTTGTTATATTTTGTTGTTTCTCTT 560
Wa51
Wa71

                                             Wa1-5
Wa42
Wa32 TACCCTTCTCTGGCTTGGGCAAAGGTAAAACAAAACCAAAAAAAAAAAA 609
Wa51
Wa71
```

Microsatellite repeats: Wa1-1, Wa1-2, Wa1-3, Wa1-4

Random repeats: Wa1-A, Wa1-B, Wa1-C, Wa1-D, Wa1-E, Wa1-F

**Figure S4-(10).** Multiple sequence alignment of *W. attu* ITS1. The highlights are showing the position of microsatellite repeats Wa1-1 to Wa1-5 and the random repeats Wa1-A to Wa1-F.
