## Supplementary material for "Exploring unique sequence repeat patterns and secondary structures in rDNA internal transcribed spacers ITS1 and ITS2 for characterization of catfish species": Supplementary Data S4.pdf

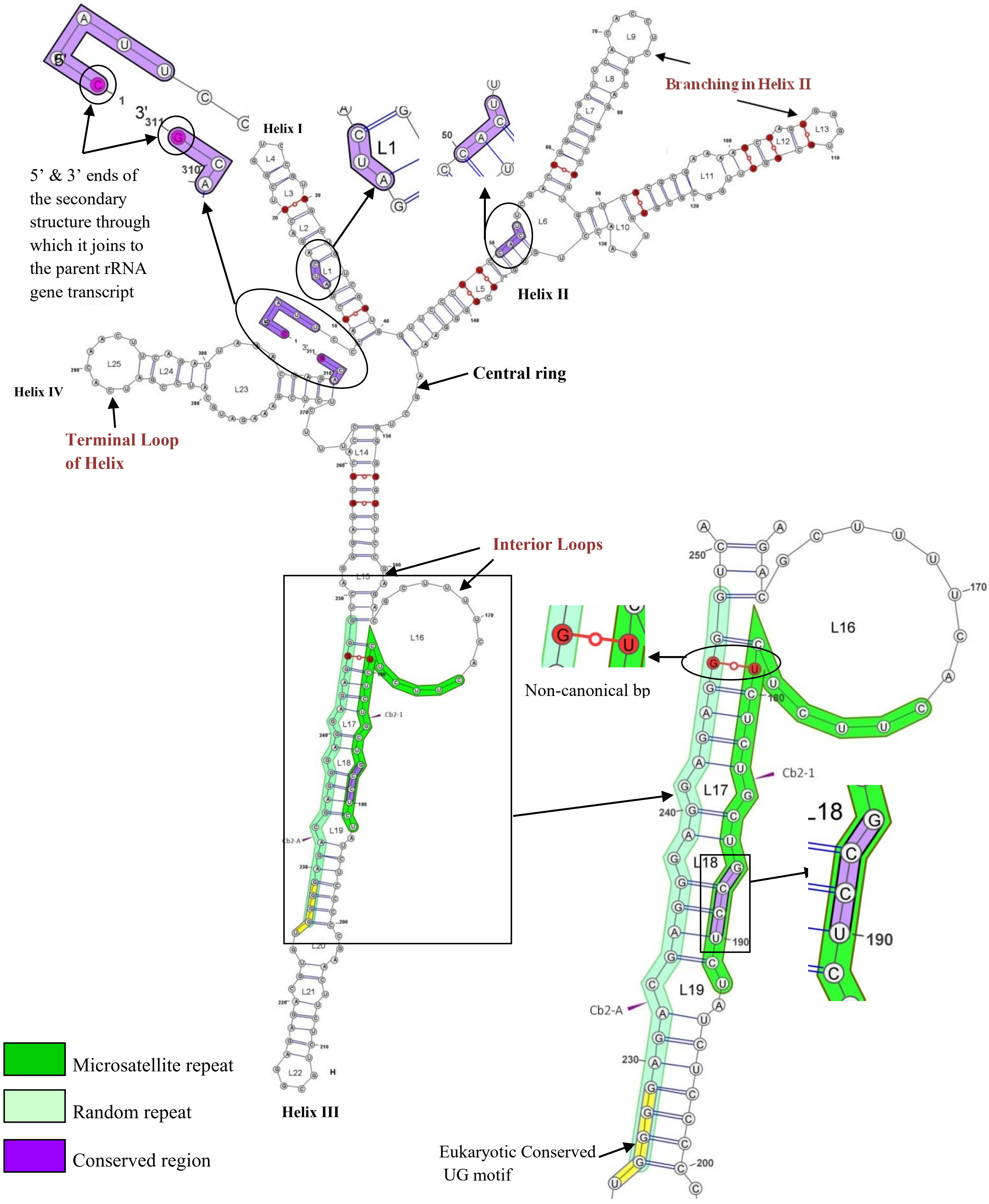


**Figure S4-(1)**. ITS2 secondary structure of *Clarias batrachus*. Microsatellite repeats, Random repeats, eukaryotic conserved motifs, catfish conserved motifs (highlighted in blue), non-canonical bp are shown in magnified view round the secondary structure.

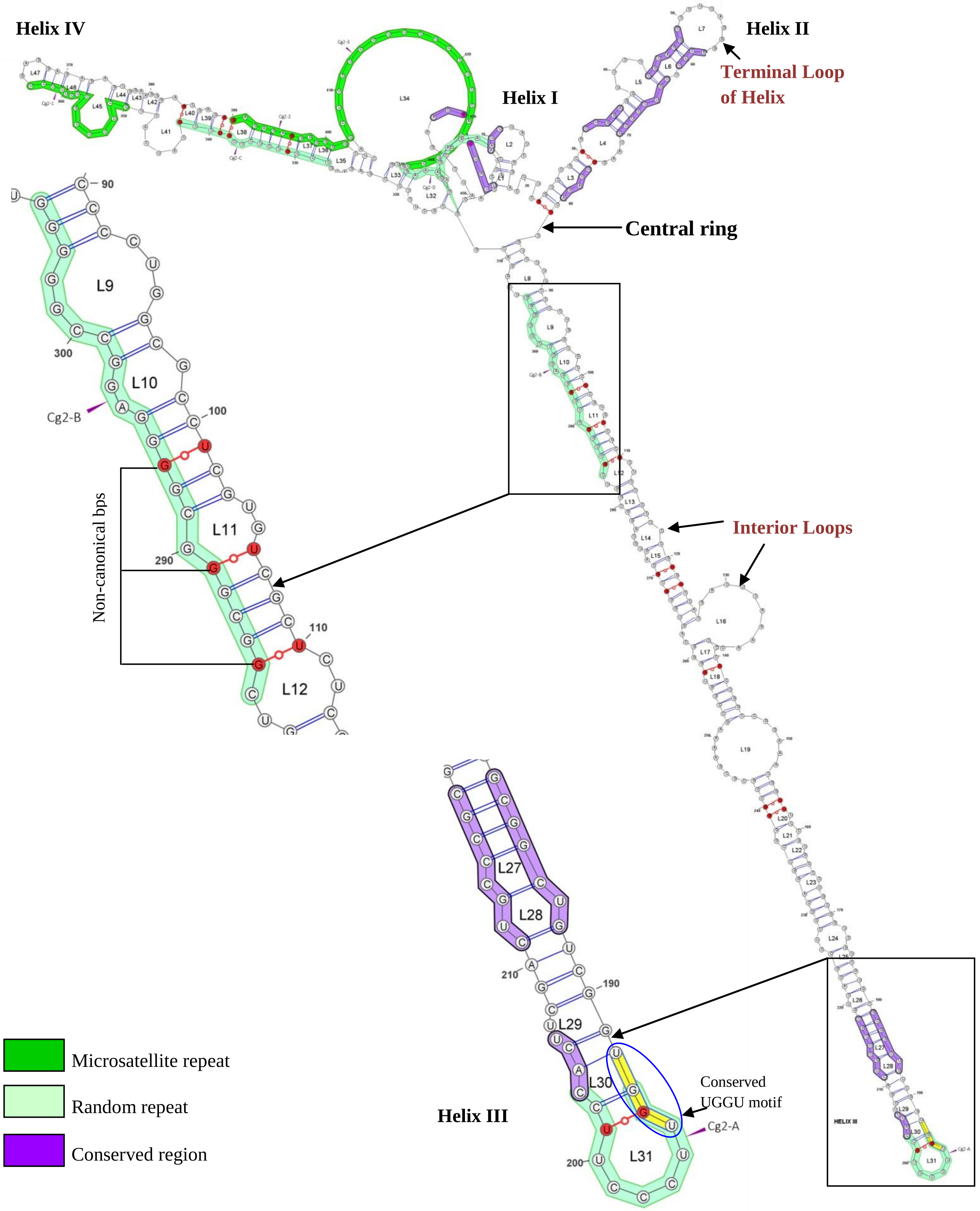


**Figure S4-(2A).** ITS2 secondary structure of *Clarias gariepinus*. The 4-helix secondary structure is showing the central ring with its 4 helices from Helix I to Helix IV arranged in clockwise direction. The helices are interrupted with loops (L) of variable size labeled as L1 to L34. Microsatellite repeats, Random repeats, eukaryotic conserved motifs, catfish conserved motifs (highlighted in blue), non-canonical bp are shown in magnified view round the secondary structure.

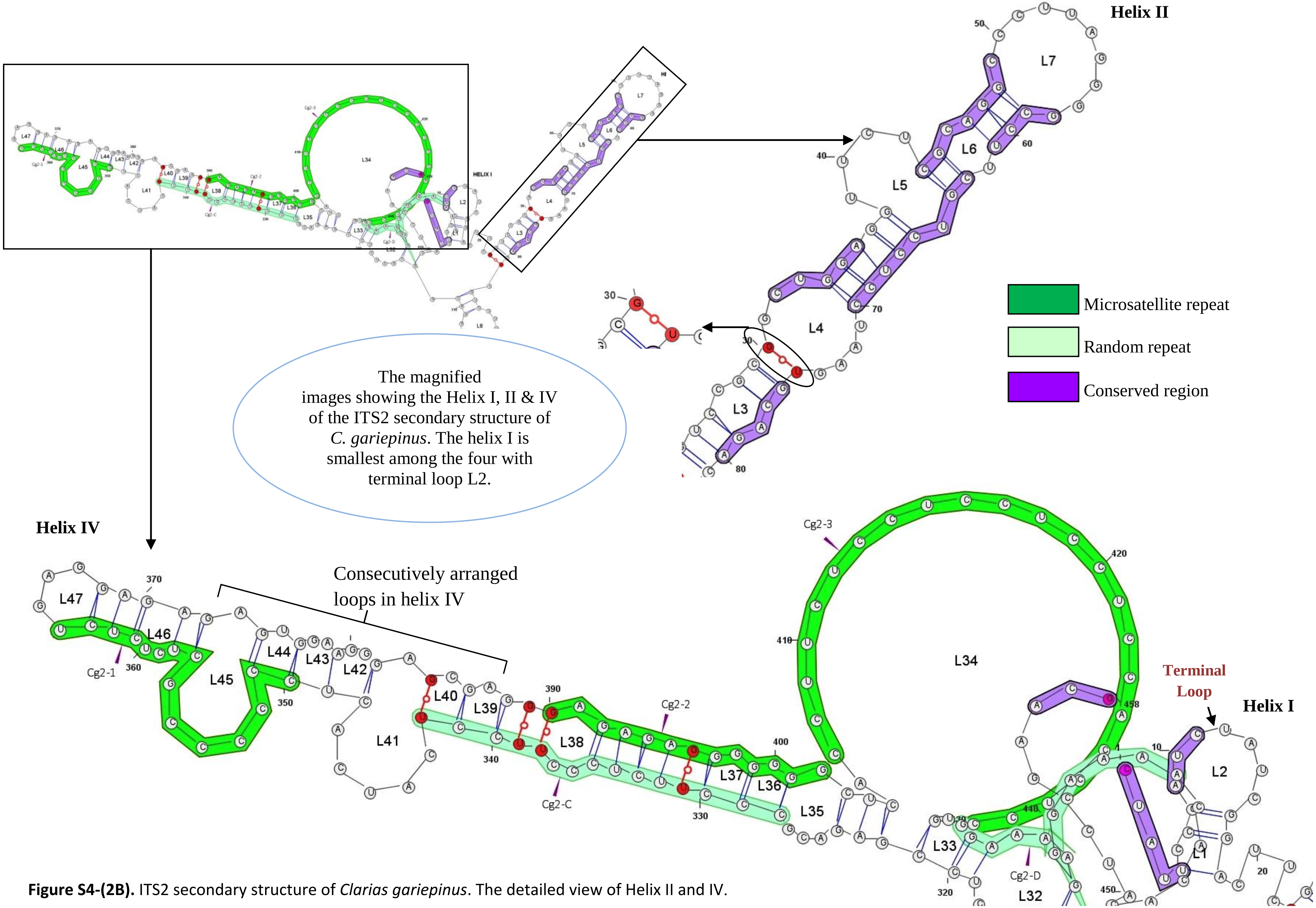


**Figure S4-(2B).** ITS2 secondary structure of *Clarias gariepinus*. The detailed view of Helix II and IV.

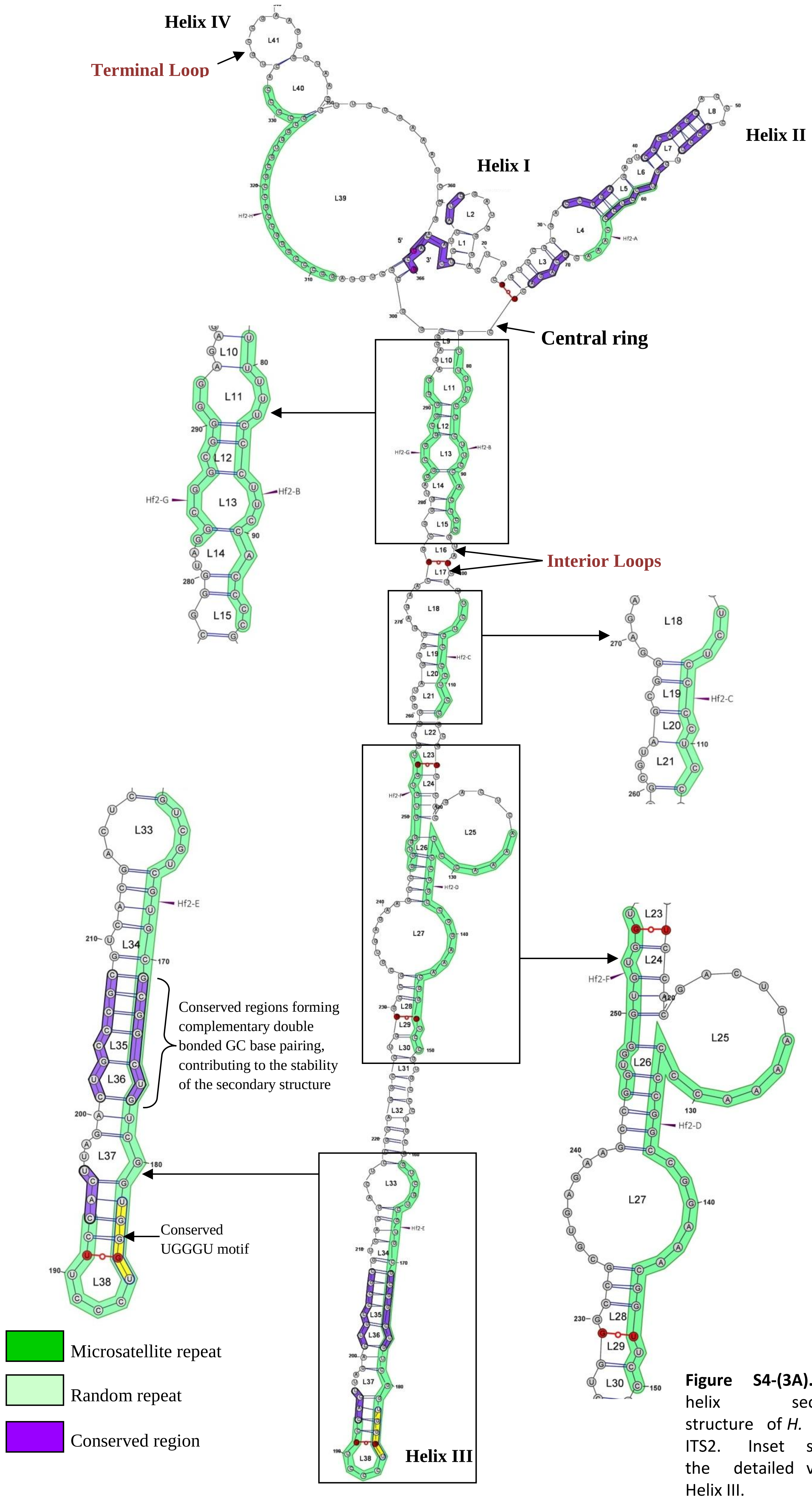


**Figure S4-(3A).** 4-helix secondary structure of *H. fossilis* ITS2. Inset showing the detailed view of Helix III.

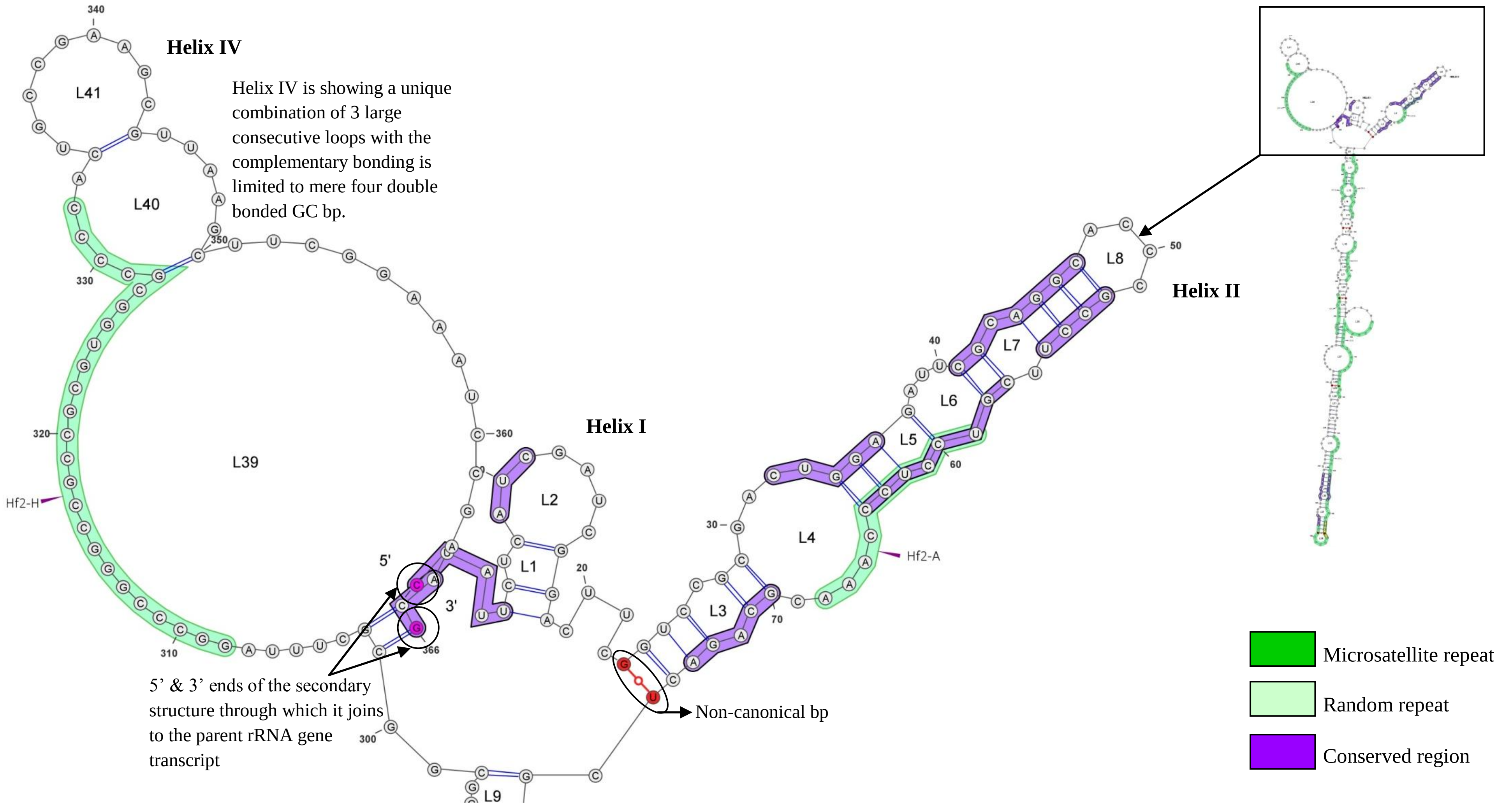


**Figure S4-(3B).** ITS2 secondary structure of *H. fossilis.* Inset showing the detailed view of central ring and the helix I, II and IV.

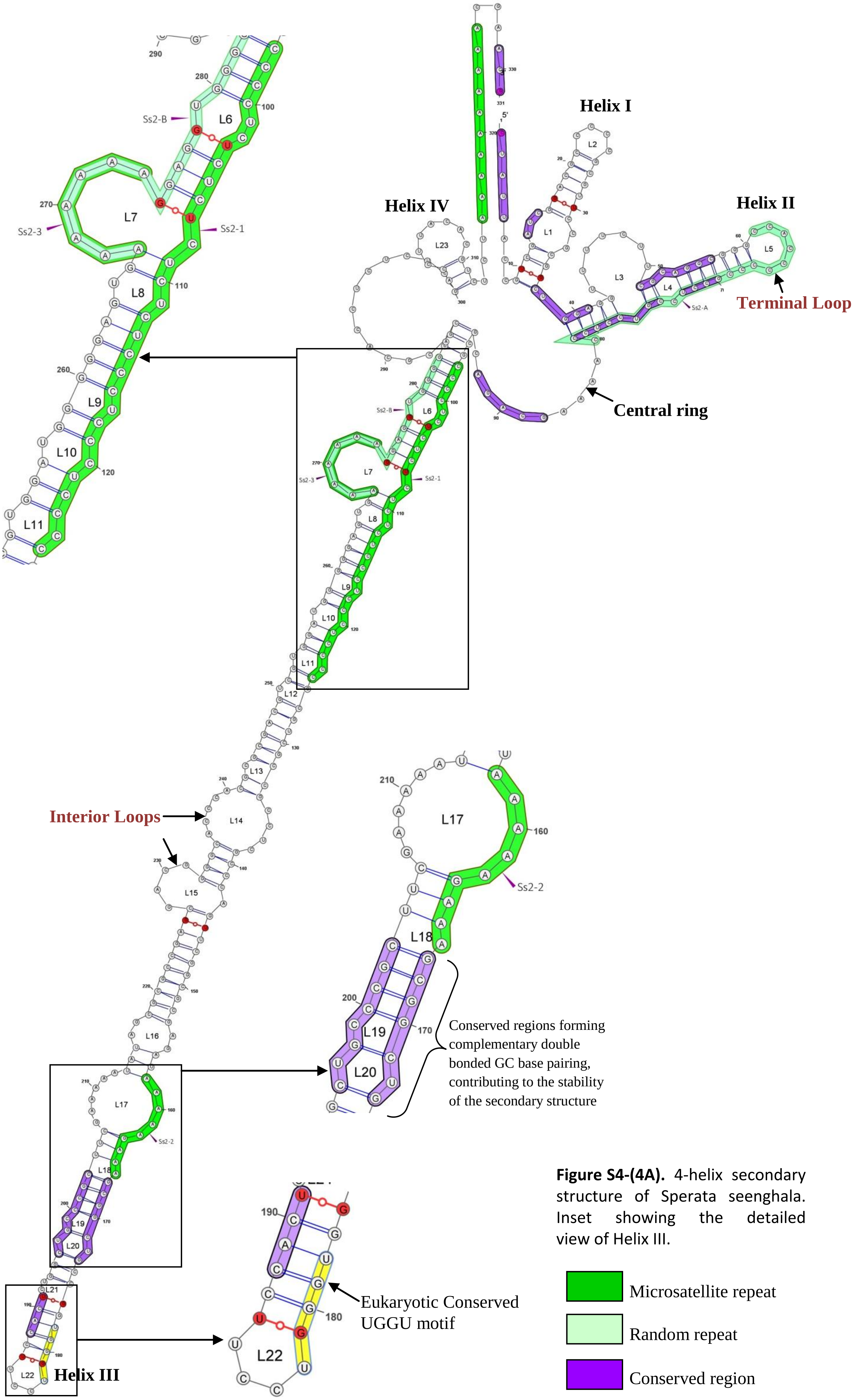


**Figure S4-(4A).** 4-helix secondary structure of Sperata seenghala. Inset showing the detailed view of Helix III.

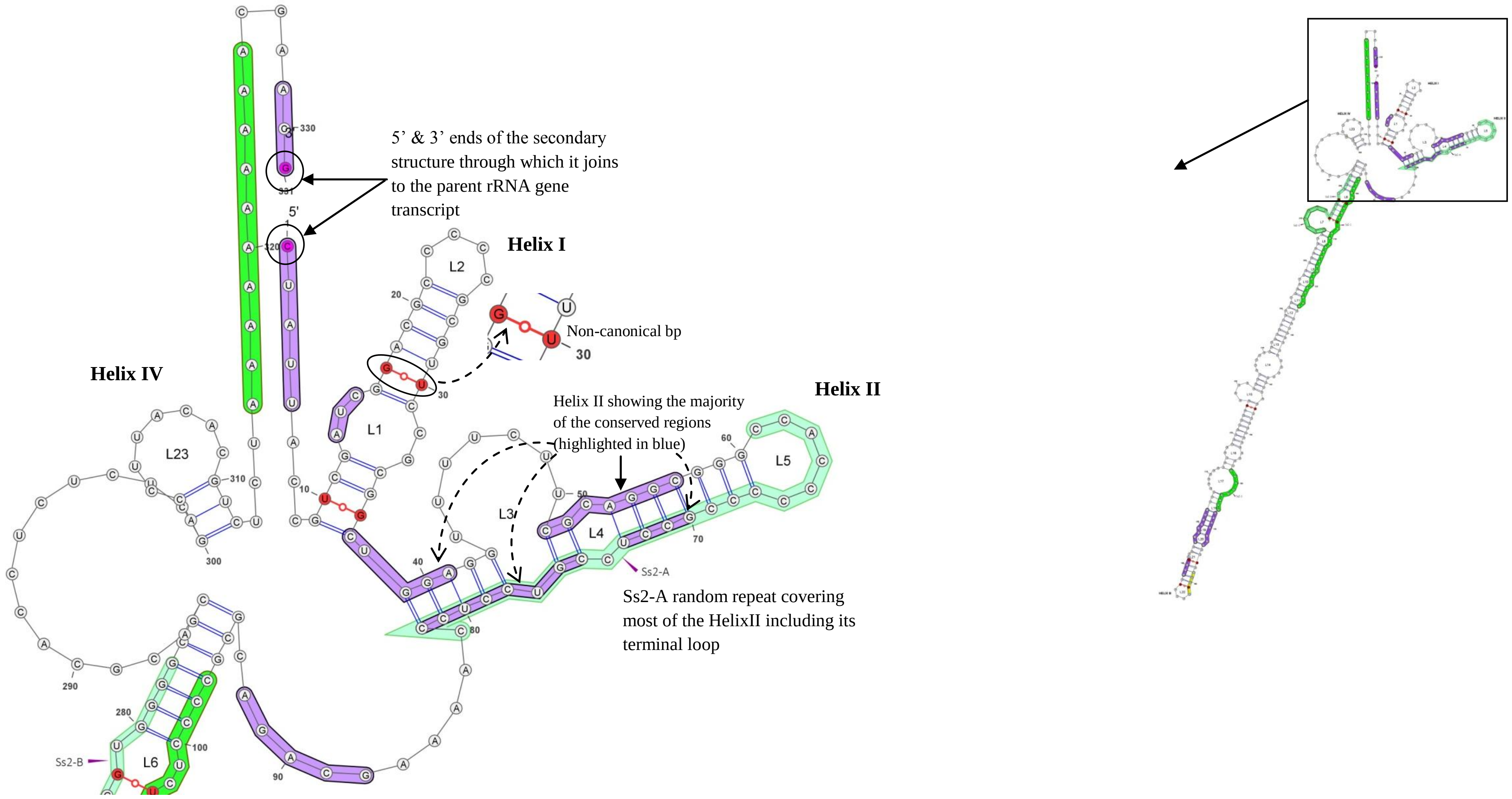


**Figure S4-(4B).** 4-helix secondary structure of *Sperata seenghala* ITS2. Inset showing the detailed view of central ring and the helix I, II and IV.

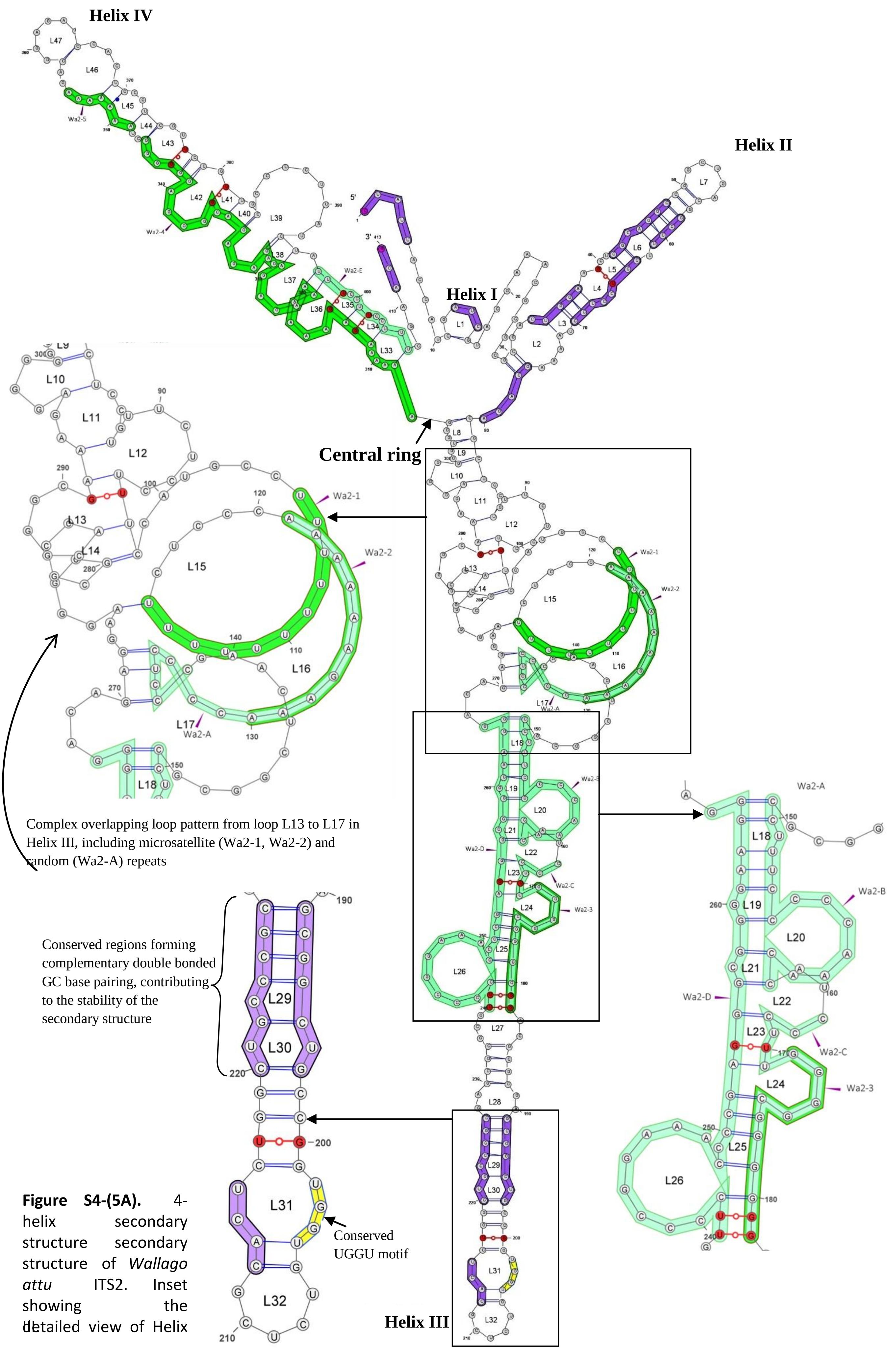


**Figure S4-(5A).** 4-helix secondary structure secondary structure of *Wallago attu* ITS2. Inset showing the detailed view of Helix III

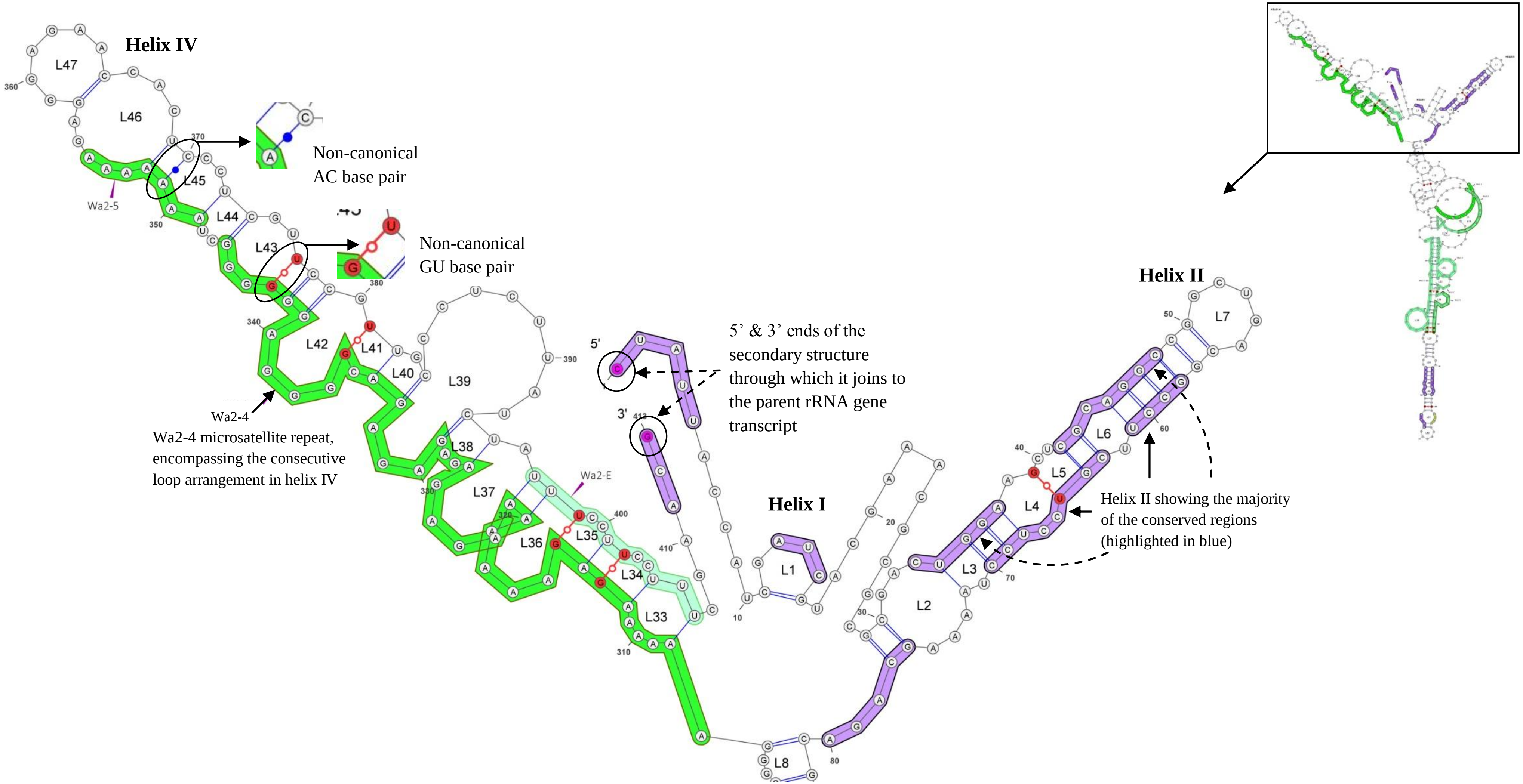


**Figure S4-(5B).** 4-helix secondary structure of *Wallago attu* ITS2. Inset showing the detailed view of central ring and the helix I, II and IV.

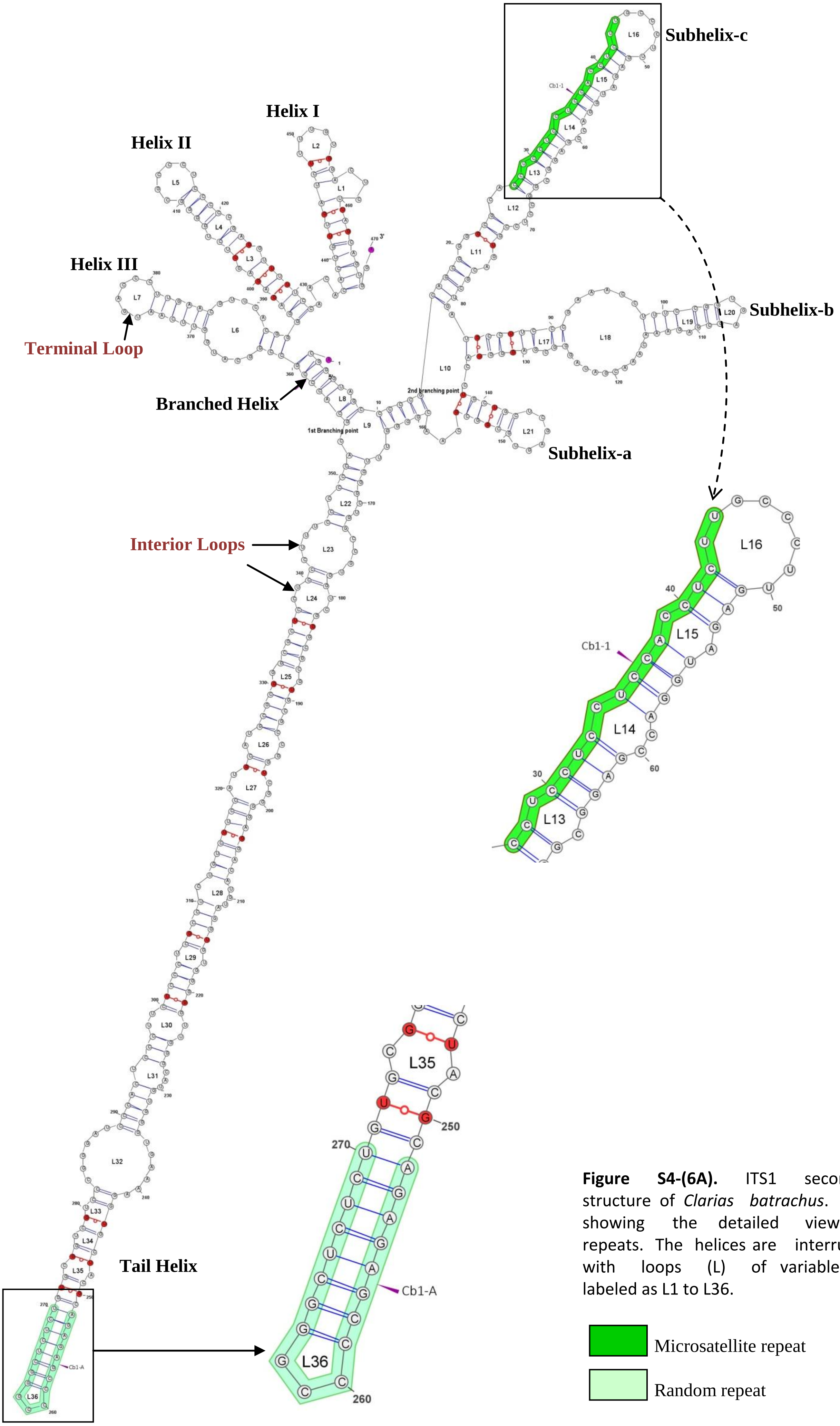


**Figure S4-(6A).** ITS1 secondary structure of *Clarias batrachus*. Inset showing the detailed view of repeats. The helices are interrupted with loops (L) of variable size labeled as L1 to L36.

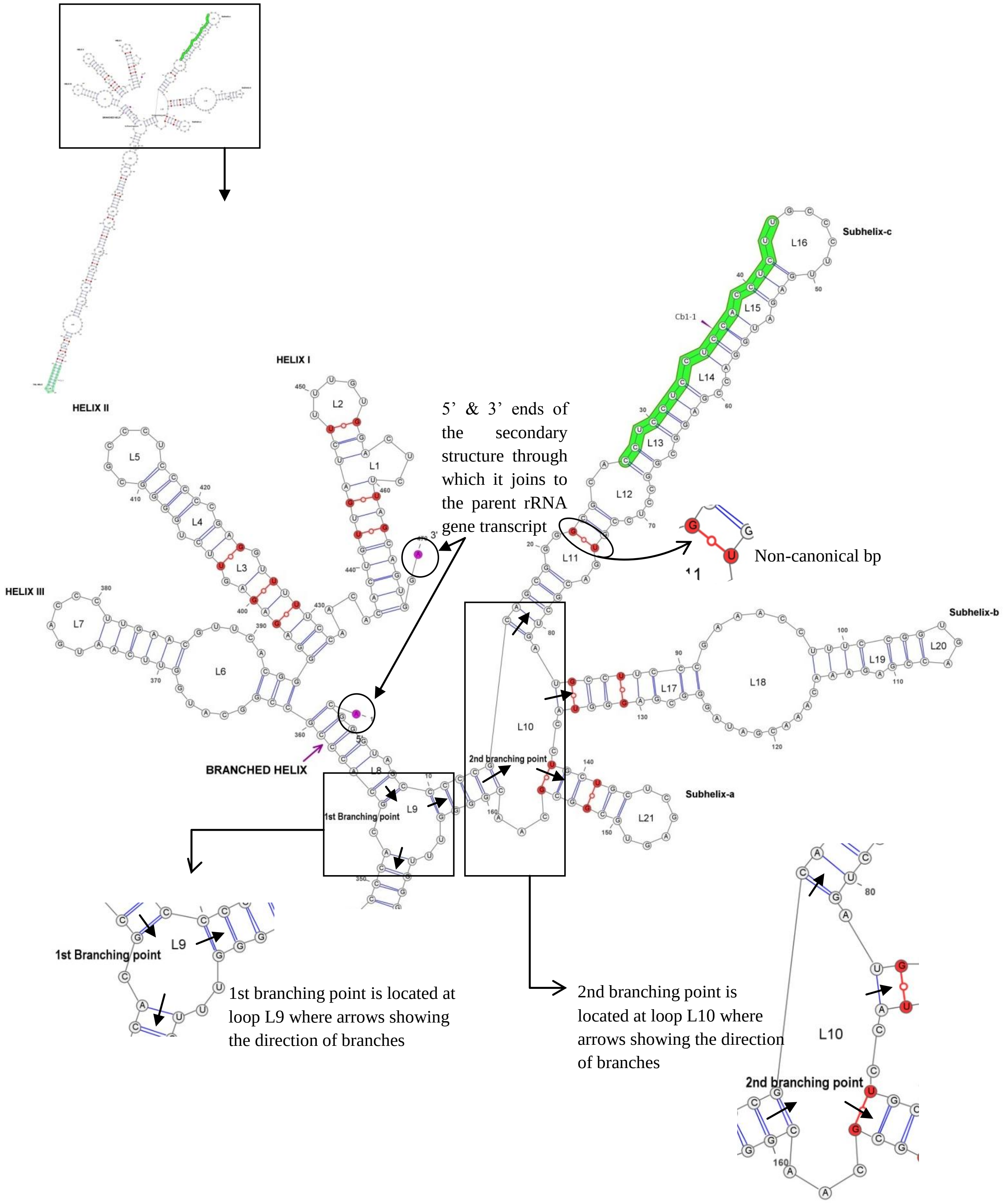


**Figure S4-(6B).** ITS1 secondary structure of *Clarias batrachus*. Inset showing the detailed of branched helix.

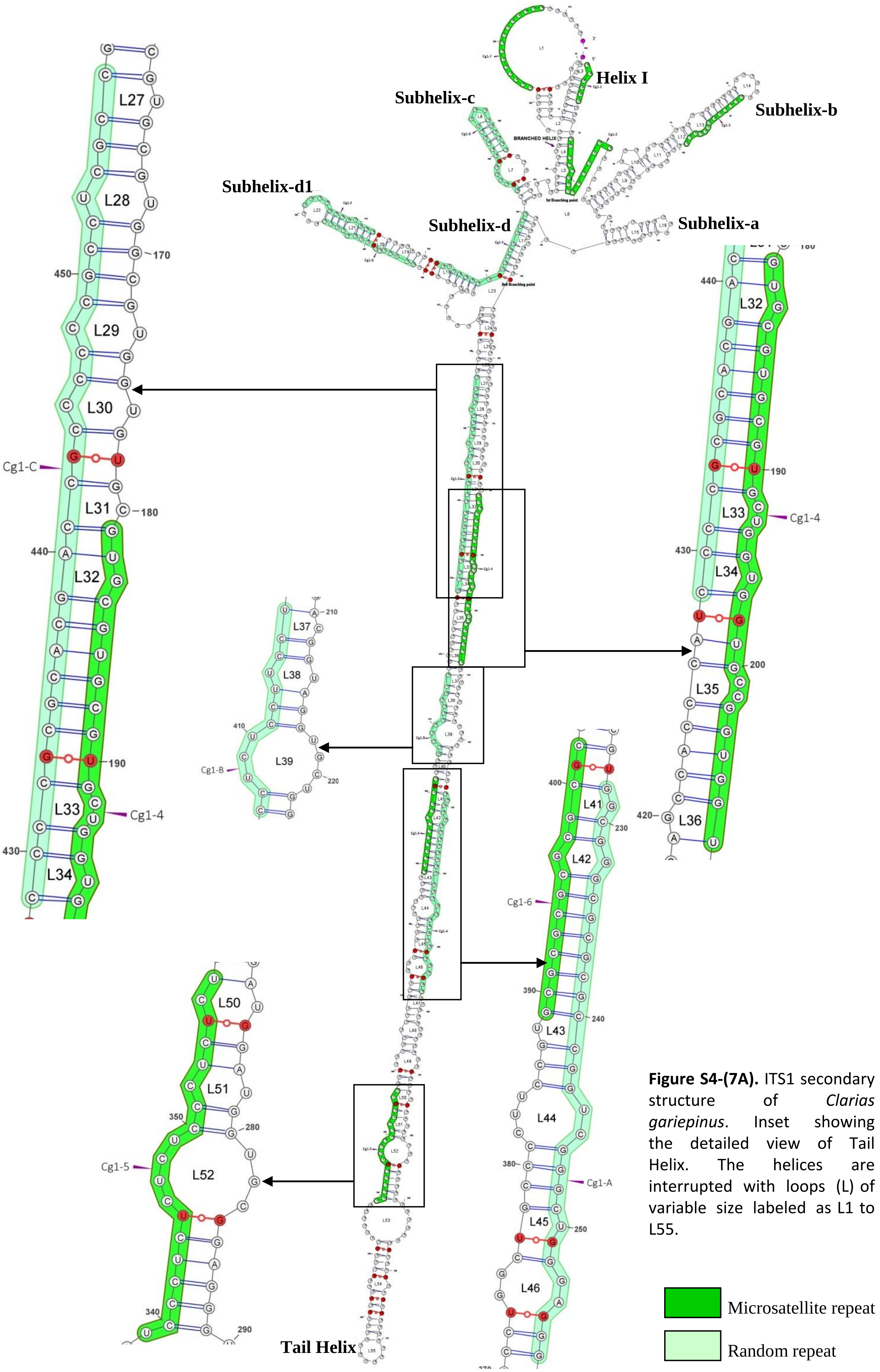


**Figure S4-(7A).** ITS1 secondary structure of *Clarias gariepinus*. Inset showing the detailed view of Tail Helix. The helices are interrupted with loops (L) of variable size labeled as L1 to L55.

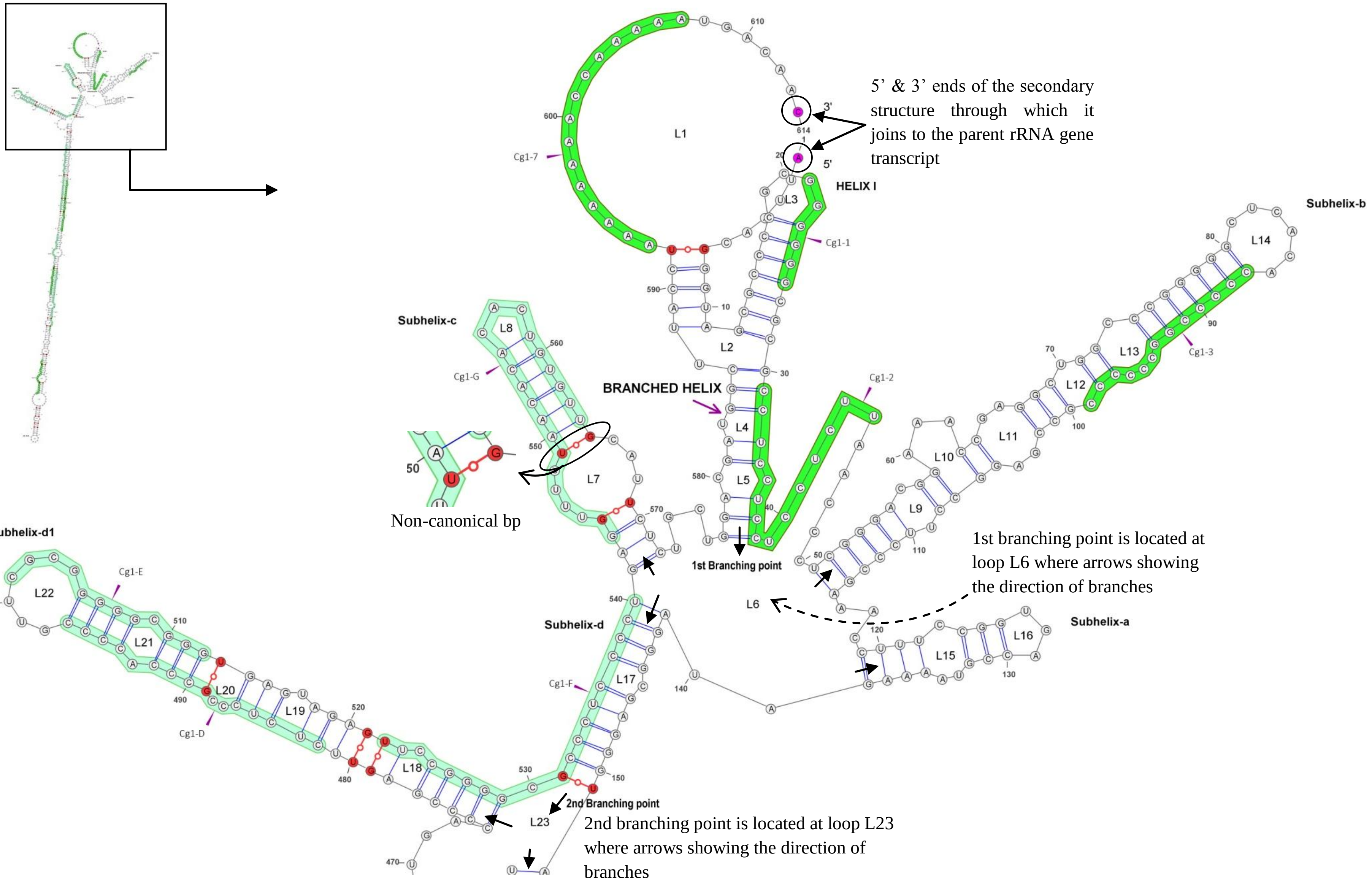


**Figure S4-(7B).** ITS1 secondary structure of *Clarias gariepinus*. Inset showing the detailed view of upper branched region.

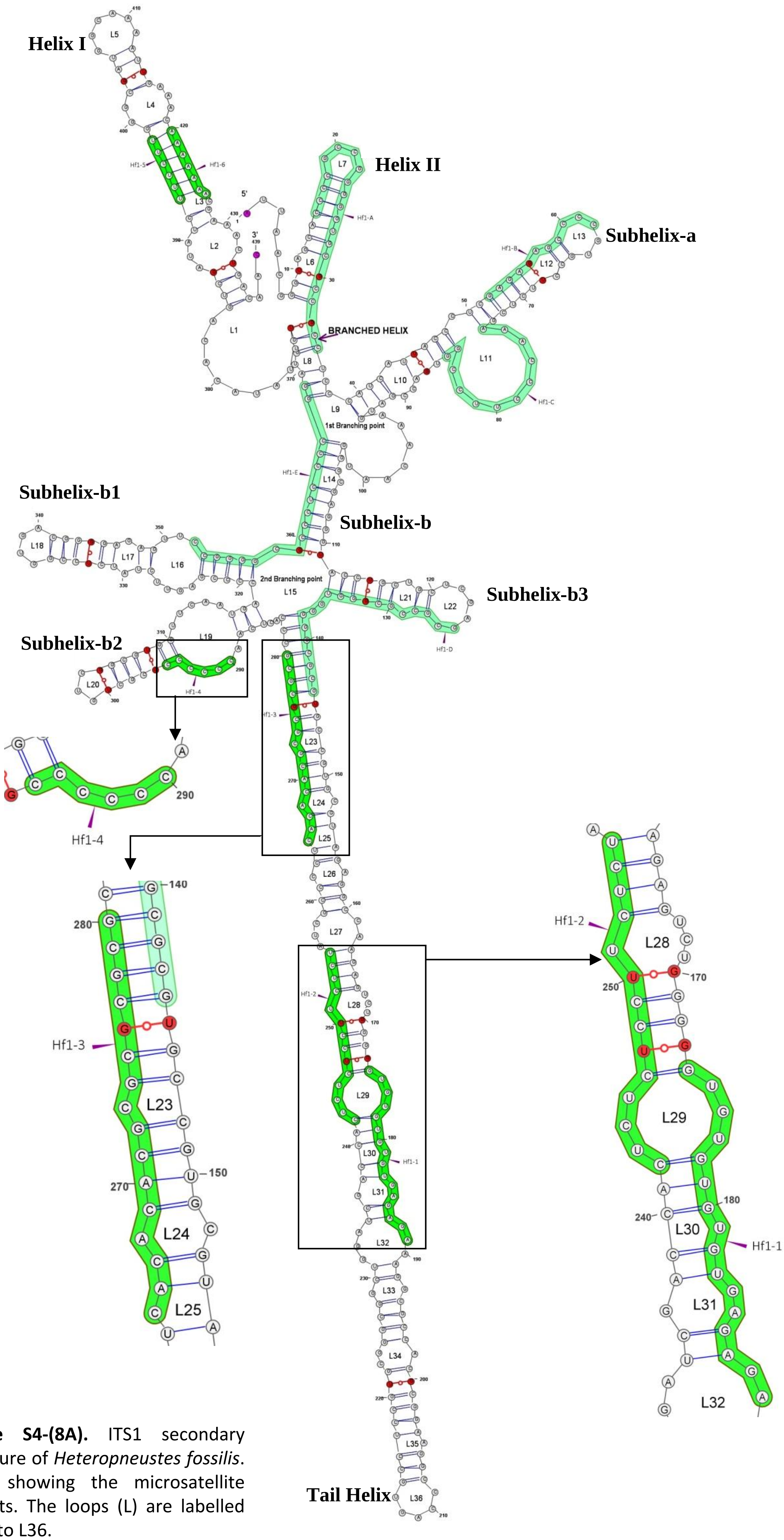


**Figure S4-(8A).** ITS1 secondary structure of *Heteropneustes fossilis*. Inset showing the microsatellite repeats. The loops (L) are labelled as L1 to L36.

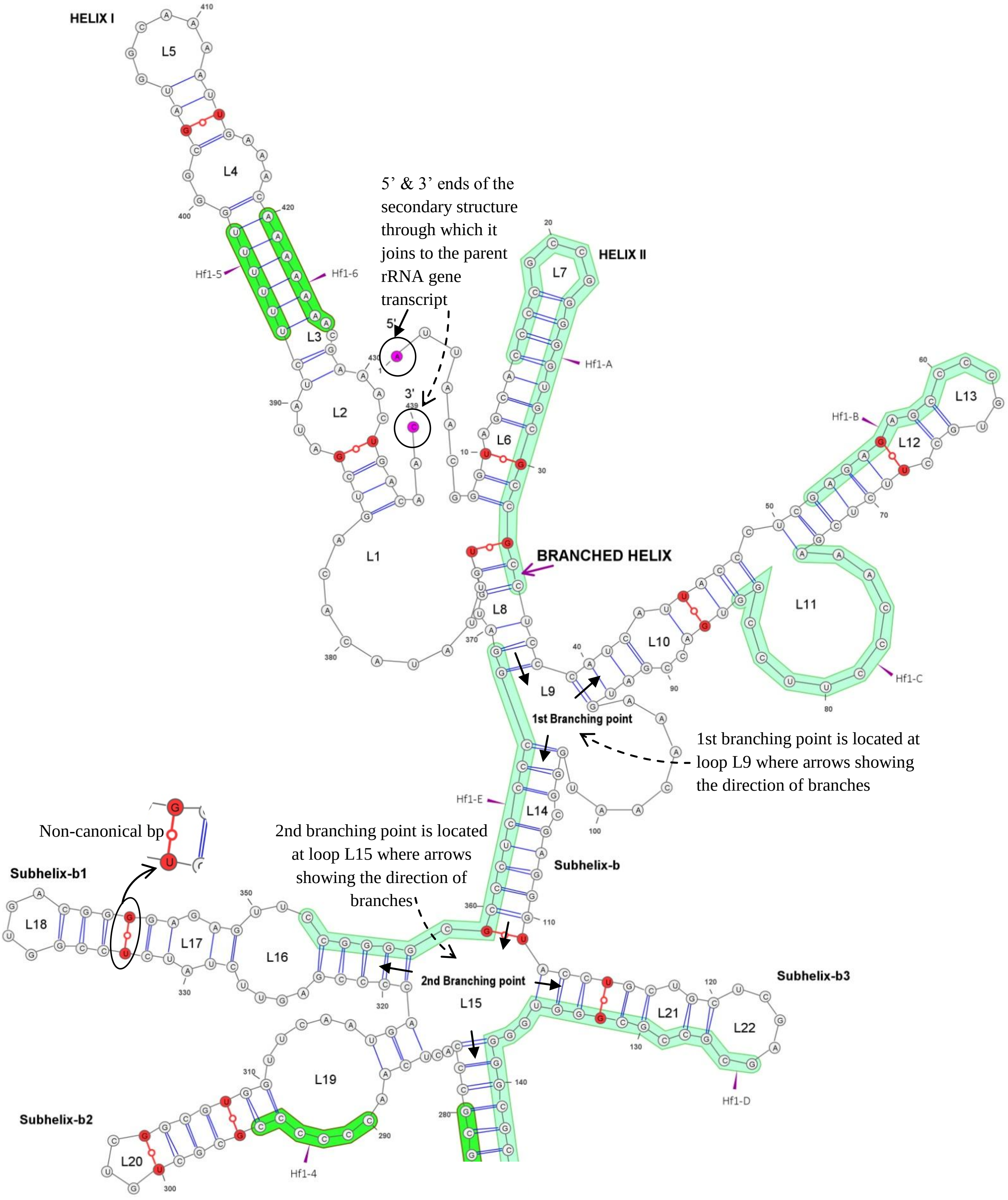


**Figure S4-(8B).** ITS1 secondary structure of *Heteropneustes fossilis*. Detailed view of upper branched region showing the branching pattern of helix and subhelices.

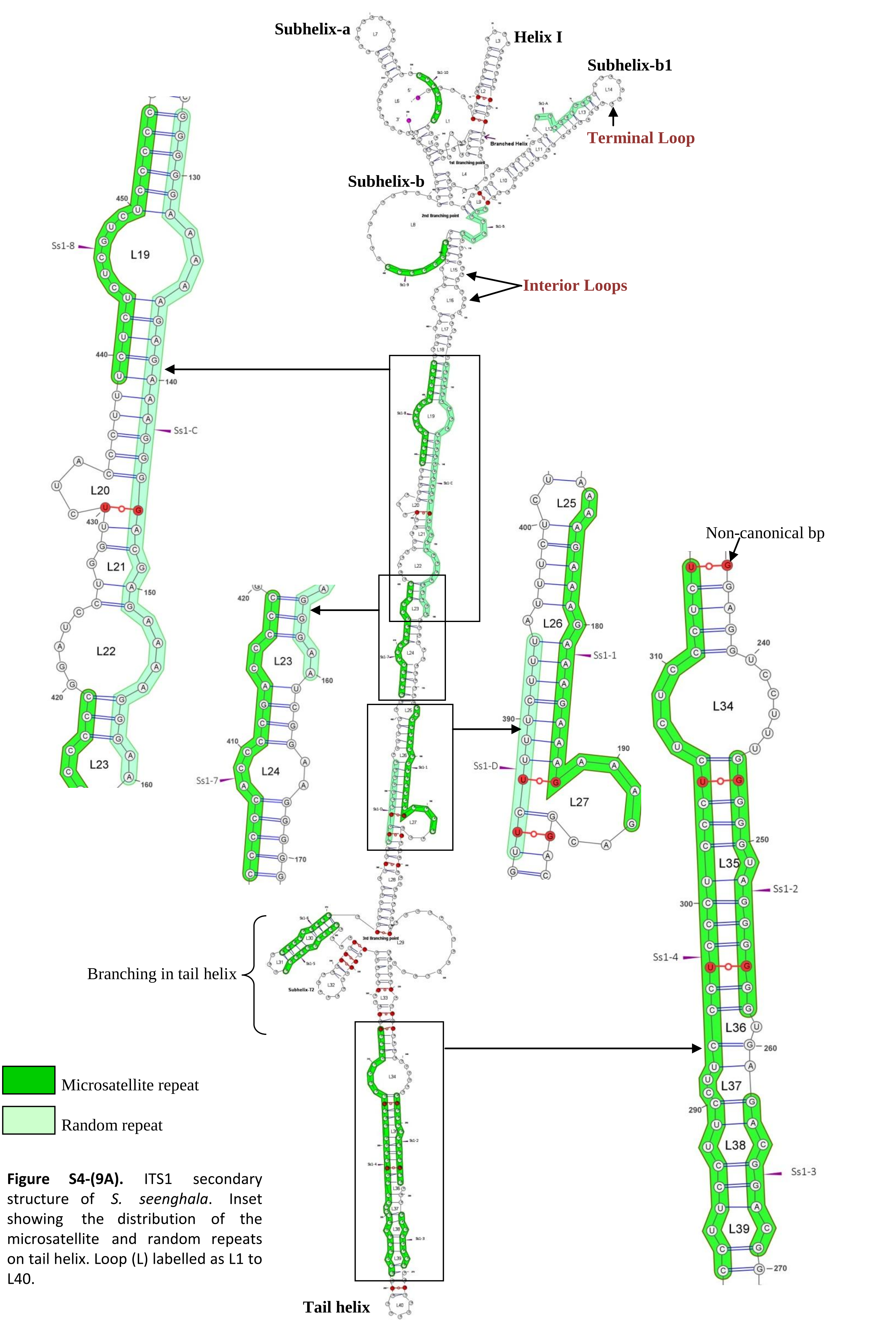


**Figure S4-(9A).** ITS1 secondary structure of *S. seenghala*. Inset showing the distribution of the microsatellite and random repeats on tail helix. Loop (L) labelled as L1 to L40.

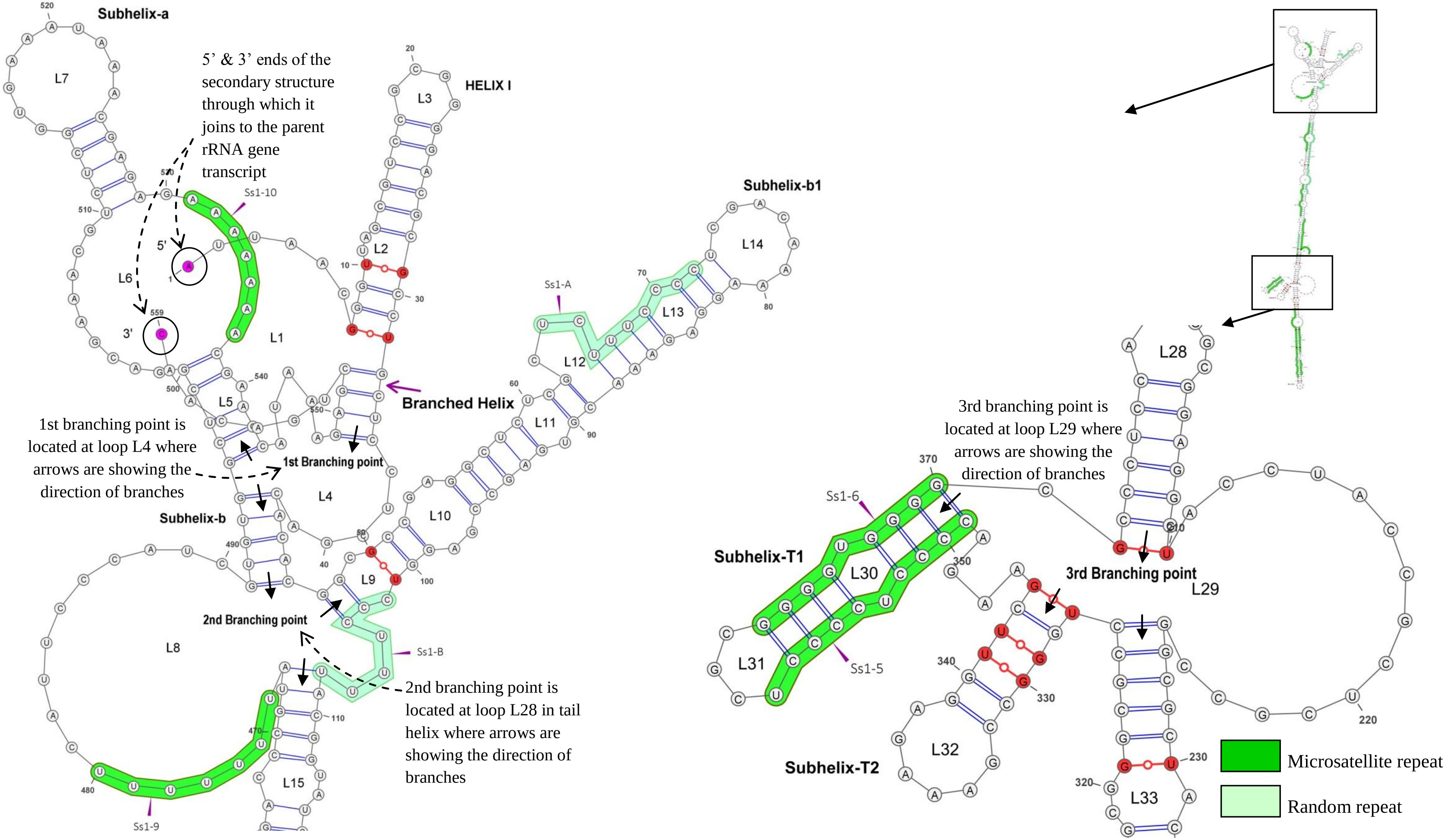


**Figure S4-(9B).** ITS1 secondary structure of *S. seenghala*. Inset showing the detailed view of upper branched region and branching on tail helix.

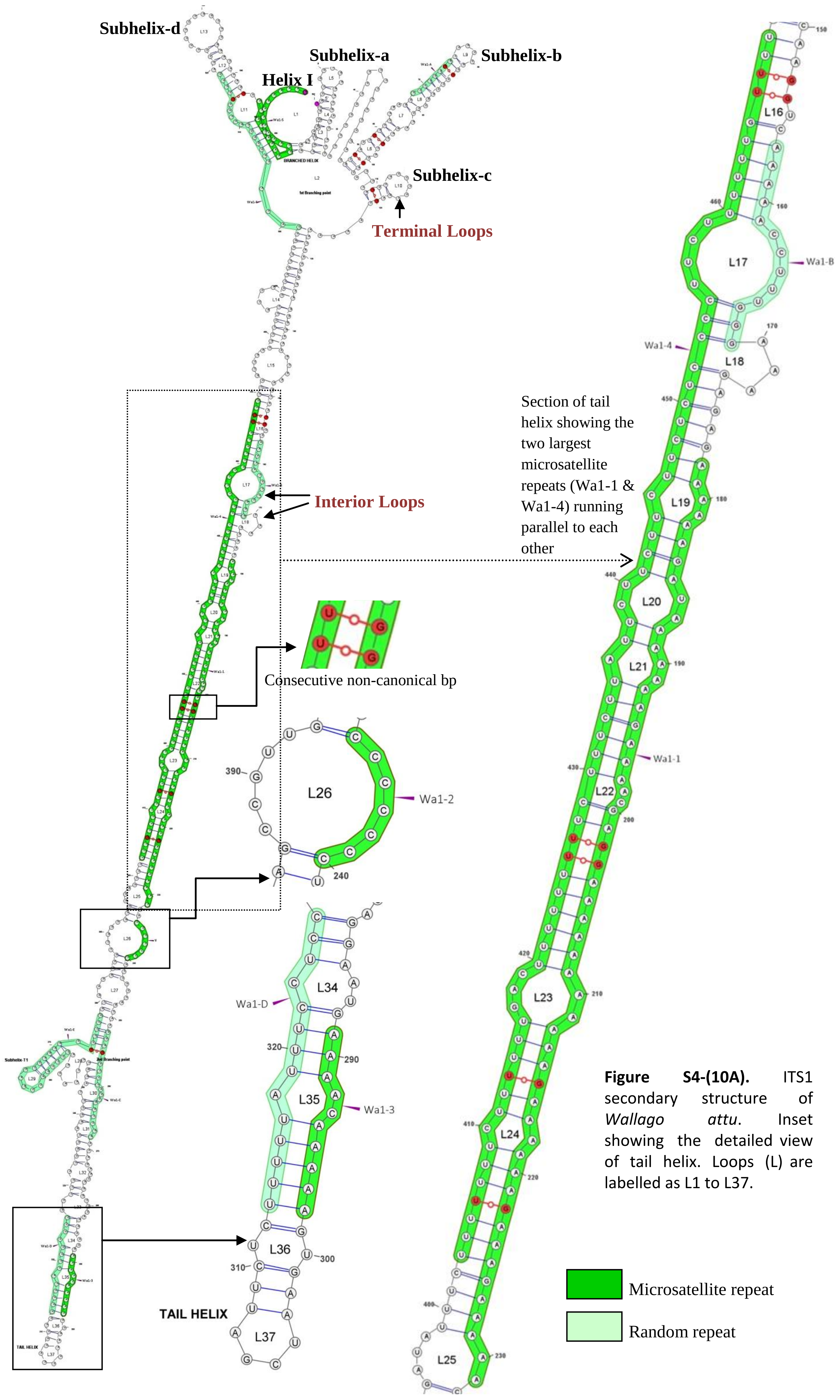


**Figure S4-(10A).** ITS1 secondary structure of *Wallago attu*. Inset showing the detailed view of tail helix. Loops (L) are labelled as L1 to L37.

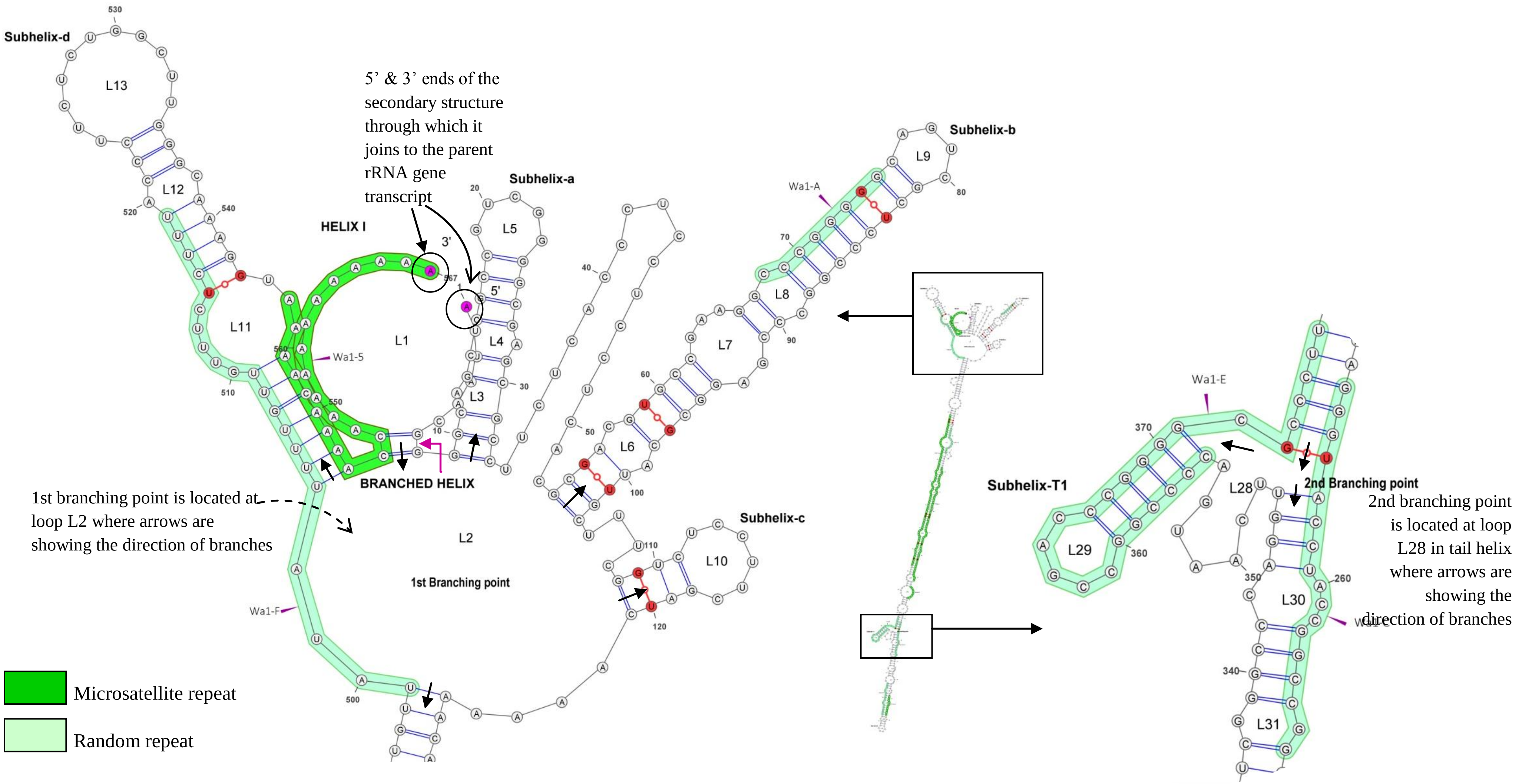


**Figure S4-(10B).** ITS1 secondary structure of *Wallago attu*. Inset showing the detailed view of upper branched region and branching on tail helix.
