## Supplementary material for "Exploring unique sequence repeat patterns and secondary structures in rDNA internal transcribed spacers ITS1 and ITS2 for characterization of catfish species": Supplementary tables.docx

**Supplementary Table S1:** Loop distribution comparison in the helices of ITS2 secondary structures of five catfish species

| **Species** | **Helix I** | **Helix II** | **Helix III** | **Helix IV** | **Total Loops** |
| --- | --- | --- | --- | --- | --- |
| *C. batrachus* | 4 | 9 | 9 | 3 | 25 |
| *C. gariepinus* | 2 | 5 | 24 | 16 | 47 |
| *S. seenghala* | 2 | 3 | 17 | 1 | 23 |
| *W. attu* | 1 | 6 | 25 | 15 | 47 |
| *H. fossilis* | 2 | 6 | 30 | 3 | 41 |

**Supplementary Table S2:** Non- canonical bp (G-U) comparison in the helices of ITS2 secondary structures of five catfish species

| **Species** | **Helix I** | **Helix II** | **Helix III** | **Helix IV** | **Total** |
| --- | --- | --- | --- | --- | --- |
| *C. batrachus* | 2 | 7 | 3 | 0 | 12 |
| *C. gariepinus* | 0 | 2 | 9 | 4 | 15 |
| *S. seenghala* | 2 | 0 | 5 | 0 | 07 |
| *W. attu* | 0 | 1 | 5 | 4 | 10 |
| *H. fossilis* | 0 | 1 | 4 | 0 | 05 |

**Supplementary Table S3:** Helix length comparison in the ITS2 secondary structures of catfish species.

| **Species** | **Helix I** | **Helix II** | **Helix III** | **Helix IV** |
| --- | --- | --- | --- | --- |
| *C. batrachus* | 33 | 105 | 114 | 42 |
| *C. gariepinus* | 14 | 60 | 288 | 137 |
| *S. seenghala* | 29 | 44 | 193 | 13 |
| *W. attu* | 06 | 49 | 226 | 100 |
| *H. fossilis* | 14 | 69 | 234 | 67 |

**Supplementary Table S4:** Tabular representation of ITS1 secondary structure. The table shows the variations in helix length, number of loops and non-canonical bp across the different helix and sub-helices of ITS1 secondary structure in *C. batrachus, C. gariepinus, S. seenghala, W. attu* and *H. fossilis*.

| **Species** | **Branched Helix** | | | | **Linear Helix** | | | |
| --- | --- | --- | --- | --- | --- | --- | --- | --- |
|  | **Branched Helix** | **length** | **No. of loop** | **Non-canonical bp** | **Linear Helix** | **length** | **No. of loop** | **Non-canonical bp** |
| ***C. batrachus*** | Branched helix | 259 | 29 | 14 | Helix I | 30 | 2 | 3 |
|  | Subhelix-a | 18 | 1 | 2 | Helix II | 38 | 3 | 3 |
|  | Subhelix-b | 53 | 4 | 2 | Helix III | 34 | 2 | - |
|  | Subhelix-c | 67 | 6 | 1 |  |  |  |  |
|  | Tail helix | 187 | 15 | 9 |  |  |  |  |
| ***C. gariepinus*** | Branched helix | 557 | 52 | 20 | Helix I | 18 | 1 | - |
|  | Subhelix-a | 20 | 2 | - |  |  |  |  |
|  | Subhelix-b | 65 | 6 | - |  |  |  |  |
|  | Subhelix-c | 32 | 2 | 2 |  |  |  |  |
|  | Subhelix-d1 | 57 | 5 | 3 |  |  |  |  |
|  | Tail helix | 314 | 32 | 14 |  |  |  |  |
| ***S. seenghala*** | Branched helix | 520 | 37 | 15 | Helix I | 26 | 2 | 2 |
|  | Subhelix-a | 48 | 3 | - |  |  |  |  |
|  | Subhelix-b | 454 | 32 | 15 |  |  |  |  |
|  | Subhelix-b1 | 58 | 6 | 1 |  |  |  |  |
|  | Tail helix | 366 | 26 | 14 |  |  |  |  |
|  | Subhelix-T1 | 22 | 2 | - |  |  |  |  |
|  | Subhelix-T2 | 18 | 1 | 3 |  |  |  |  |
| ***W. attu*** | Branched helix | 549 | 36 | 12 | Helix I | 04 | 1 | - |
|  | Subhelix-a | 35 | 3 | - |  |  |  |  |
|  | Subhelix-b | 51 | 4 | 3 |  |  |  |  |
|  | Subhelix-c | 14 | 1 | 1 |  |  |  |  |
|  | Subhelix-d | 51 | 3 | 1 |  |  |  |  |
|  | Tail helix | 374 | 24 | 7 |  |  |  |  |
|  | Subhelix-T1 | 17 | 1 | - |  |  |  |  |
| ***H. fossilis*** | Branched helix | 342 | 29 | 12 | Helix I | 53 | 4 | 2 |
|  | Subhelix-a | 56 | 4 | 2 | Helix II | 25 | 2 | 1 |
|  | Subhelix-b | 266 | 23 | 9 |  |  |  |  |
|  | Subhelix-b1 | 39 | 3 | 1 |  |  |  |  |
|  | Subhelix-b2 | 33 | 2 | 2 |  |  |  |  |
|  | Subhelix-b3 | 24 | 2 | 1 |  |  |  |  |
|  | Tail helix | 146 | 14 | 4 |  |  |  |  |
